# Identification of state-specific proteomic and transcriptomic signatures of microglia-derived extracellular vesicles

**DOI:** 10.1101/2023.07.28.551012

**Authors:** Juliet V. Santiago, Aditya Natu, Christina C. Ramelow, Sruti Rayaprolu, Hailian Xiao, Vishnu Kumar, Nicholas T. Seyfried, Srikant Rangaraju

## Abstract

Microglia are resident immune cells of the brain that play important roles in mediating inflammatory responses in several neurological diseases via direct and indirect mechanisms. One indirect mechanism may involve extracellular vesicle (EV) release, so that the molecular cargo transported by microglia-derived EVs can have functional effects by facilitating intercellular communication. The molecular composition of microglia-derived EVs, and how microglial activation states impacts EV composition and EV-mediated effects in neuroinflammation, remain poorly understood. We hypothesize that microglia-derived EVs have unique molecular profiles that are determined by microglial activation state. Using size-exclusion chromatography to purify EVs from BV2 microglia, combined with proteomic (label-free quantitative mass spectrometry or LFQ-MS) and transcriptomic (mRNA and non-coding RNA seq) methods, we obtained comprehensive molecular profiles of microglia-derived EVs. LFQ-MS identified several classic EV proteins (tetraspanins, ESCRT machinery, and heat shock proteins), in addition to over 200 proteins not previously reported in the literature. Unique mRNA and microRNA signatures of microglia-derived EVs were also identified. After treating BV2 microglia with lipopolysaccharide (LPS), interleukin-10, or transforming growth factor beta, to mimic pro-inflammatory, anti-inflammatory, or homeostatic states, respectively, LFQ-MS and RNA seq revealed novel state-specific proteomic and transcriptomic signatures of microglia-derived EVs. Particularly, LPS treatment had the most profound impact on proteomic and transcriptomic compositions of microglia-derived EVs. Furthermore, we found that EVs derived from LPS-activated microglia were able to induce pro-inflammatory transcriptomic changes in resting responder microglia, confirming the ability of microglia-derived EVs to relay functionally-relevant inflammatory signals. These comprehensive microglia-EV molecular datasets represent important resources for the neuroscience and glial communities, and provide novel insights into the role of microglia-derived EVs in neuroinflammation.

## Background

Microglia are the resident immune cells of the central nervous system (CNS). Given their major role in the innate immune system, the cell surface of microglia contains transporters, channels, and receptors for neurotransmitters, cytokines, and chemokines^1^. Microglia are constantly surveying their environment and can rapidly become activated to initiate immune responses^2^. Detection of immune activators and stimulating agents can cause microglia to alter their morphology and adopt heterogenous molecular profiles which can exert complex functions in different disease contexts. For example, microglia play an important role in neurodegenerative disease and neuroinflammation^3^. In the context of neurodegeneration, chronically activated microglia release inflammatory cytokines, such as tumor necrosis factor (TNF), IL-6, and reactive oxygen species. Furthermore, to detect immune activators, microglia are equipped with toll-like receptors (TLRs). Activation of TLRs on microglia can lead to the production of pro-inflammatory cytokines and release of chemokines^4^. Comprehensive single-cell RNA sequencing analyses have revealed novel microglia types associated with neurodegenerative diseases, also referred to as disease associated microglia (DAM)^5^. This subset of microglia displays unique transcriptional features, expressing typical microglial protein markers, such as Iba1 and Hexb but with a decrease in signature microglia homeostatic genes, such as *Cx3cr1* and *P2ry12/13*, and upregulation of genes involved in lipid metabolism and phagocytic pathways, such *Apoe, Lpl, CD9, Cst7,* and *Trem2*. Furthermore, DAM can play dual roles in Alzheimer’s disease (AD) pathogenesis, involving the production and release of pro-inflammatory factors including cytokines and toxic factors as well as neuroprotective, anti-inflammatory functions^6^. While distinguishing the function of microglia as either “protective” or “detrimental” is difficult, there is accumulating evidence that microglia can dynamically switch phenotypes in response to stimuli. However, further investigation is needed to fully understand the involvement of different microglia states in CNS diseases.

A newly identified mechanism of microglia-mediated neuroinflammatory responses in neurodegenerative disease involves secretion of extracellular vesicles (EVs)^7^. EVs are lipid bound vesicles that are composed of lipids, proteins, metabolites, and nucleic acids. EVs can be classified as microvesicles, apoptotic bodies, or exosomes^8^. EV subtype is determined by biogenesis, release pathway, size, density, function, and cargo. However, due to significant overlap in protein profiles and size, along with difficulty in proving the origin of EVs, it is often preferred to refer to exosomes as EVs within a particular size range^9, 10^. Exosomes are small EVs (30-150 nm) of endocytic origin and are secreted by almost all cell types. Exosomes transport specific cargo such as proteins, messenger RNAs (mRNAs), and microRNAs (miRNAs) between cells to facilitate intercellular communication and influence downstream signaling events^11^. Exosomes can interact with recipient cells through endocytosis, fusion with the plasma membrane, or ligand receptor interaction^12^. Accumulating studies have demonstrated that microglia-derived exosomes can serve as key mediators in pathologies associated with neurodegenerative diseases. However, the molecular compositions of microglia-derived small EVs remain poorly understood.

Classical hallmarks of AD are abnormal aggregation of amyloid beta (Aβ) proteins into plaques and tau misfolding^13^. Microglia-derived exosomes in tau models of AD, release and spread pathologic tau between neurons. Recent evidence suggests that depleting microglia and inhibiting the synthesis of exosomes significantly suppresses pathologic tau propagation^14^. This study indicates the role microglia-derived exosomes may have in the pathological spread of tau. Verderio et al. found that microvesicles and exosomes derived from LPS-preactivated cultured microglial cells, induced a dose-dependent activation of resting astrocytes and microglia^15^. The findings of this study suggest that the cargo from EVs can transfer an inflammatory signal to recipient cells, thus exacerbating neuroinflammatory conditions.

Proteomic analysis of exosomes derived from the N9 microglial cell line identified the proteomic composition of microglia-derived exosomes from cell culture medium^16^. Several hallmark EV proteins previously identified from EVs derived from cell types were found in the N9-derived EV proteome; such as, cytoskeletal proteins, heat shock proteins, integrins, and tetraspanin proteins. Of particular interest, was the aminopeptidase N (CD13) found in microglial exosomal proteins but not in exosomes derived from B cells and DC cells. Functional assays looking at aminopeptidase activity revealed that microglia exosomal CD13 is active in neuropeptide degradation.

Yang et al. characterized microglial EV protein cargo following lipopolysaccharide (LPS) and TNF inhibitor treatment^17^. Following 12-hour LPS stimulation of BV2 microglia, EVs were found to have significantly higher levels of pro-inflammatory cytokines TNF and interleukin (IL-10), seen by ELISA. Furthermore, inhibition of the TNF signaling pathway resulted in a reduction of EVs released from LPS activated microglia. Mass spectrometry (MS)-based experiments identified 49 unique proteins in EVs derived from LPS activated microglia compared to control, with a majority of the proteins associated with transcription and translation. From this study, it can be inferred that microglia respond to an LPS challenge by releasing EVs with unique cargoes that may be implicated in inflammatory mechanisms. The data from both these studies suggest that microglia derived EVs have distinct proteomic profiles which may play a role in inflammatory responses in neurodegenerative diseases. However, low-depth of proteomic coverage remains a limitation of these studies.

Additionally, accumulating evidence suggests that EV miRNAs have the potential to influence disease pathogenesis and treatment outcomes. MiRNAs are small non-coding RNAs that regulate the expression of specific gene targets^18^. Release of miRNAs from EVs can influence target cell function^19^. Huang et al. found that increased miR-124-3p expression in microglia-derived exosomes after traumatic brain injury can inhibit neuronal inflammation and contribute to neurite outgrowth in recipient neurons^20^. Even though miRNAs are more abundant in EVs than larger species of RNA, mRNAs have also been found to be loaded into EVs and transferred to recipient cells^21, 22^. Ratajcazk et al. provided the first evidence that mRNAs transferred to recipient cells are functional and can be translated into proteins, leading to biological changes in recipient cells^23^. Overall, several studies have highlighted that the activation of microglia can have both a direct or indirect influence on neuroinflammation and disease progression. The uptake of EVs from recipient cells appears to mediate the indirect mechanisms by which microglia can affect disease progression. However, there is a lack of proteomic and transcriptomic characterization of EVs from distinct microglia states. Identification of the proteomic and transcriptomic cargo in EVs from distinct microglia states may elucidate key targets and pathways that are involved in EV mediated neuroinflammation.

In this study, we have generated comprehensive datasets of proteomics and transcriptomics from BV2 microglia cells and BV2-derived EVs under resting states as well as following inflammatory challenge with either LPS, IL-10, or transforming growth factor β (TGF-β)^24–26^, which have been documented to induce distinct molecular phenotypes in vitro. Using label-free quantitative mass spectrometry (LFQ-MS) and RNA sequencing of mRNA and miRNA species, we identified unique molecular signatures of EVs at the proteomic and transcriptomic levels, including several features not previously described. We also identified unique state-dependent molecular characteristics of microglia-derived EVs, in which LPS-effects were most predominant at the level of proteins, mRNA and miRNA. Next, we asked whether EV cargo from distinct microglia states can impact the gene expression profile of resting (responder) BV2 cells. To address this, we isolated EVs from the cell culture media (CCM) of four BV2 cell conditions (control, LPS, IL-10, TGF-β) and then equally dosed individual wells of BV2 cells with those EVs to assess gene expression changes in recipient microglia using RNA sequencing. Taken together, our study provides several comprehensive molecular datasets on microglia-derived EVs and provides novel insights into the role of microglia-derived EVs in neuroinflammation, including increased packaging of mRNAs into EVs following inflammatory activation.

## Results

### Verification of EV purification from BV2 cell culture medium

We applied size exclusion chromatography (Izon qEV) based methods to isolate EVs from BV2 microglia culture supernatants. Prior to MS and RNA seq studies of BV2 microglia-derived EVs, we performed several quality control studies to confirm successful enrichment of EVs from cell culture media. We characterized EV morphology, size, concentration, and classical markers using transmission electron microscopy (TEM), immunogold TEM, nanoparticle tracking analysis (NTA), and western blot analyses (**Figure 1A**). To characterize the morphology of BV2-derived EVs, isolated EVs were examined using high resolution TEM following negative staining. TEM revealed vesicles with consistent round and cup-shaped morphology within ∼50-150nm size range, as would be expected for EVs (**Figure 1B**). Immunogold-labeled TEM studies using 6nm gold particles confirmed CD9 labeling on the surface of EVs (**Figure 1C**). NTA of several EV preparations also verified that our isolated EVs are typically within the range 50-200nm in diameter (mean particle size =87.9 +/- 6.1 nm, mode particle size = 44.2 +/- 9.0 nm) (**Figure 1D**). Lastly, we performed western blot analyses of cell lysates and EVs from BV2 microglia and found that canonical EV protein markers CD9 and TSG101 were enriched in EV lysates, in contrast with enrichment of Calnexin in BV2 cell lysates (**Figure 1E**). These results using complimentary validation methods as recommended by the International Society for Extracellular Vesicles^27^, confirm the validity of our EV isolation approach and its suitability for subsequent molecular characterization studies.

**Figure 1:**
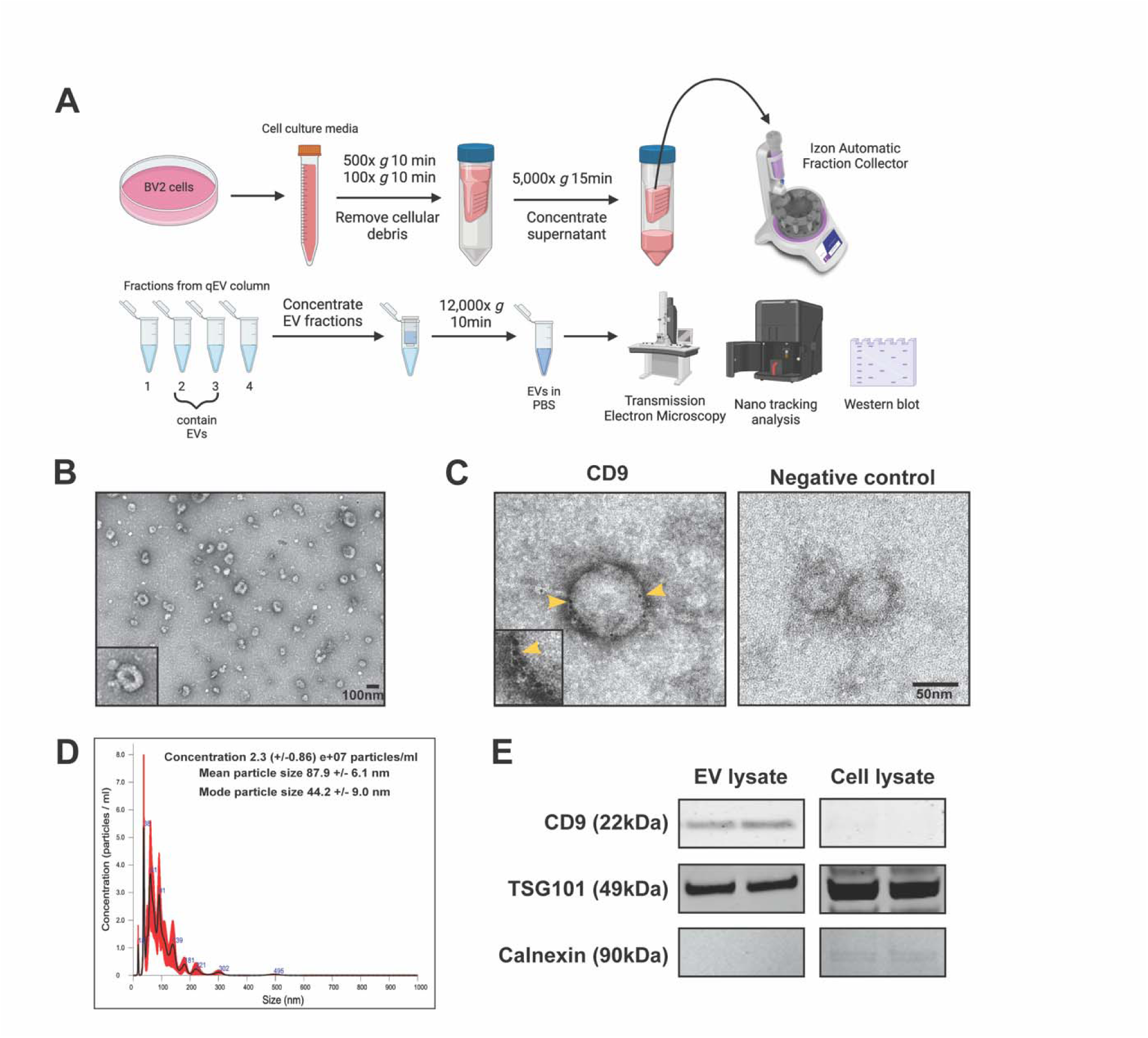
Purification of BV2 microglia-derived extracellular vesicles (EVs). **A.** Illustration outlining the process for isolating EVs from cell culture media. Created with BioRender. **B.** Transmission electron microscopy (TEM) image of isolated EVs from BV2 cells. Scale bar is 100nm. **C.** TEM immunogold labeling of EVs (6nm gold particles) shows CD9 protein on the surface of EVs. CD9 is a canonical EV surface marker. **D.** Nanoparticle tracking analysis (NTA) shows size and concentration of isolated EVs from BV2 cell culture media. **E.** Western blot analyses of EVs and cell lysate, probed with antibodies to detect EV-enriched and cytosolic markers. CD9 and TSG101 are positive EV markers and Calnexin is a negative EV marker. Two biologically-independent experimental samples were blotted for each condition.

### Identification of proteomic signatures of microglia-derived EVs by quantitative MS

We performed LFQ-MS on 16 cell lysates and 16 EV lysates derived from BV2 microglia. These included four groups of BV2 cells (n=4/group) that were treated (72 hours) with either LPS (100 ng/mL) to polarize to a pro-inflammatory state, IL-10 (50 ng/mL) to polarize to a protective state, TGF-β (50 ng/mL) to polarize to a homeostatic state or untreated controls (**Figure 3A**). LFQ-MS identified 533 proteins in the EV proteome and 1,882 proteins in BV2 cell proteome, across all conditions (**Additional file 1 and Additional file 2**). To contrast the EV proteome with the whole cell proteome, we first compared BV2 EVs (n=533 proteins) with BV2 cell lysate proteomes (n=1,882 proteins). The effect of treatment was not considered when comparing all EVs to all cell lysates. Forty-six proteins were differently enriched in EVs while 1,178 proteins were differentially enriched in cell lysates (**Figure 2A, Additional file 3**). Canonical exosome/EV related proteins expected to be present in all EVs independent of cell type of origin (including SDCBP, IGSF8 and three tetraspanins namely CD9, CD81, and CD63) were significantly enriched in the EV proteome compared to the whole cell proteome (**Figure 2A).** In contrast, endoplasmic reticulum proteins expected to be present in cell lysates but not in EVs, namely SSR1 and Calnexin, were highly enriched in the cell proteome but not the EV proteome. Gene set enrichment (GSEA) of EV-abundant proteins showed enrichment of extracellular region (APOE, SAA), membrane organization (CD9, ITGB1, MFGE8), and localization (RAB6B, SLC1A5, SLC38A2) terms while cytosolic and intracellular organelle terms were enriched in the cellular proteomes (**Figure 2B**).

**Figure 2:**
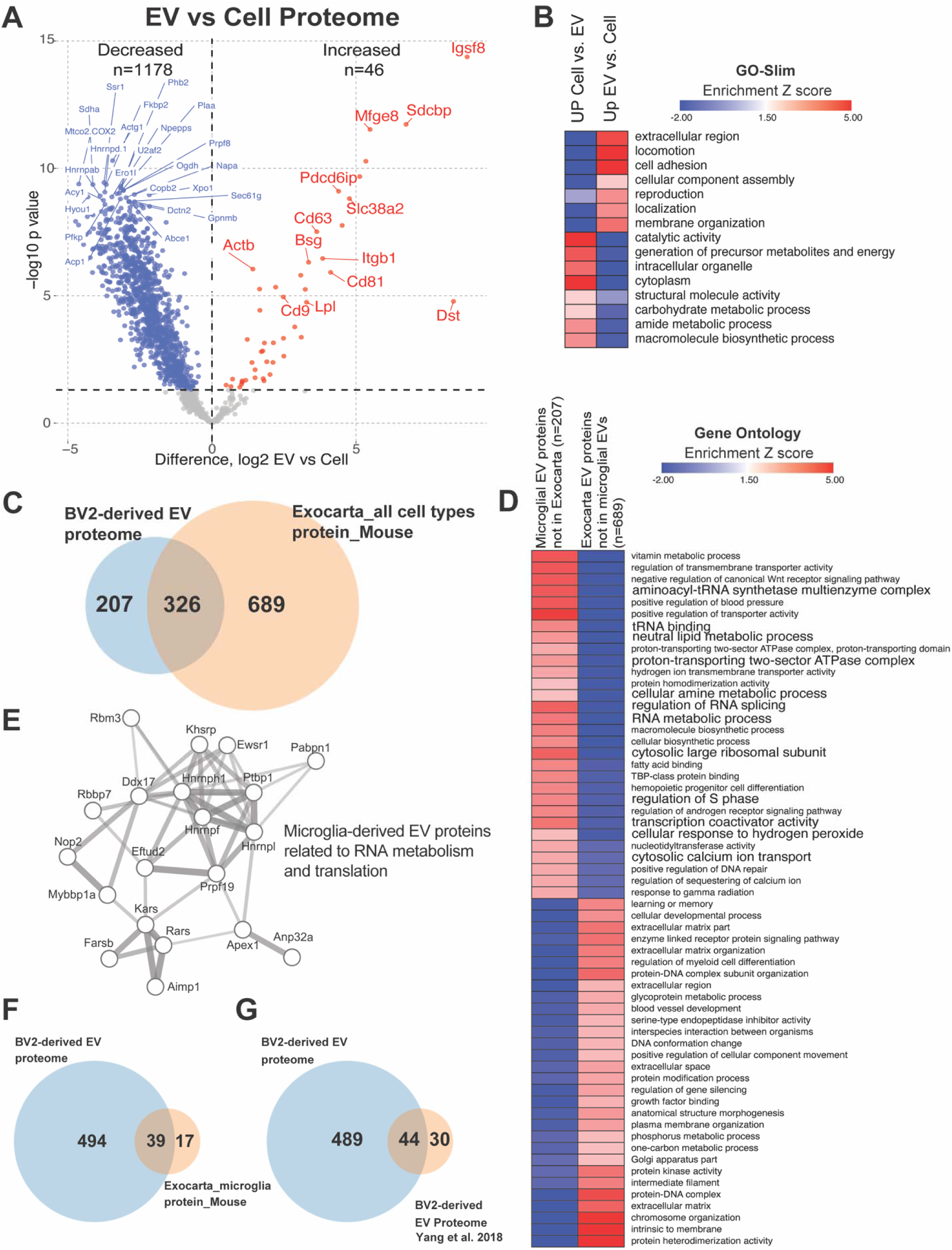
Comparison of BV2 whole-cell proteome and BV2-derived EV proteome. **A.** Enrichment of EV related proteins in EV proteome compared to BV2 cell proteome. Red labeled circles are in the Exocarta top 100 list. **B.** Heatmap representation, based on enrichment Z-scores, of GOSlim for list of differentially enriched proteins (DEPs) from Figure 2A. **C.** Comparison of 533 identified proteins from BV2-derived extracellular vesicles across all conditions to online database Exocarta mouse, all cell types, protein. **D.** Heatmap representation, based on enrichment Z-scores, of Gene Ontology for list of 207 proteins and list of 689 proteins from Figure 2B. **E.** STRING protein-protein-interaction network highlighting protein groups related to RNA metabolism and protein translation, that are unique to BV2 microglia-derived EVs as compared to other cell type-derived EVs. Edges represent protein-protein associations. Line thickness represents edge confidence. **F.** Comparison of 533 identified proteins from BV2 derived extracellular vesicles across all conditions to online database Exocarta mouse, microglia, protein. **G.** Comparison of 533 identified proteins from BV2 derived extracellular vesicles across all conditions to Yang et al. 2018 BV2-derived EV proteome.

**Figure 3:**
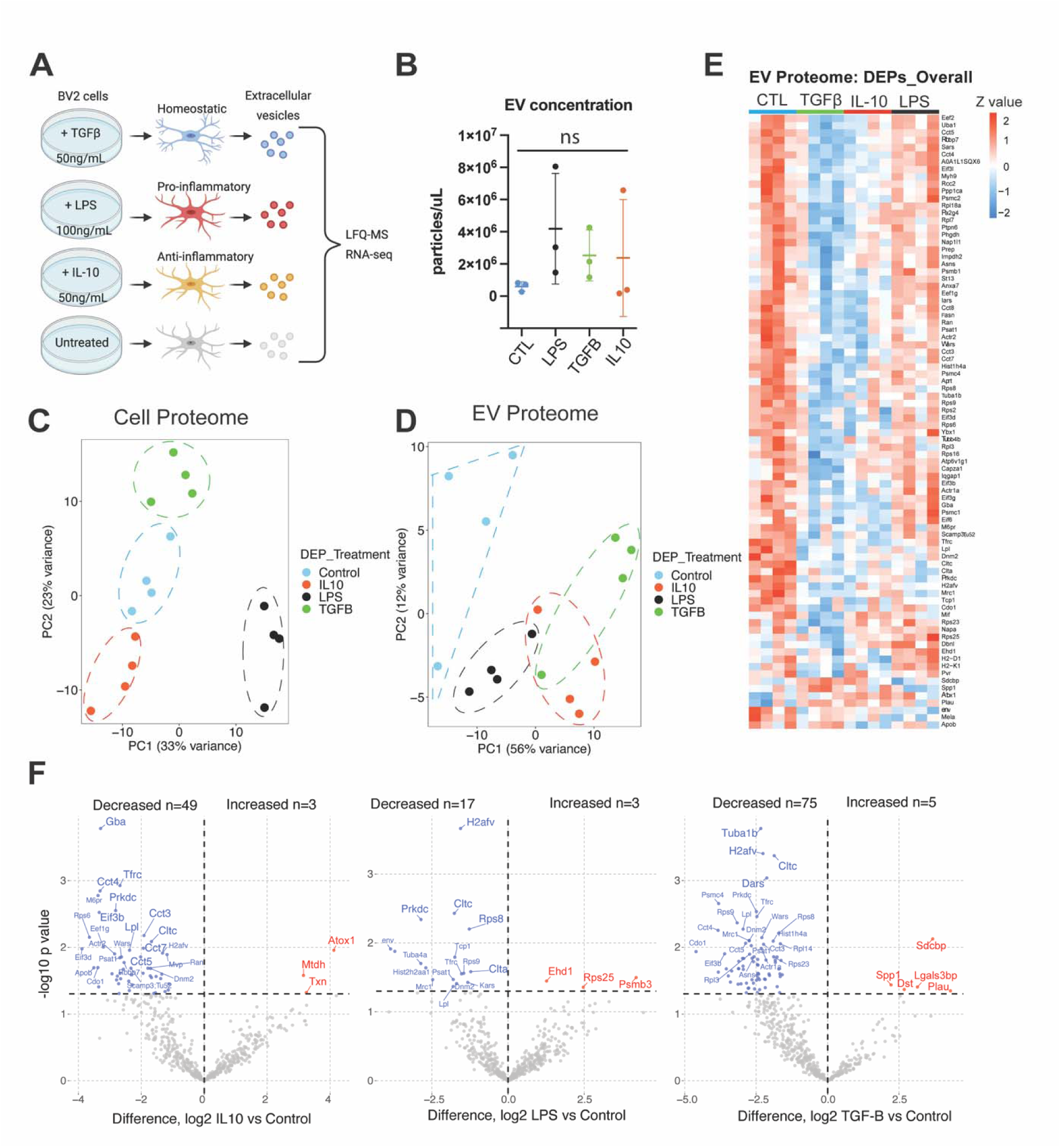
BV2 cells of distinct states and their EVs have unique protein signatures. **A.** llustration outlining the experimental setup. Created with BioRender. **B.** Comparison of EV yield from BV2 cell culture media, across different treatment conditions. One-way ANOVA, p-value is not significant (p=0.4684). **C.** Principal component analysis (PCA) plot of BV2 cell proteome – p<0.05, one-way ANOVA (n=333 proteins). **D.** PCA plot of BV2-derived EV proteome – p<0.05, one-way ANOVA (n=84 proteins)**. E.** Heatmap of DEPs – p<0.05, one-way ANOVA (n=84 proteins). **F.** Volcano plots showing DEPs in EV proteome comparing each treatment group to the control group (p<0.05).

**Figure 4:**
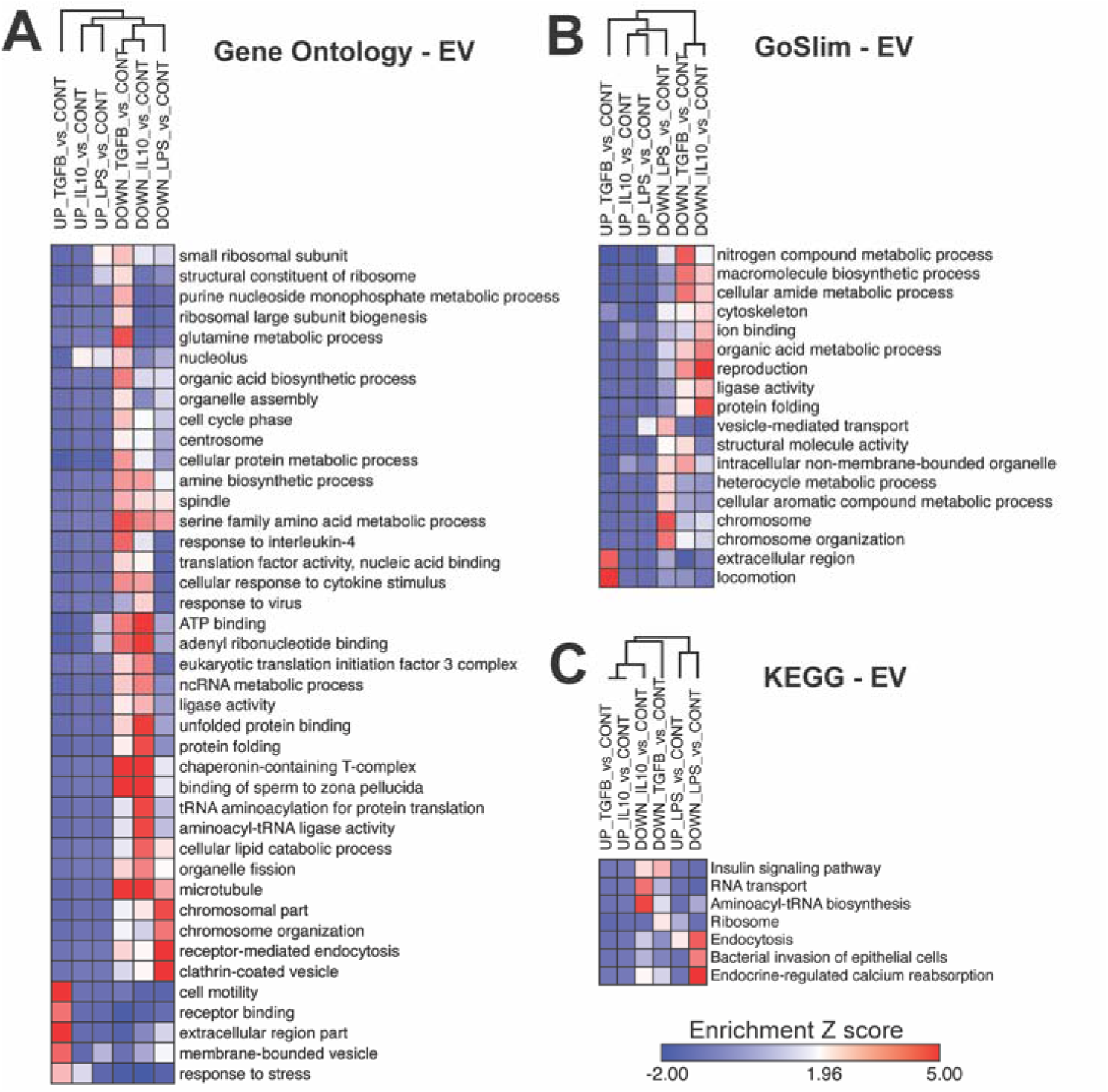
Gene ontology (GO) analysis of identified proteins in EVs from distinct microglia states. **A.** Heatmap representation, based on enrichment Z-scores, of Gene Ontology for EVs from polarized BV2 cells (p<0.05). **B.** Heatmap representation, based on enrichment Z-scores, of GO Slim for EVs from polarized BV2 cells (p<0.05). **C.** Heatmap representation, based on enrichment Z-scores, of KEGG for EVs from polarized BV2 cells (p<0.05).

Next, we compared our BV2 microglia-derived EV proteome to existing lists of proteins previously detected in proteomic studies of EVs from any mouse cell types (1,015 EV proteins in ExoCarta) (**Figure 2C**), the Exocarta protein list for mouse microglia (**Figure 2F**, 56 proteins), and the Yang et al. 2018 BV2-derived EV proteome (**Figure 2G**, 74 proteins) (Additional file 4). Our BV2-derived EV proteome (533 proteins) shared 326 proteins with the ExoCarta protein list of 1,015 EV proteins detected across all mouse cell types (207 novel proteins), 39 proteins with the Exocarta protein list of 56 mouse microglia (494 novel proteins) and 44 proteins with Yang et al. 2018 BV2-derived EV proteome that included 74 proteins (489 novel proteins). We also performed GSEA (**Additional file 4**) on three groups of proteins based on these analyses (207 novel BV2 microglial EV proteins not reported in other EV proteomes, 689 proteins reported in non-microglial EV proteomes but not in our data, and 326 proteins common to both microglial and non-microglial EV proteomes). Proteins unique to BV2 microglia-derived EVs were enriched in ontologies related to positive regulation of transporter activity, regulation of RNA splicing, cytosolic large ribosomal subunit, and aminoacyl-tRNA synthetase complex (**Figure 2D**). On the other hand, the 689 proteins not captured in our BV2-derived EV proteome showed enrichment of proteins related to protein heterodimerization activity, chromosome organization, and protein kinase activity (**Figure 2D**). We also identified networks of known direct (protein-protein) and indirect (functional) interactions (STRING) within these core protein signatures related to RNA metabolism and protein translation, that are unique to our BV2 microglia-derived EVs as compared to other cell type-derived EVs (**Figure 2E)**.

Taken together, these analyses verify the validity of our EV proteomes by confirming enrichment abundances of canonical EV markers, highlight the increased depth of our EV proteome as compared to prior microglial studies, and identify novel proteomic characteristics of microglia-derived EVs that are distinct from non-microglial EVs which may have functional implications.

### Microglial activation state impacts proteomic characteristics of EVs

Three groups of BV2 cells (n=4/group) were treated with either LPS (100 ng/mL) to polarize to a pro-inflammatory state, IL-10 (50 ng/mL) to polarize to a protective state, or TGF-β (50 ng/mL) to polarize to a homeostatic state (**Figure 3A**). Untreated BV2 cells served as a control group. To ensure that any state-related differences in EV proteomes are not related to impacts of EV yield, we used NTA to compare EV concentrations across treatment conditions and observed no significant differences across treatment groups (**Figure 3B**). To confirm that in vitro conditions induced distinct microglial activation states, we analyzed BV2 cell LFQ-MS proteomes and 334 differentially enriched proteins (DEPs) across polarization states, as compared to control (one-way ANOVA FDR-adjusted p<0.05 & log_2_FC>0) (**Supplementary Figure 1, Additional file 1**). Principal component analysis (PCA) using these DEPs showed that 56% of variance was explained by 2 principal components (PCs), of which PC1 explained 33% of variance while PC2 explained 23% of variance. Notably, the PCA identified four distinct proteomic clusters (Figure 3B). As compared to untreated control BV2 microglia, LPS treatment increased levels of several pro-inflammatory proteins (e.g., OAS1L, IRG1, ACSI1) (**Supplementary Figure 1A**). For the BV2 cell proteome, GSEA using the generated up and down lists for each treatment against the background list (1,882 proteins) showed unique enrichment patterns across treatment groups (**Supplementary Figure 1B**). For example, LPS treated cells demonstrated an enrichment in proteins related to antigen processing, defense response, and immune response, whereas there was an observed decreased abundance in proteins lipid binding and phospholipase inhibitor activity. These results agree with prior proteomic studies of BV2 microglia using similar in vitro conditions, thereby confirming successful polarization of BV2 microglia to distinct molecular states^28, 29^. We also found that TGF-β treated cells demonstrated an enrichment in proteins related to positive regulation of homeostatic regulation and gluconeogenesis following, while IL-10 treated cells showed enrichment of proteins related to lipid binding and lipase inhibitor activity.

We next examined BV2-derived EV proteomes to determine whether microglial polarization state impacts proteomic composition of EVs. To account for unequal protein loading per sample, we performed our analyses of EV proteomes after normalizing MS data to total protein content in each sample, based on sum intensities across all quantified proteins. Our analyses of EV proteomes also accounted for batch effects since 4 different batches of experiments were performed, each with 1 replicate per condition (**Supplementary Figure 2**). We identified 83 DEPs across microglia states compared to control (p<0.05 & log_2_FC>0) (**Additional file 2**). PCA based on these DEPs identified four clusters among EVs (**Figure 3C**). Interestingly, all polarized BV2 microglia-derived EVs clustered away from control BV2-derived EVs and majority of these DEPs showed lower levels in polarized conditions as compared to the control group. Distinct signatures across polarization states compared to control are shown in **Figure 3D and 3E**.

We performed GSEA based on the DEPs for each treatment (compared to the control group), against the background list of 533 proteins identified in the EV proteome. We identified “Biological processes” and “Molecular function” terms along with KEGG pathways (**Figure A-C**) enriched in EV signatures from distinct BV2 microglial polarization states. GSEA of increased DEPs were limited due to small numbers of proteins that showed increased levels. EVs derived from BV2 cells treated with LPS demonstrated an enrichment in proteins related to proteasome activity (e.g., PSMB3) and reduction in proteins related to chromosomal organization (e.g., CLTC, HIST2H2AA1,TCP1), and clathrin/receptor mediated endocytosis (e.g., CLTC, DNM2). Conversely, EVs derived from BV2 cells treated with IL-10 and TGFβ demonstrated a reduction in cytosolic chaperonin Cct ring complex proteins (e.g., CCT3, CCT4), microtubule related proteins and ATP/nucleotide binding (e.g., CCT4, TCP1, TUBA1B). EVs derived from BV2 cells treated with TGFβ showed enrichment of proteins related to membrane bound vesicle and receptor binding (CD9 AND SDCBP). These proteomic analyses of EVs confirm that microglial activation states can determine the proteomic compositions of EVs.

### Novel transcriptomic signatures of microglia-derived EVs in resting and pro-inflammatory states

Beyond proteins, EVs carry mRNA and miRNA cargo which may be important mediators of microglia-mediated mechanisms of neuroinflammation. It is also possible that microglia state-dependent effects on EVs are more likely to be more pronounced at the levels of mRNA and miRNA levels. Therefore, we performed RNA seq to quantify mRNA and miRNA compositions of EVs and their corresponding cell lysates, after polarizing stimuli. Three groups of BV2 cells (n=3/group) were treated with either LPS (100 ng/mL), IL-10 (50 ng/mL) or TGF-β (50 ng/mL) to mimic conditions used for proteomic studies. Untreated BV2 cells served as a control group. Following 72 hours of treatment, whole cells and EVs were isolated from cell culture media, lysed in Trizol, followed by RNA extraction, purification and quality control steps. To assess length of RNA from BV2 cells and their EVs, RNA was characterized through capillary electrophoresis using two separate analyses kits: Pico and small RNA kits for bioanalyzer. These analyses (Additional file 5) revealed that cells display peaks at the ∼2,000 nt (18s) and ∼4,000 nt (28s), while EVs displayed peaks predominately between ∼20-100 nt which correspond to small RNAs. To characterize the different subtypes of RNAs in EV and whole cells, we proceeded with mRNA sequencing and small RNA sequencing. We first discuss our mRNA sequencing results.

Sequencing of mRNAs identified over 12,000 mRNA species (transcripts) in EVs and cells across all conditions. As we observed at the proteomic level, different polarizing conditions induced distinct transcriptomic states in BV2 microglial cells, as evidenced by PCA in which PC1 explained 73% of variance while PC2 explained 19% of variance (**Figure 5A**). Similarly, PCA revealed a large effect of LPS on EV transcriptomic composition as compared to other polarizing stimuli (**Figure 5B**). We first assessed unique transcriptomic signatures of EVs as compared to whole cells (control BV2 microglia only) (**Figure 5C**). EVs contain higher levels of over 1,000 mRNA species, including *Pak7, Arhgef40, and Saa3*. In contrast, cellular transcriptomes had higher expression of over 1,500 mRNAs including *Cebpb, Prkaca, and Pip5k1c*. Interestingly, *Cebpb* is a transcription factor known to regulate immune genes including genes critical for microglia switch from homeostatic to DAM^30^. GSEA of highly-enriched mRNAs (>=2-fold change in EVs or whole cells) showed enrichment of translation, ribosome, and cytokine activity in EVs, while we observed enrichment of terms such as ion binding and homeostasis and transmembrane receptor activity (**Figure 5D, Additional file 6**).

**Figure 5:**
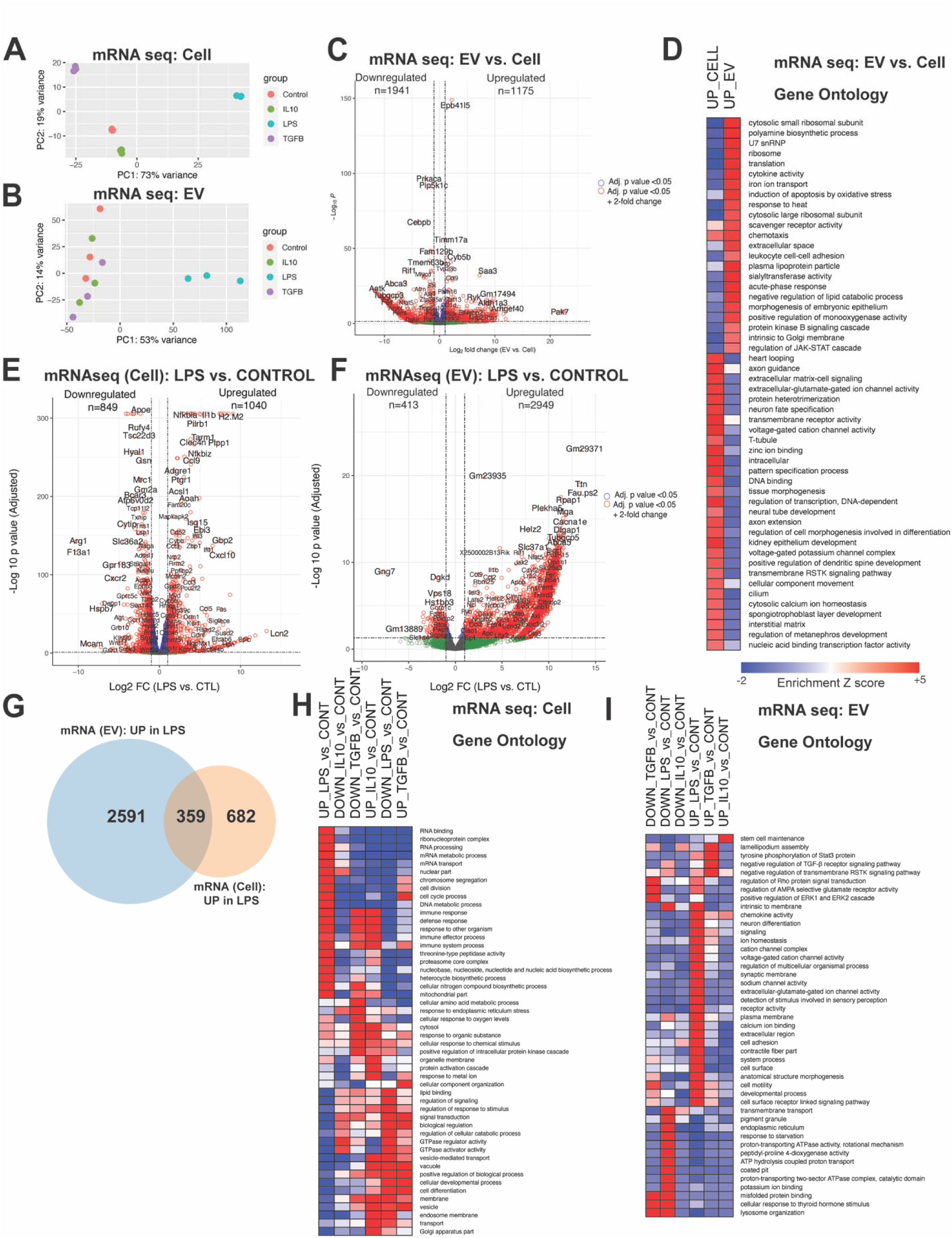
BV2 cells of distinct states and their EVs have unique mRNA signatures. **A.** PCA plot on mRNA seq data from BV2 cells – p<0.05, one-way ANOVA. **B.** PCA plot on mRNA seq data from BV2-dervied EVs – p<0.05, one-way ANOVA. **C.** Volcano plot showing differentially expressed genes (DEGs – mRNA species) in EVs compared to BV2 cells (p <0.05). Upregulated and downregulated numbers correspond to red dots (p< 0.05 and log2 fold change LFC>1 for upregulated genes and LFC<-1 for downregulated genes). **D.** Heatmap representation, based on enrichment Z-scores, of Gene Ontology for highly enriched mRNAs (>=2-fold change) in EVs or whole cells. (p<0.05). **E.** Volcano plots showing DEGs in cell mRNAseq data – LPS vs Control (p<0.05). Upregulated and downregulated numbers correspond to red dots (p-value < 0.05 and log2 fold change LFC > 1 for upregulated genes and LFC<-1 for downregulated genes). **F.** Volcano plots showing DEGs in EV mRNAseq data – LPS vs Control (p<0.05). Upregulated and downregulated numbers correspond to red dots (p<0.05 and log2 fold change LFC>1 for upregulated genes and LFC<-1 for downregulated genes). **G.** Comparison of 1,040 upregulated mRNAs from BV2 cells treated with LPS to 2,949 upregulated mRNAs from LPS treated BV2-derived EVs. **H.** Heatmap representation, based on enrichment Z-scores, of Gene Ontology for BV2 cells (p<0.05). **I.** Heatmap representation, based on enrichment Z-scores, of Gene Ontology for BV2-dervied EVs (p<0.05).

Pro-inflammatory genes (e.g., *Il1b, Il6*) and ontologies related to defense and immune response were increased in LPS-treated BV2 microglia. Interestingly, genes (e.g. *G3bp2*) and ontologies related to mRNA transport were also enriched in LPS treated cells (**Figure 5E**). TGF-β increased expression of several genes (e.g., *Cx3cr1, Flt1*) and ontologies related to homeostatic microglial function and biological regulation. IL-10 also increased expression of genes (e.g., *C1qc, Saa3*) and ontologies related to protein activation cascade and response to wounding and other organisms (**Additional file 7**).

Transcriptomic analyses of EVs identified over 3,000 differentially expressed genes (DEGs - mRNA species) across treatment groups (**Additional file 8**). As compared to TGF-β and IL-10, LPS polarization had the strongest impact on BV2 microglia-derived EV transcriptomes; therefore, we focused on LPS effects on the whole cell and EV transcriptomes. As discussed above, LPS polarization resulted in enrichment of pro-inflammatory genes (e.g., *Il1b, Il6)* related to immune response (**Figure 5E**). EVs from LPS-treated BV2 microglia showed increased levels of 2,949 mRNAs and decreased levels of 413 mRNAs (**Figure 5F**). Interestingly, transcriptomic alterations induced by LPS (DEGs upregulated with LPS, p<0.05, >1-fold change) on the whole cell (n=1,040) and EVs (n=2,949) showed modest overlap, with only 10% of EV transcriptomic changes overlapping with cellular changes (**Figure 5G**). 359 mRNAs showed shared differential expression in whole cells and EVs (**Figure 5G**) and majority of these DEGs were related to lipopolysaccharide-mediated signaling pathway, positive regulation of cell activation, and regulation of immune response. The 2,591 DEGs related to EV only showed enrichment of RNA metabolic process and nucleic acid binding terms. While the 682 DEGs related to cell only showed enrichment of terms such as cell cycle and mature B cell differentiation (**Supplementary Figure 3**).

To take a deeper look at LPS effects on the cellular and EV transcriptomes, we performed GSEA based on identified DEGs. We found that LPS treated BV2 cells showed enrichment in immune response, RNA binding, and metabolism, whereas we observed a decreased expression in signal transduction and GTPase activity (**Figure 5H**). In contrast, EV transcriptomes from LPS treated BV2 cells, showed enrichment in terms such as signaling and ion channel activity (**Figure 5I**). Conversely, the downregulated DEGs in EVs from LPS treated BV2 cells showed enrichment in terms such as ATPase activity and transmembrane activity.

Furthermore, we found that the majority of EV transcriptomic changes induced by LPS are not observed in the whole cell transcriptomes. This suggests that pro-inflammatory activation of microglia not only induces whole cell gene expression changes, but also likely impacts mechanisms that govern mRNA export into EVs, consistent with our observation of LPS-induced altered expression of genes involved in mRNA transport at the whole cell level (**Figure 5H**).

Taken together, these analyses identify novel transcriptomic signatures of EVs that are distinct from whole cell transcriptomes. Our studies also reveal distinct effects of pro-inflammatory activation on EV transcriptomic signatures, confirming microglial state-dependent effects on EV composition at the mRNA level, many of which do not occur at the whole-cell level.

### Microglia-derived EVs exhibit unique microRNA signatures under resting and LPS-treated conditions

We obtained sufficient RNA for sequencing of small RNA species from BV2 whole cell and EV RNA extracts. Small RNA sequencing identified ∼100 miRNAs in EVs and ∼270 miRNAs in whole cells (**Additional files 9 and 10**). In addition to miRNA, we also identified other small RNA species (circRNA, piRNA, snoRNA, snRNA, and tRNA), included in **Additional file 11**. PCA of post-filtered, lowly expressed miRNA molecules in the whole cell (PC1 63%, PC2 12% of variance) (**Figure 6A**) identified four distinct clusters based on polarization state, with the LPS effect showing the strongest effect (along PC1), a pattern similar to our mRNA-based results. PCA of post-filtered, lowly expressed miRNA molecules in the EVs (PC1 43%, PC2 16% of variance) (**Figure 6B**) also showed group-based clustering primarily based on LPS treatment.

**Figure 6:**
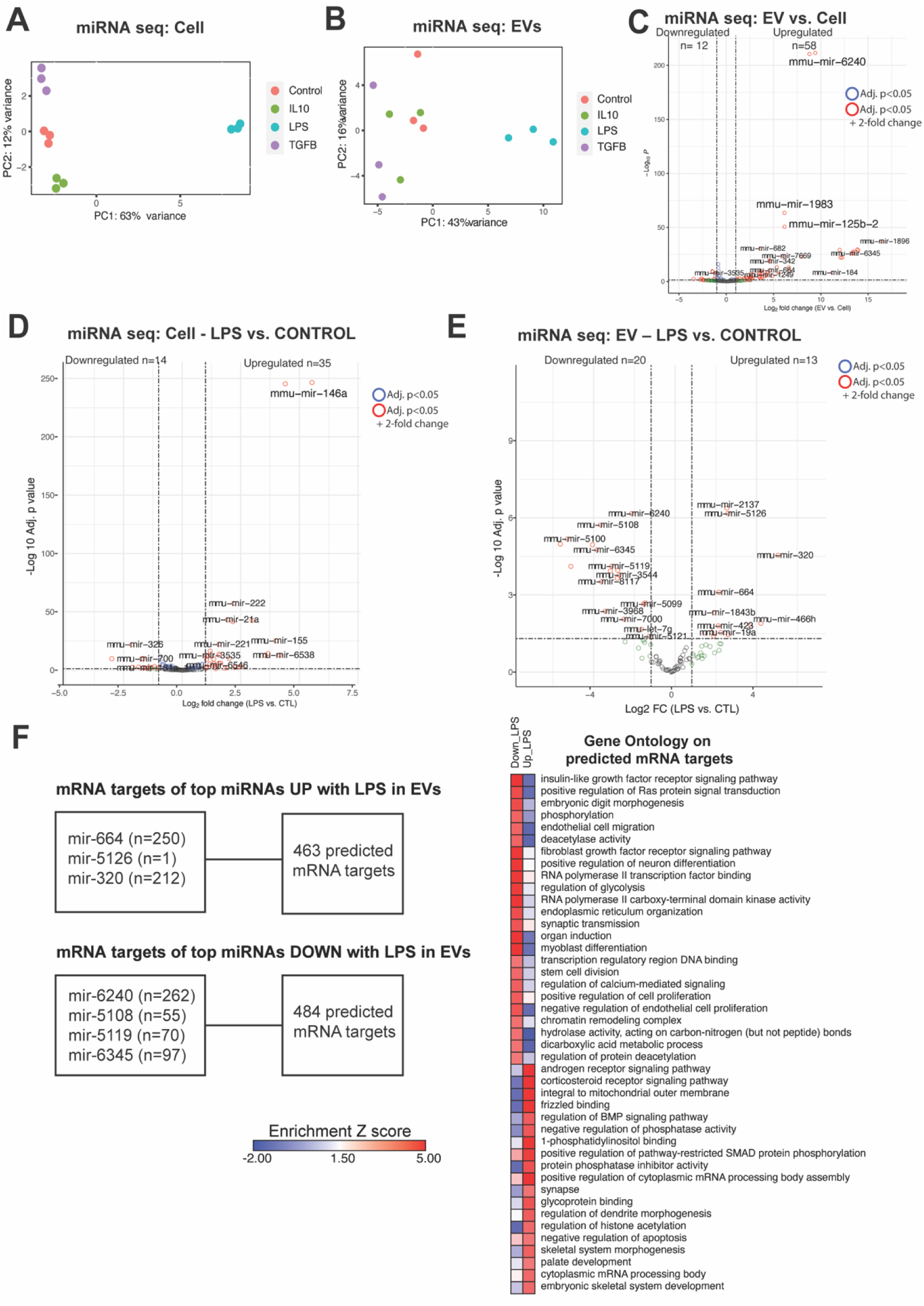
BV2 cells of distinct states and their EVs have unique microRNA signatures. **A.** PCA plot miRNA seq data from BV2 cells – p<0.05, one-way ANOVA. **B.** PCA plot miRNA seq data from BV2-dervied EVs – p<0.05, one-way ANOVA. **C.** Volcano plot – EV differentially expressed (DEX) miRNAs vs. Cell DEX miRNAs (p<0.05). Upregulated and downregulated numbers correspond to red dots (p<0.05 and log2 fold change LFC>1 for upregulated genes and LFC<-1 for downregulated genes). **D.** Volcano plot – Cell miRNAs: LPS vs. CTL (p<0.05). Upregulated and downregulated numbers correspond to red dots (p<0.05 and log2 fold change LFC>1 for upregulated genes and LFC<-1 for downregulated genes). **E.** Volcano plot – EV miRNAs – LPS vs. CTL (p<0.05) Upregulated and downregulated numbers correspond to red dots (p<0.05 and log2 fold change LFC>1 for upregulated genes and LFC<-1 for downregulated genes). **F.** Top DEX miRNAs in EVs comparing LPS vs. CTL (p<0.05 and log2 fold change LFC>2 or LFC<-2) and the number of mRNA targets (target prediction score > 80 on miRDB) for those miRNAs. Heatmap representation, based on enrichment Z-scores, of Gene Ontology for mRNA targets.

When comparing the whole cell and EV miRNA transcriptomes (**Figure 6C, Additional file 12**), we found that EVs contained very high levels of miRNAs mmu-mir-6240, mmu-mir-1983, and mmu-mir-1896. Interestingly, mir-6240 has been previously reported to be found at higher than expected levels in EVs compared to cell, in both human and mouse samples^31^.

Focusing on LPS treatment effects, we observed that LPS-treated BV2 cells and their EVs enrich for unique miRNA signatures. More specifically, we see increased expression of mmu-mir-320, mmu-mir-5126, and mmu-mir-466h. Interestingly, mir-320 has been found to be upregulated in plasma EVs from individuals at high risk of lung cancer^32^. Furthermore, one study found that mmu-mir-5126 was dysregulated in IFN-γ primed mesenchymal stromal/stem cells (MSC) and mmu-mir-466q found in MSC-derived exosomes modulated the pro-inflammatory phenotype of activated N9 microglia cells^33^. Overall, these results implicate that miRNAs in EVs from LPS treated BV2 cells could play a role in modulating inflammation.

Given that miRNA binds with target mRNA to direct gene silencing via mRNA cleavage or translation^34^, we used the miRDB^35^ online database to generate a list of predicted targets from our differentially expressed miRNAs in EVs following LPS treatment. We found that significantly enriched miRNAs that are increased with LPS treatment in EVs (mir-664, mir-5126, and mir-320) had 462 predicted targets (target prediction score > 80) which showed an enrichment in ontologies related to androgen and corticosteroid receptor signaling pathway and cytoplasmic mRNA processing pathway (**Figure 6F**). On the other hand, we found that significantly enriched miRNAs that are decreased with LPS treatment in EVs (mir-6240, mir-5108, mir-5119, and mir-6345) had 487 predicted targets (target prediction score > 80) which showed an enrichment in ontologies related to regulation of cell proliferation, chromatin remodeling structure and regulation of glycolysis. Overall, these results demonstrate that EVs from LPS activated BV2 cells could contain a set of miRNAs that target distinct mRNAs, thus modulating gene expression in a responder cell.

### EVs derived from LPS-treated microglia induce pro-inflammatory changes in responder microglia

Our results from proteomic and transcriptomic characterization of microglia-derived EVs demonstrate EV-specific molecular changes induced by microglial polarization state, particularly by LPS induced pro-inflammatory activation. Since EVs carry these protein and RNA cargo to other cells, we hypothesized that state-dependent alterations in EV composition by LPS can relay inflammatory signals to responder cells and impact their transcriptomic state. To accomplish this, we isolated EVs from untreated and LPS-treated BV2 microglia, estimated EV concentration by NTA, and then dosed resting responder BV2 microglia with equal amounts of EVs and assessed the transcriptomic alterations induced by EVs using RNA seq. To ensure that LPS (endotoxin) contamination of EV preparations does not confound these EV transfer assays, we measured endotoxin levels in cell culture supernatants as well as in EV preparations. Cell culture supernatants from LPS-treated BV2 microglia contained 80-90 ng/mL of LPS, consistent with the dose of 100 ng/mL that was added. EV preparations from control and LPS-treated BV2 cells had equal endotoxin levels (approximately 4ng/mL, p=0.511060, not significant), confirming that prior to addition to responder cells, any endotoxin contamination in the EV preparations was equal in both groups. EVs were dosed at 2,000,000 EVs/well (2mL of media/well) across all conditions, and the estimated final endotoxin concentration added to responder cells was minimal (p=0.103, not significant) and also similar in control and LPS-treated conditions (**Figure 7B**). These results confirm that any potential effects of EVs in our transfer experiments cannot be explained by spurious endotoxin contamination of EV preparations.

**Figure 7:**
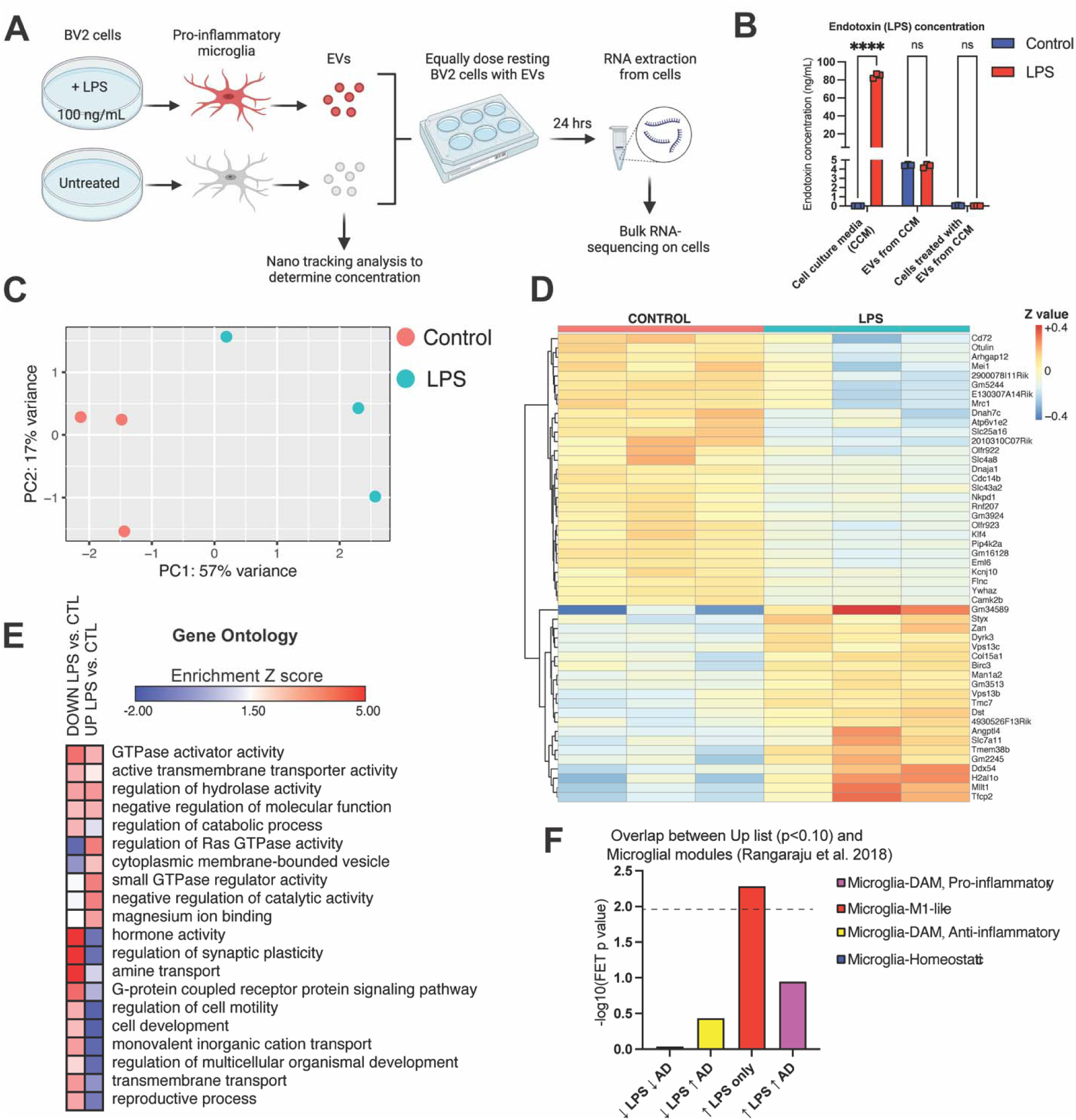
State specific effects of LPS treated microglia-derived EVs on resting BV2 cells. **A.** Illustration outlining experimental setup. Created with BioRender. **B.** The endotoxin concentrations (ng/mL) of cell culture media (untreated and treated with LPS 100ng/mL) p<0.000001, significant, EVs from CCM (untreated and treated with LPS) p=0.511, not significant, and cells treated with EVs from CCM (untreated and treated with LPS) p=0.103, not significant. **C.** PCA plot on bulk RNA seq data from responder cells dosed with EVs derived from LPS treated BV2 cells or Control BV2 cells – p value <0.05, one-way ANOVA. **D.** Heatmap of DEGs in responder cells (LFC < or >0, p<0.05). **E.** Heatmap representation, based on enrichment Z-scores, of Gene Ontology in responder cells (LFC < or >0, p<0.10). **F.** FET Analysis demonstrating overlap between Up list (p<0.10) and microglia microarray analysis in Rangaraju et al. 2018. Bar color indicates microglial module color from referenced paper.

PCA of post-filtered, lowly expressed genes showed 74% of variance was explained by 2 PCs, of which PC1 explained 57% of variance while PC2 explained 17% of variance (**Figure 7C**). The PCA identified two distinct clusters. RNA sequencing analyses of responder BV2 microglia exposed to control or LPS-treated BV2-derived EVs identified 148 genes that were differently expressed at the FDR<0.10 level (n=71 increased DEG including *Ddx54* and *Mllt1*; n=73 decreased DEGs including *Atp6v1e2* and *Cd72*) (**Figure 7D, Additional file 13**). GSEA revealed enrichment of terms such as G-protein coupled receptor protein signaling and regulation of synaptic plasticity for the down list; whereas, there was enrichment of GTPase activity for the up list (**Figure 7E**).

We also conducted FET analyses to determine whether gene expression changes induced by transfer of EVs from pro-inflammatory microglia overlap with specific homeostatic, pro-inflammatory and disease-associated microglial gene signatures previously identified by weighted gene co-expression network analyses of microglial transcriptomes^6^ (**Figure 7F**). We found that genes that were increased in responder cells following exposure to EVs from LPS-polarized microglia were enriched (unadjusted FET p=0.005) in a LPS-induced pro-inflammatory gene module (Red Module), including genes *4932438A13Rik, Dst, Fyb, Gnai, Heatr6, Slc25a22, and Tmem72*. These results from EV transfer experiments demonstrate the ability of EVs to relay the inflammatory state of microglia of origin to responder microglia.

### Conclusions

Microglia, the resident immune cell type of the brain, play an essential role in mediating inflammatory responses in the CNS. Microglia can alter their morphology, molecular profile, and function in response to immune activators, thus exerting different functions in different diseases^2^. Microglia-mediated neuroinflammation is a key pathological component of several neurodegenerative diseases^3^. Compelling evidence has suggested that EVs can have an indirect role in regulation of inflammatory signals and propagation of pathogenic cargo^14, 36^. Since microglia demonstrate heterogenous states in response to stimuli and signals from other brain cell types, it is possible that microglia-derived EVs may also exhibit state-related heterogeneity. However, the characterization of microglia-derived EVs at the proteomic and transcriptomic level, and how these are impacted by microglial state, have not been well studied. In the present study, we have characterized the proteomic and transcriptomic signatures of EVs from distinct microglia states using in vitro models. Our results demonstrate that upon treatment with LPS, TGF-β, or IL-10 to elicit distinct microglial states, BV2 cells and their EVs indeed demonstrate unique proteomic and transcriptomic signatures; indicating polarization of BV2 cells impacts the cargo of EVs. LPS treatment, in particular, had the most profound impact on transcriptomic compositions of microglia-derived EVs, and interestingly, these changes in EV mRNA cargo were not apparent at the whole cell level, suggesting unique effects of LPS on mechanisms that direct mRNA to EVs. Lastly, EVs derived from LPS-activated microglia were able to induce pro-inflammatory transcriptomic changes in resting responder microglia, confirming the ability of microglia-derived EVs to relay functionally-relevant inflammatory signals. Collectively, our results provide a critical resource of microglia-specific EV signatures, many of which are distinct from EVs from other cell types. State-specific EV signatures at the levels of proteins, mRNA, miRNA as well as other non-coding RNA species, will serve as important resources for the fields of EV biology as well as neuroscience.

The microglial EV proteomes in our study are also deeper than current microglial EV proteomes, identifying 494 proteins not previously reported. Over 207 proteins in microglia EVs were also identified that have not been reported in proteomic studies of EVs from other cell types. This increased depth of the microglia EV proteome was not due to non-EV contamination, because we verified EV enrichment via several complimentary methods, including TEM, Western blotting, as well as high-level enrichment of canonical EV surface proteins such as CD9. Our proteomic studies of EVs identified unique functional groups of proteins as compared to the whole cell proteome of BV2 microglia. We found that BV2 microglia-derived EVs were enriched in ontologies related to positive regulation of transporter activity, regulation of RNA splicing, cytosolic large ribosomal subunit, and aminoacyl-tRNA synthetase complex in comparison to non-microglial EV proteomes. This suggests that proteins contained within microglia-derived EVs could play a role in mRNA processing as well as protein synthesis in the recipient cell. Furthermore, Oh et al. 2022 reported that Aminoacyl tRNA synthetase (ARS) complex-interacting multifunctional protein 1 (AIMP1), a structural component of the multienzyme ARS complex, can induce microglial activation and has been associated with several inflammatory diseases^37^. Based on this, microglia specific-EV signatures could be related to protein-synthesis machinery and influence activation of recipient cells.

Another major finding in our study is that microglial state influences EV proteomic composition. Specifically, we show that EVs derived from BV2 cells treated with IL-10 and TGF-β demonstrated a reduction in cytosolic chaperonin Cct ring complex proteins (e.g., Cct3, Cct4). Given that IL-10 is supposed to induce a protective response in microglia cells and TGF-β a homeostatic response, we would expect a decrease abundance in proteins related to pathogenic pathways. Previous studies have shown that chaperonins Cct subunits show increased quantities in disease states. For example, studies with glioblastomas have demonstrated an increase in the Cct subunits in EVs derived from tumor tissues^38^. Since Cct proteins (as part of the TRiC complex) also participate is regulation of protein folding of actin and tubulin as well as pathological protein aggregation, it is also plausible that CCT changes in microglia-derived EVs could impact pathogenic mechanisms of neurodegeneration^39^. The results of these studies align well with the reduction in Cct ring complex proteins following treatment with IL-10 and TGF-β. Furthermore, we found that EVs derived from LPS activated BV2 microglia demonstrated an enrichment in proteins related to proteasome activity (e.g., PSMB3). Prior work has demonstrated that proteasome systems are involved in regulating pro-inflammatory pathways^40^. Thus, EVs derived from LPS-activated microglia can potentially enhance inflammation through enrichment of proteasomes. These findings together with our results suggest that microglial EVs from distinct microglia states have unique proteomic profiles.

Beyond elucidating proteomic features of microglia-derived EVs, we also comprehensively characterized RNA cargo of microglia-derived EVs and the corresponding RNA content from whole cells as well. Our RNA sequencing studies (mRNA and miRNA) showed that microglia-derived EVs had a relatively larger number of miRNA species relative to mRNA, when compared to whole cell mRNA and miRNA profiles. This aligns with the general consensus that smaller RNAs are more abundant than mRNAs in exosomes. Interestingly, following LPS activation of BV2 cells we noticed an increase in mRNA transcripts in EVs compared to cells. Furthermore, we found that LPS activated BV2 microglia demonstrated an enrichment in genes related to mRNA transport and RNA binding. Given the accumulating evidence showing that mRNAs in EVs could serve as templates for novel protein translation in recipient cells^41, 42^, it is possible that there is a specific mechanism involved in the shuttling of mRNAs into EVs that is increased following LPS microglial activation. We also found that a majority of EV transcriptomic changes induced by LPS activation are not observed in the whole cell transcriptomes. This suggests that pro-inflammatory activation of microglia not only induces whole cell gene expression changes, but also likely impacts mechanisms that govern mRNA export into EVs. However, further investigation is necessary to determine whether these mRNAs in EVs are translated into proteins in responder cells thereby influencing the phenotype of the cell.

Next, we sought to explore the miRNA contents of microglia-derived EVs given their role in regulating the expression of specific gene targets^18^. In our studies, we observed that LPS-treated BV2 microglia and their EVs bear unique miRNA signatures that may be important in modulating inflammation. Specifically, mir-320, which was increased in EVs from LPS-treated BV2 cells, has been previously found to be upregulated in the cortex of human sporadic AD cases^43^. Furthermore, we identified several miRNAs that are contained at very high levels in EVs but not cells (e.g. mir-6240 and mir-6236). Focusing on the miRNAs that are differentially expressed in EVs from the LPS treated group versus control, we found that miRNAs in the EVs from LPS treated group target unique mRNAs compared to control, which are involved in androgen and corticosteroid receptor signaling pathway and the cytoplasmic mRNA processing pathway. By targeting distinct mRNAs, the miRNA cargo in these EVs could impact gene expression and modulate inflammation in a recipient cell. However, further mechanistic studies are needed to test this hypothesis.

The complementary multi-omics characterization of microglia-derived EVs and microglial state-dependent effects on EVs suggest that EVs from distinct microglia states have unique protein and RNA cargo, which may be able to impact other cells. Therefore, we hypothesized that state-dependent alterations in EV composition by LPS can relay inflammatory signals to responder cells and impact their transcriptomic state. To accomplish this, we dosed resting responder BV2 microglia with equal amounts of EVs from untreated and LPS-treated BV2 microglia and then assessed the transcriptomic alterations induced by EVs using RNA seq. Our RNA sequencing analyses revealed that responder BV2 microglia exposed to LPS-treated BV2- derived EVs display upregulation of genes related to GTPase activity and downregulation of genes related to G-protein coupled receptor protein signaling and regulation of synaptic plasticity. Mukai et al. reported that LPS stimulation significantly increased Rho-GTPase activity levels in microglia^44^. These results from EV transfer experiments demonstrate the ability of EVs to relay the inflammatory state of microglia of origin to responder microglia. Although these transcriptomic effects were not very large, the overall effect of LPS-microglia-derived EVs on responder microglia was consistent with a pro-inflammatory profile previously reported in mouse microglia. Importantly, we confirmed that the observed effects of EVs cannot be explained by passive transfer of LPS itself. We attribute the smaller effect size on responder transcriptomes to dosing of EVs and duration of exposure.

Despite the strengths of our in vitro studies, some limitations of our work should be considered. For our study we chose to use BV2 microglia cells, a well characterized immortalized murine microglial cell line^45^, because of their ability to capture major cellular phenotypes of microglia, suitability as an alternative model for primary microglia culture^46^, and feasibility with obtaining sufficient EVs for EV proteomics and transcriptomics analyses. However, it should be noted that BV2 cells represent transformed cells which could change their phenotype compared to primary microglia cells in the central nervous system^47^. Therefore, further studies will be needed in either human iPSC-induced microglia cells or primary mouse microglia to confirm that there are similar proteomic and transcriptomic changes as observed in our BV2 cell studies. While we chose untreated BV2 microglia as responder cells for our EV dosing studies, other cell types (e.g. neurons and astrocytes) may be considered in future studies to understand the effects of microglia-derived EVs on non-microglial cellular phenotypes. Another potential limitation of our study is the sensitivity to detect low quantities of protein, mRNA, and miRNAs from EVs; however, we employed several steps in our experimental design and data analysis pipeline to account for low input.

To conclude, our findings suggest EVs from distinct microglia states have unique proteomic and transcriptomic profiles. Furthermore, we have identified novel EV proteins in microglia not seen in other EVs, thereby increasing the depth of the microglia-derived EV proteome that has not been previously reported on. Through our transcriptomic analysis, we discovered that LPS activation of BV2 cells has the strongest impact on EV composition, possibly increasing the amount of mRNAs transported and packaged into the EVs. Lastly, our data suggests that EVs from LPS-activated microglia can elicit a transcriptomic change in resting recipient microglia that mimics that of a pro-inflammatory response. Our study highlights the value of characterizing the cargo of EVs from distinct cell types. EVs are a promising therapeutic as well as diagnostic tool for neurodegenerative disease. Given that microglia are highly dynamic cells and can polarize to multiple phenotypes in response to their environment, it is to be inquired if their EVs might have distinct functions and contents. Therefore, the uptake of microglia-derived EVs by other cells could mediate indirect downstream signaling events. More research is needed to understand the mechanisms of microglia-derived EVs in neurodegenerative disease, specifically whether the presence or absence of certain cargo can impact disease progression.

## Abbreviations

EV: Extracellular vesicles
CNS: Central nervous system
AD: Alzheimer’s disease
LPS: Lipopolysaccharide
IL-10: Interleukin-10
TGF-β: Transforming growth factor beta
TEM: Transmission electron microscopy
NTA: Nano tracking analysis
LFQ-MS: Label free mass spectrometry
mRNA: messenger RNA
miRNA: micro RNA
TLR: Toll-like receptor
TNF: Tumor necrosis factor
CCT: Chaperonin containing TCP1 (CCT/TRiC)
GSEA: Gene set enrichment analysis
GO: Gene ontology
KEGG: Kyoto Encyclopedia of Genes and Genomes
DEP: Differentially expressed protein
DEG: Differentially expressed gene
DEX: Differential expression
PCA: Principal component analysis

## Methods

### Antibodies, buffer, and reagents

A complete table of antibodies & reagents are provided (Tables 1 & 2).

**Table 1.**
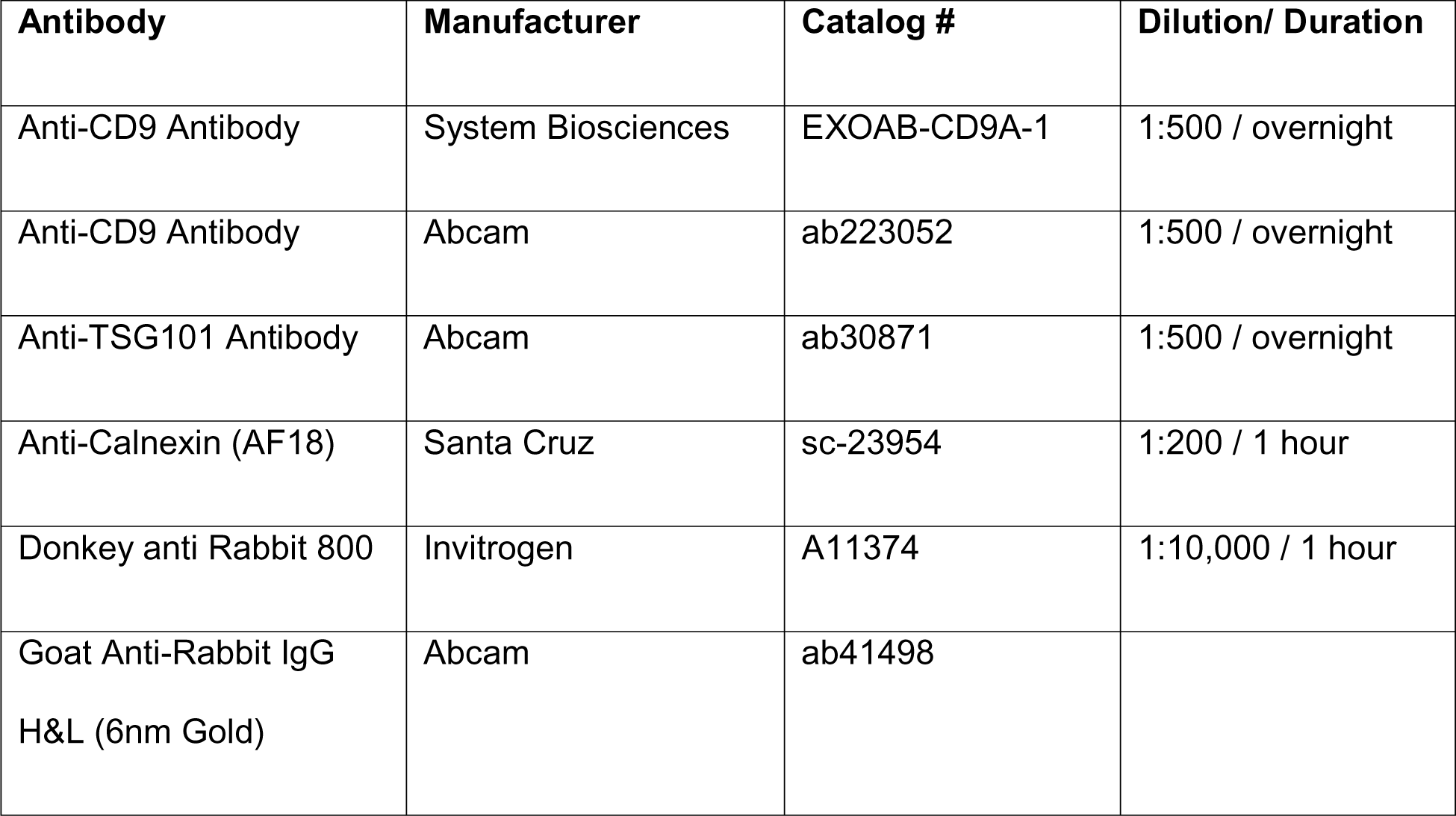
Antibodies used and their corresponding dilutions.

**Table 2.**
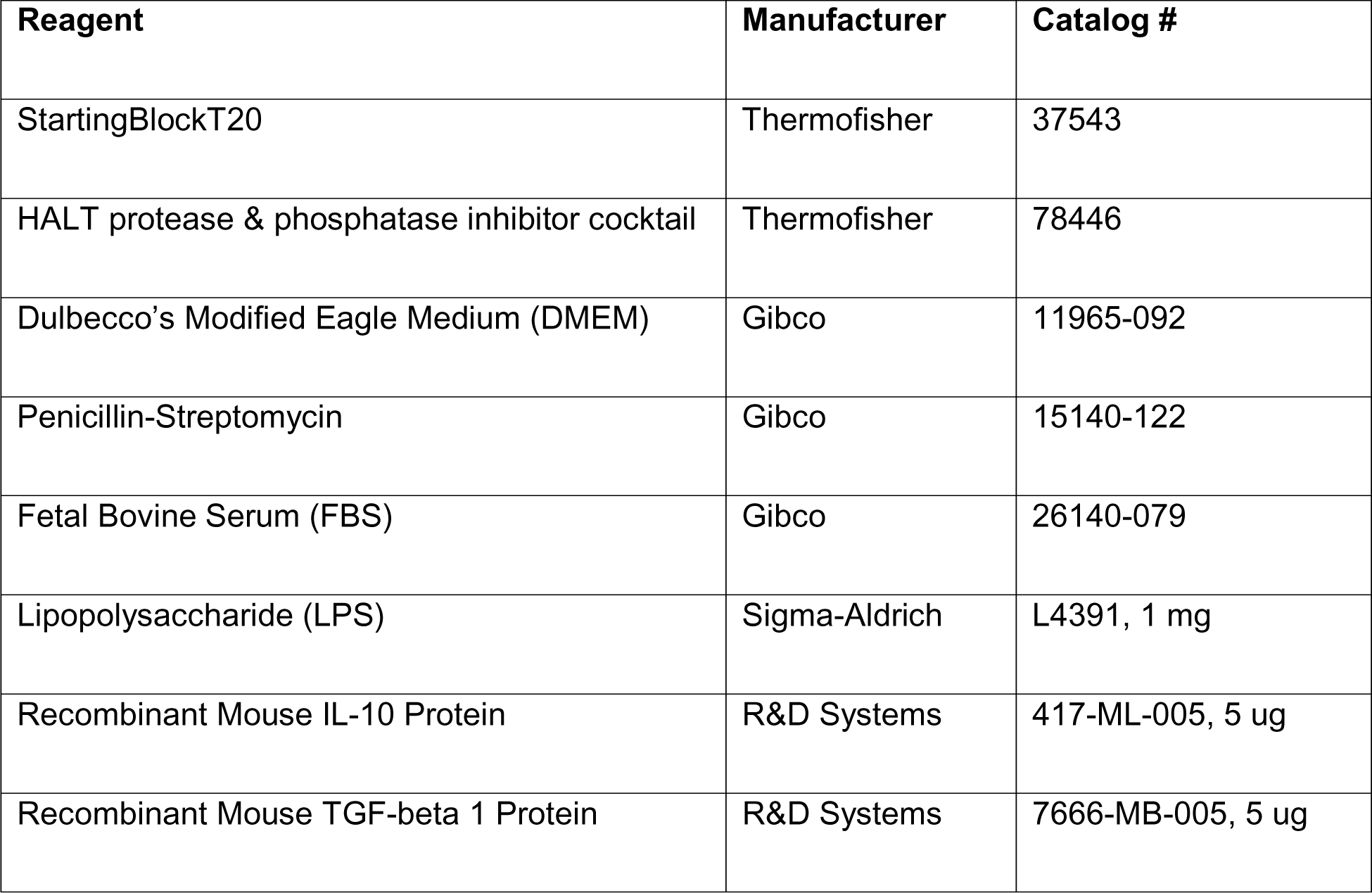
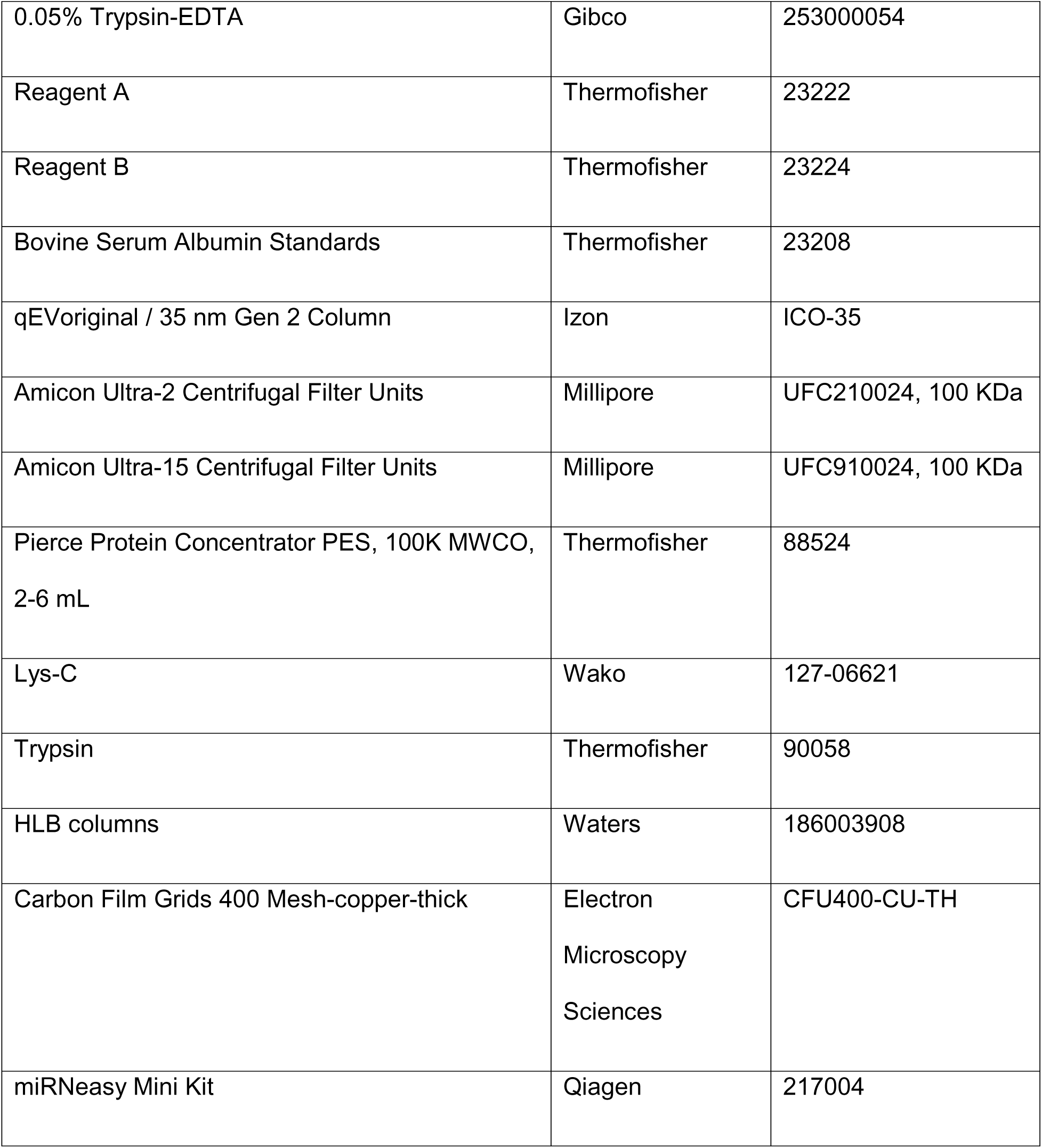
Reagents used and their manufacturer and catalog numbers.

### Cell culture studies

BV2, an immortalized murine microglial cell line, were cultured in filtered Dulbecco’s Modified Eagle Medium (DMEM) supplemented with high glucose and L-glutamine containing 1% penicillin/streptomycin, and 10% Fetal Bovine Serum (FBS). All media was vacuum-filtered with 0.2 µm filters. The cells were incubated at 37 degrees Celsius (°C) and 5% CO2 until reaching 80% confluency. The splitting regimen took place twice weekly, plating one million cells onto a 100 mm culture plate to a final volume of 12 mL culture media. In preparation for experiments, one million cells were plated in 100 mm plates with 12 mL of media. 24 hours after plating, existing media was swapped for filtered serum-free media (DMEM containing 1% penicillin/streptomycin) and BV2 cells were treated with either TGF-β (50 ng/mL), IL-10 (50 ng/mL), and LPS (100 ng/mL) for 72 hours^24–26^. Control cells were left untreated. For dosing resting recipient cells, EVs were isolated from cell culture media following treatment of BV2 as described above. The concentration of isolated EVs for each sample were determined using NTA. Recipient BV2 cells (n=3/condition) in a 6 well plate were dosed for 24 hours with 2 million EVs/well from either TGF-β, IL-10, or LPS treated BV2 cells. Cell culture media in 6 well plates was swapped with serum-free media prior to dosing recipient cells with EVs.

After 72 hours in culture, cell culture medium was collected from the plates, transferred into 15 mL tubes, and placed on ice. Next, the plates were washed twice with ice-cold 1x phosphate buffer saline (PBS). Cell pellets were harvested in 500 μL Urea lysis buffer (8 M Urea, 10 mM Tris, 100 mM NaH2PO4, pH 8.5) with 1x HALT protease & phosphatase inhibitor cocktail without EDTA. Cell lysates were then sonicated at 30% amplitude thrice in 5-second on-off pulses to disrupt nucleic acids and cell membrane. All cell lysates were centrifuged at 4°C for 2 minutes at 12,700 rpm. The supernatants were transferred to a fresh 1.5 mL LoBind Eppendorf tube. The protein concentrations of whole cell lysates were determined by Bicinchoninic acid (BCA) assay reagents using Bovine Serum Albumin Standards.

### EV isolation

EV isolation was conducted as per Izon manufacturer’s protocol^48^. Cell culture media underwent several centrifugation steps to remove cellular debris (10 min at 500xg and 10 min 10,000xg). After each spin, the supernatant was collected and subjected to the subsequent spin. The final supernatant was collected and added to an Amicon Ultra-15 Centrifugal Filter (molecular weight cut-off 100 kDa) to concentrate the sample and then was added to a qEV column resin column for size exclusion chromatography using the Izon qEV system with a qEVoriginal / 35 nm Gen 2 Column. Sterile PBS was used as the flushing buffer. The second fraction containing the most abundant amount of EVs was then concentrated using a Amicon Ultra-2 Centrifugal Filter (molecular weight cut-off 100 kDa) or Pierce Concentrator 0.5mL, PES (molecular weight cut-off 100 kDa). The resulting concentrate was used for any downstream analysis (TEM, immunoblotting, NTA, proteomics, or transcriptomic analyses).

### Transmission Electron Microscopy (TEM) of EVs

For electron microscopic analysis, 5 μL from the EV samples were loaded onto the carbon side of charged copper/carbon-coated electron microscopic grids. After 5 minutes, sample loaded grids were washed 3 times in distilled water and then stained with 1-3% uranyl acetate for 1 minute in the dark. Once grids were dry, EVs were observed under TEM at 80 kV. TEM grids were stored in the appropriate grid storage boxes for future use. Hitachi HT7700 transmission electron microscope operating at 80 kV was used for imaging.

#### Immunogold labeling of EVs with CD9

For the preparation of samples for immunogold electron microscopy, purified EVs were fixed for 1 hour in a mixture of 0.1% glutaraldehyde, 2.5% paraformaldehyde, 0.03% picric acid, and 0.03% CaCl2 in 0.01 M cacodylate buffer at a pH of 7.2. Fixed samples were then adsorbed for 1 hour to freshly glow discharged copper Formvar-coated grids. For immunogold staining, grids with adsorbed EVs were incubated sample-side down on drops of blocking buffer for 30 minutes (a mixture of 0.1% BSA and 0.01 M glycine in 0.01 M PBS). They were then transferred to drops containing rabbit polyclonal antibody to CD9 (Abcam ab223052) diluted in diluent buffer (1% BSA in 0.01 M PBS) at a dilution of 1:20 and incubated for 1 hour at room temperature and then overnight at 4°C. Next, grids were washed through a series of drops of blocking buffer and subsequently transferred to drops containing gold-labeled secondary antibodies. Secondary antibodies were goat polyclonal antibody to rabbit IgG labeled with 6-nm gold particles (Abcam ab41498), at a dilution of 1:20 in diluent buffer. After EVs were incubated with secondary antibodies for 1 hour at room temperature, they were once again washed by transferring them through a series of drops of blocking buffer, then through a series of drops of 0.01 M PBS, and finally, through a series of drops of deionized water. Grids were then fixed for 15 min on drops of 2.5% aqueous glutaraldehyde. EVs were subsequently negatively stained with 2% aqueous uranyl acetate and examined in a JEOL JEM 1400 transmission electron microscope operated at 80 kV.

### Nano tracking Analysis (NTA) of isolated EVs

The size and total number of EVs were measured by using NanoSight NS300 (Malvern, UK) with the technology of Nanoparticle Tracking Analysis (NTA). Size distribution and concentration of EVs in an aqueous buffer was obtained by utilizing Brownian motion and light scattering properties^49^. Samples were diluted with 1X PBS to obtain optimal concentration for detection (10^6^–10^9^ particles/ml) and injected with a continuous syringe system for 60 sL×L3 times at speed 100 μl/min. Data acquisition was undertaken at ambient temperature and measured 3 times by NTA. Data were analyzed with NTA 3.2 software (Malvern, UK) with minimum expected particle size 10 nm.

### Immunoblotting studies

In each well, 15 µg of protein from cell lysates and 21 µL of EV lysate resolved in a 4-12% polyacrylamide gel and transferred onto iBlot 2 Transfer Stack containing nitrocellulose membrane using the BOLT transfer system. The membranes incubated for 1 hour at room temperature in StartingBlockT20 before receiving rabbit anti CD9 primary antibody overnight at 4 °C. After primary antibody incubation, the membranes underwent three 5-minute washes with 1x TBS-T. The membranes incubated for 1 hour at room temperature in a secondary antibody cocktail of donkey anti rabbit 800. The membranes were then washed again as previously described before undergoing imaging via the Odyssey infrared Imaging System (LI-COR Biosciences).

### Proteomic studies of BV2 microglia and EVs

#### Protein digestion

Sample preparation for MS was performed according to our laboratory protocols modified from previous publications^50–52^. To prepare enriched samples for mass spectrometry analysis, EV fractions and cells were lysed in 8M urea containing protease and phosphatase inhibitors. 100 µg of protein from the cell lysates and the entire volume of the EV lysates were then reduced with 1 mM dithiothreitol (DTT) at room temperature for 30 min and alkylated with 5 mM iodoacetamide (IAA) in the dark for 30 min with rotation. Proteins were digested overnight with 2 µg of lysyl (Lys-C) endopeptidase (Wako, 127-06621) at RT on shaker. Samples were then diluted (7-fold) with 50 mM ammonium bicarbonate (ABC) to bring the urea concentration to 1 M. Samples were then digested overnight with 2 µg of trypsin (Thermo, 90058) at RT on shaker. The resulting peptide solutions were acidified to a final concentration of 1% formic acid (FA) and 0.1% triflouroacetic acid (TFA), desalted with a HLB columns (Waters, 186003908), and dried down in a vacuum centrifuge (SpeedVac Vacuum Concentrator).

#### Mass spectrometry (MS)

Dried peptides were resuspended in 15 μL of loading buffer (0.1% FA and 0.03% TFA in water), and 7–8 μL was loaded onto a self-packed 25 cm (100 μm internal diameter packed with 1.7 μm Water’s CSH beads) using an Easy-nLC 1200 or Dionex 3000 RSLCnano liquid chromatography system. The liquid chromatography gradient started at 1% buffer B (80% acetonitrile with 0.1% FA) and ramps to 5% in 10 s^53^. The spectrometer was operated in data dependent mode in top speed mode with a cycle time of 3 s. Survey scans were collected in the Orbitrap with a 60,000 resolution, 400 to 1600 m/z range, 400,000 automatic gain control (AGC), 50 ms max injection time and RF lens at 30%. Higher energy collision dissociation (HCD) tandem mass spectra were collected in the ion trap with a collision energy of 35%, an isolation width of 1.6Lm/z, AGC target of 10000, and a max injection time of 35 ms. Dynamic exclusion was set to 30 s with a 10Lppm mass tolerance window.

#### Protein identification and quantification

MS raw files were searched using the search engine Andromeda, integrated into MaxQuant, against 2020 mouse Uniprot database (91,441 target sequences). Methionine oxidation (+15.9949LDa) and protein N-terminal acetylation (+42.0106LDa) were variable modifications (up to 5 allowed per peptide); cysteine was assigned as a fixed carbamidomethyl modification (+57.0215LDa). Only fully tryptic peptides were considered with up to 2 missed cleavages in the database search. A precursor mass tolerance of ±20Lppm was applied prior to mass accuracy calibration and ±4.5Lppm after internal MaxQuant calibration. Other search settings included a maximum peptide mass of 4600LDa, a minimum peptide length of 6 residues, 0.05LDa tolerance for orbitrap and 0.6LDa tolerance for ion trap MS/MS scans. The false discovery rate (FDR) for peptide spectral matches, proteins, and site decoy fraction were all set to 1 percent. Quantification settings were as follows: re-quantify with a second peak finding attempt after protein identification has completed; match MS1 peaks between runs; a 0.7 min retention time match window was used after an alignment function was found with a 20 min RT search space. Quantitation of proteins was performed using summed peptide intensities given by MaxQuant. The quantitation method only considered razor plus unique peptides for protein level quantitation.

The MaxQuant output data were uploaded onto Perseus (Version 1.6.15) for analyses. The categorical variables were removed, and intensity values were log (base 2) transformed. The data were filtered based on 50% missingness values and missing values were further imputed from normal distribution (width: 0.3, down shift: 1.8). The MaxQuant output data (LFQ intensities) from the EV samples were first normalized based on column sum weight (column sum of given sample divided by average sum across all samples) for each sample to account for volumetric loading of EV samples. The data was then processed as described above with Perseus.

#### Data Analysis related to proteomic studies

Proteomic data analysis involved several approaches including differential expression analysis, gene ontology (GO) analysis, principal component analysis (PCA), and hierarchical clustering using the average linkage method with one minus Pearson correlation. To visualize the data, heat maps of the normalized data were generated using Morpheus from the Broad Institute (https://software.broadinstitute.org/morpheus), and additional graphical representations were created using R software (version R-4.2.0) and Prism (GraphPad, version 9) software.

#### Cell Proteome: Data Normalization, Log Transformation, and Filtering

LFQ intensities and raw intensity values were uploaded onto Perseus (version: 1.6.15) for analyses. Categorical variables were removed, LFQ intensity values were log-2 transformed, and data were in general filtered based on 50% missingness across the group of samples that were selected for each analysis. Missing values were imputed from normal distribution.

#### Cell Proteome: PCA and Differential Expression Analysis

The inputted data was then preprocessed to remove duplicates. The code used for this preprocessing is referenced from the Supplemental materials. Post processing, the inputted data file from the cell proteome contained 1882 proteins. To identify patterns, relationships, and important variables in the dataset and gain insight into the main sources of variation, we performed principal component analysis (PCA), a widely used statistical technique for analyzing high-dimensional datasets. To identify significantly differentially enriched proteins (with an unadjusted p-value ≤ 0.05), we conducted an unpaired t-test comparing the treatment groups against the control group. Furthermore, to identify proteins of interest that were differentially expressed, we performed a one-way analysis of variance (ANOVA) on all samples (n = 16), comparing the control, LPS, IL-10, and TGF-β groups (n = 4 each). The code for the one-way ANOVA analysis was adapted from the “parANOVA” repository on GitHub (https://github.com/edammer/parANOVA). Based on the one-way ANOVA calculation, we calculated the PCA (https://rdrr.io/bioc/M3C/src/R/pca.R) for the differentially expressed gene “cleanDat.” To determine whether a gene was differentially expressed, we used a threshold of a p-value < 0.05 in the one-way ANOVA analysis. Out of the 1882 proteins in cleanDat, we identified 333 proteins that met this criterion.

Additionally, we generated one-way ANOVA significant proteins and created heatmap (https://nmf.r-forge.r-project.org/aheatmap.html) and PCA plots to identify patterns in the dataset. Differentially enriched proteins were presented as volcano plots. Overall, these methods allowed us to analyze the proteomic data, identify differentially enriched and differentially expressed proteins, and gain insights into the underlying structure and variation within the dataset.

#### EV Proteome: Data Normalization, Log Transformation, and Filtering Missingness

Before conducting any analysis, protein abundances in the EV samples were normalized. This involved calculating the column sum (LFQ intensities) for each sample in the raw file and determining weights by dividing the column sum of a given sample by the average sum across all samples. The LFQ data was then normalized to the column sum weight for each sample.

Following normalization, LFQ intensities and raw intensity values were uploaded onto Perseus (version: 1.6.15) for analyses. Categorical variables were removed, LFQ intensity values were log-2 transformed, and data were in general filtered based on 50% missingness across group of samples that were selected for each analysis. Missing values were imputed from normal distribution.

#### EV Proteome: Batch Regression, Differential Expression Analysis, and PCA

The EV analysis involved 533 proteins divided into Control, TGF-β, IL10, and LPS groups, each with four samples. For statistical analysis, batch regression was performed using bootstrap regression to minimize the impact of batch on sample variance. Variance partitioning was employed to generate violin plots illustrating the contributions of treatment and batch to the observed variance in the data. The primary objective was to reduce the variance caused by batch and eliminate the batch effect. Volcano plots were then generated to visualize the differentially enriched or depleted proteins. The regressed data, obtained after applying batch regression, was used in an unpaired t-test comparing the control group to the treatment groups. The top proteins identified from the volcano plots were further examined for their ontologies. Principal component analysis (PCA) was conducted, incorporating one-way ANOVA (P < 0.05) and batch regression. Throughout the EV analysis, the “parANOVA” repository on GitHub (https://github.com/edammer/parANOVA) was referenced for the one-way ANOVA code. These analyses allowed for a comprehensive investigation of the proteomic data, encompassing normalization, statistical tests, visualization, ontological analysis, and enrichment analysis, providing valuable insights into the characteristics and differential expression patterns of the proteins under study.

#### Gene Set Enrichment Analysis (GSEA)

Gene set enrichment analysis (GSEA) was conducted using AltAnalyze (http://altanalyze.org) (Version 2.0). Three treatment groups were compared to the Control group. Specific parameters were employed for the analysis. Additionally, differentially enriched proteins with unadjusted p-values ≤ 0.05 and fold change ≥ 2 from the differential analyses were included in the input lists. The background gene list consisted of unique gene symbols for all proteins identified and quantified in the mouse brain.

### Transcriptomic studies of BV2 microglia and EVs

#### RNA extraction, library prep, and RNA sequencing

Total RNA was extracted from BV2 cells and BV2 cell-derived extracellular vesicles using the Qiagen miRNeasy Mini Kit (Cat. No. 217004). 20 µL of EV fraction was resuspended in 700 µL of TRIzol lysis buffer. BV2 cell pellets were harvested in 700 μL of TRIzol lysis buffer. Added 140ul of chloroform was added to each sample and shook vigorously. Samples were then centrifuged for 15 min at 12,000 x g at 4°C. After centrifugation, the sample separated into 3 phases: an upper, colorless, aqueous phase containing RNA; a white interphase; and a lower, red, organic phase. The top aqueous phase was carefully isolated without disturbing the other phases. The top aqueous phase was transferred to a new collection tube and 1.5 volumes of 100% ethanol was added and mixed by pipetting up and down. 700 μL of the sample was added to the provided spin column in collection tubes. Samples were spin at ≥8000 x g for 15 s at room temperature and the flow-through was discarded. This was repeated for the rest of the sample.

Bound RNA was washed 2 times using the included wash solution (Buffer RPE). Finally, elution solution (RNase-free water) was used to elute the RNA in 40 µL volume for cells and 20 µL for extracellular vesicles. RNA was stored at −80°C and RNA quality was assessed by Bioanalyzer (Agilent 2100 Bioanalyzer, RNA 6000 Nano Kit for cells, RNA 6000 Pico Kit for EVs, Agilent Technologies).

cDNA Libraries were prepared for small RNAs using the SMARTer smRNA-seq kit and for mRNAs using the SMART-Seq v4 and Nextera XT kit. A total of 18 cycles of PCR were carried out to obtain a good yield of cDNA from cells and EVs. Final library quality was verified with Qubit and Bioanalyzer. Negative (no RNA) and positive controls provided expected results. Next-generation RNA sequencing was performed using a HiSeq X Illumina 2x150, 40M total reads per sample (20M each direction) at Admera Health.

#### Data Analysis related to RNA seq studies of BV2 microglia and EVs: mRNA and non-coding RNA

RNA sequencing analysis was conducted to identify genes exhibiting differential expression. The counts data provided information on the abundance of reads mapped to the reference genome or the number of fragments assigned to each gene. The primary objective was to identify systematic changes between the conditions of interest, specifically Control vs Treatment. The analysis utilized the DESeq2 package, developed by Michael I. Love, Simon Anders, and Wolfgang Huber (Package Authors). A DESeq2DataSetFromMatrix object was created, and the design parameter was set to ∼Treatment, which captured the information on how the traits file was structured. For instance, in the case of n=4 Control vs n=4 LPS, the design matrix reflected these conditions. To ensure data quality, the raw counts data were aligned to the traits file, following the documentation guidelines. Lowly expressed genes were filtered out based on the criterion of rowSums(counts(dds)) ≥ 30, meaning that only genes with 30 reads in total across all samples (5 X 6 samples = 30) were retained. This step was performed to remove genes with insufficient expression levels. Additionally, if required by the dataset, Ensembl IDs were converted to gene names for ease of downstream analysis. The control condition was designated as the reference for comparison, and genes exhibiting differential expression were identified using the criteria of adjusted p-value < 0.05 and log2 fold change (LFC > 0 for upregulated genes and LFC < 0 for downregulated genes). It should be noted that in the comparison of Bulk RNA-seq on responder BV2 cells, the LPS samples were contrasted against the control (n = 4), deviating from the general protocol. Principal component analysis (PCA) (https://www.rdocumentation.org/packages/netresponse/versions/1.32.2/topics/plotPCA) was employed as an initial analysis step to identify patterns, relationships, and important variables in the high-dimensional dataset.

The RNA-seq analysis, utilizing the DESeq2 package, allowed for the identification of differentially expressed genes, subsequent gene enrichment analysis, and visualization of the results through various plots. This comprehensive approach provided valuable insights into the expression patterns and functional characteristics of the genes under investigation.

#### GSEA of transcriptomic data

Gene enrichment analysis was performed on the lists of differentially expressed genes using AltAnalyze, with specific parameters specified. Volcano plots (https://github.com/kevinblighe/EnhancedVolcano) and heatmaps (https://cran.r-project.org/web/packages/pheatmap/index.html) based on the top 50 genes with adjusted p-values were generated to visualize the results of the differential expression analysis. For gene enrichment analysis of the predicted targets for specific miRNAs, we used miRDB^35^ and MirTarget^54^. Input lists were relaxed to an adjusted p-value of 0.10, as opposed to the usual 0.05, in order to enhance the coverage of genes for Bulk RNA-seq on responder BV2 cells, and the LPS samples were contrasted against the control (n=4). To assess the statistical significance between two categorical variables, a Fisher’s exact test was conducted.

## Declarations

### Ethics approval and consent to participate

Not applicable.

### Consent for publication

Not applicable.

### Availability of data and materials

R code associated with the data analyses for the manuscript are published to GitHub (https://github.com/adityanatu1/proteomic-and-transcriptomic-signatures-of-microglia-derived-extracellular-vesicles-.git)

The mass spectrometry proteomics data have been deposited to the ProteomeXchange Consortium via the PRIDE partner repository with the dataset identifier PXD043959.

The RNA sequencing data have been deposited to the NCBI Gene Expression Omnibus (GEO). GEO accession number is pending.

Additional datasets supporting the conclusions of this article are included within the article as Additional files (13).

ExoCarta^55^ (http://www.exocarta.org) was used for lists of proteins previously identified in EVs from different mouse tissues and cell types.

Rangaraju et al. 2018^6^ was used for WGCNA module membership data used for analyses comparing the transcriptomes of responder BV2 microglia to known microglial transcriptomic profiles.

### Competing interests

The authors declare that they have no competing interests.

### Funding

Research in this publication is supported by the National institute of Aging of the National Institutes of Health and the National Institute of Neurological Disorders and Stroke of the National Institutes of Health: F31NS127530 (J. Santiago), R01AG075820 (S. Rangaraju), RF1AG071587 (S. Rangaraju), R01NS114130 (S. Rangaraju).

### Authors’ contributions

Conceptualization: J.V.S, S.Rayaprolu, N.T.S., S.Rangaraju.

Methodology: J.V.S., C.C.R., H.X., V.K, S.Rayaprolu, N.T.S., S.Rangaraju.

Investigation J.V.S., A.N., C.C.R., S.Rayaprolu, N.T.S., S.Rangaraju.

Writing-Original draft: J.V.S., A.N., S.Rangaraju.

Writing-Review and Editing: J.V.S., S.Rangaraju.

Funding acquisition: J.V.S, S.Rangaraju, N.T.S.

Resources: S.Rangaraju, N.T.S.

Supervision: S.Rangaraju

## Acknowledgements

Research reported in this publication was also supported in part by the Emory Integrated Proteomics Core (EIPC) and the Emory Robert P. Apkarian Integrated Electron Microscopy Core which are subsidized by the Emory University School of Medicine and Emory College of Arts and Sciences. Additional support was provided by the Biopolymer Characterization Core at Georgia Institute of Technology. Additional support was provided by the Georgia Clinical & Translational Science Alliance of the National Institutes of Health under award number UL1TR000454. The content is solely the responsibility of the authors and does not necessarily reflect the official views of the National Institutes of Health. Additionally, the TEM data described here was supported by the Georgia Clinical & Translational Science Alliance of the National Institutes of Health under award number UL1TR000454. Transmission electron micrographs were collected on the JEOL JEM-1400, 120kV TEM supported by the National Institutes of Health Grant S10 RR025679 or the Hitachi HT7700 120kV TEM supported by the Georgia Clinical and Translational Science Alliance under award number UL1TR002378. We would like to acknowledge Duc Duong (Emory University) for performing the LFQ-mass spectrometry on the samples, Admera for performing the RNA sequencing on the samples, Dr. Eric Dammer (Emory University) for helping us with data analyses, Samantha Reed (Emory University) for advice on extracellular vesicle isolation, and Dr. Navneet Dogra and Tina Chen (Icahn School of Medicine at Mount Sinai) for advice on RNA seq pipeline for extracellular vesicles. Lastly, I would like to acknowledge my dissertation committee members, Dr. Allan Levey, Dr. Nicholas Seyfried, Dr. Tamara Caspary, and Dr. Ellen Hess for the support they have provided over the years.

## Notes

### Competing Interest Statement

The authors have declared no competing interest.

