## Supplementary Figures for "Identification of state-specific proteomic and transcriptomic signatures of microglia-derived extracellular vesicles"

**SUPPLEMENTARY FIGURES (3)**

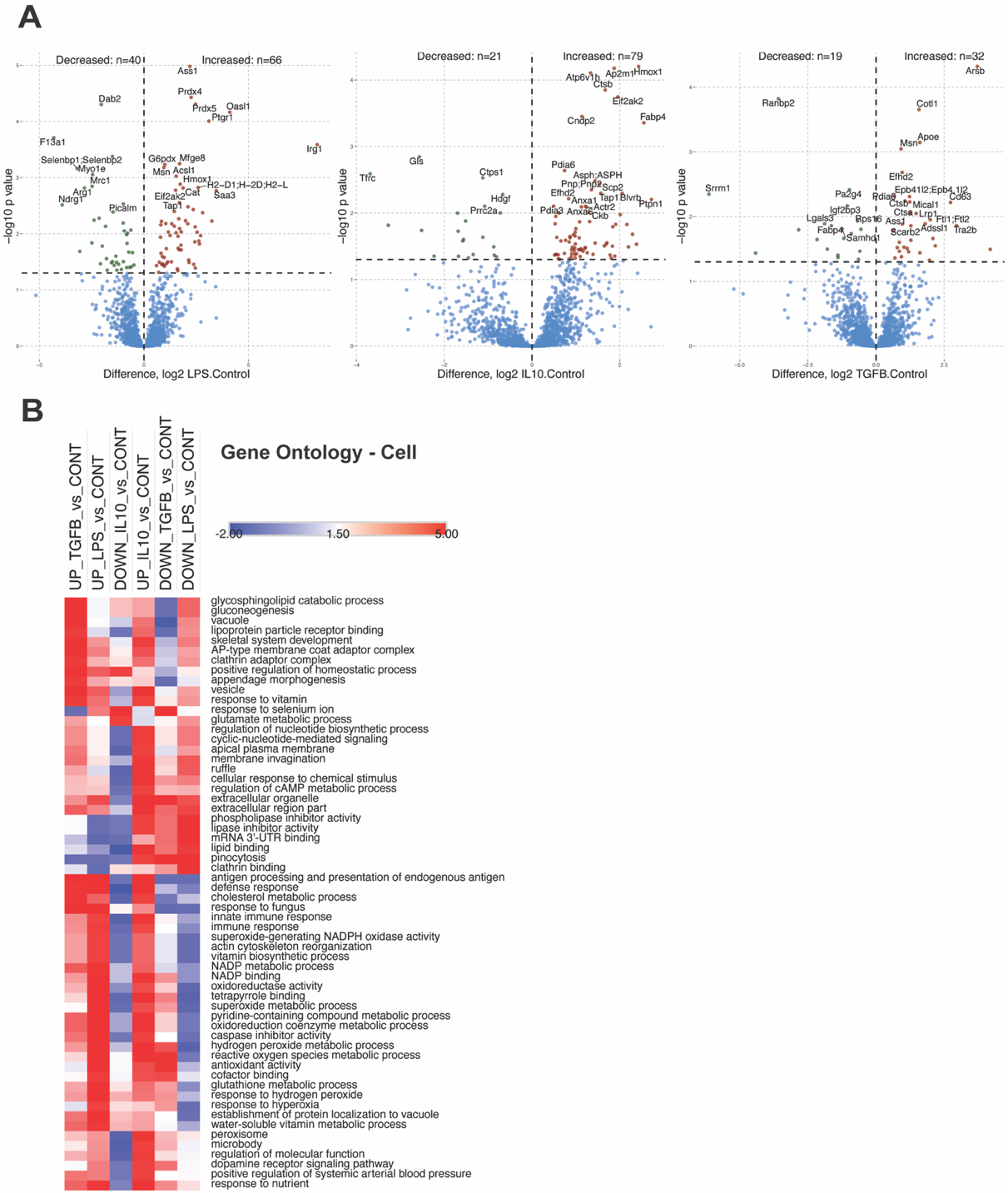

**Supplemental Figure 1: BV2 Cell Proteome. A.** Volcano plots showing differentially enriched proteins in EV proteome – IL10-CTL , LPS-CTL, TGF-β-CTL. **B.** Heatmap representation, based on enrichment Z-scores, of Gene Ontology for polarized cells (padj.<0.05).

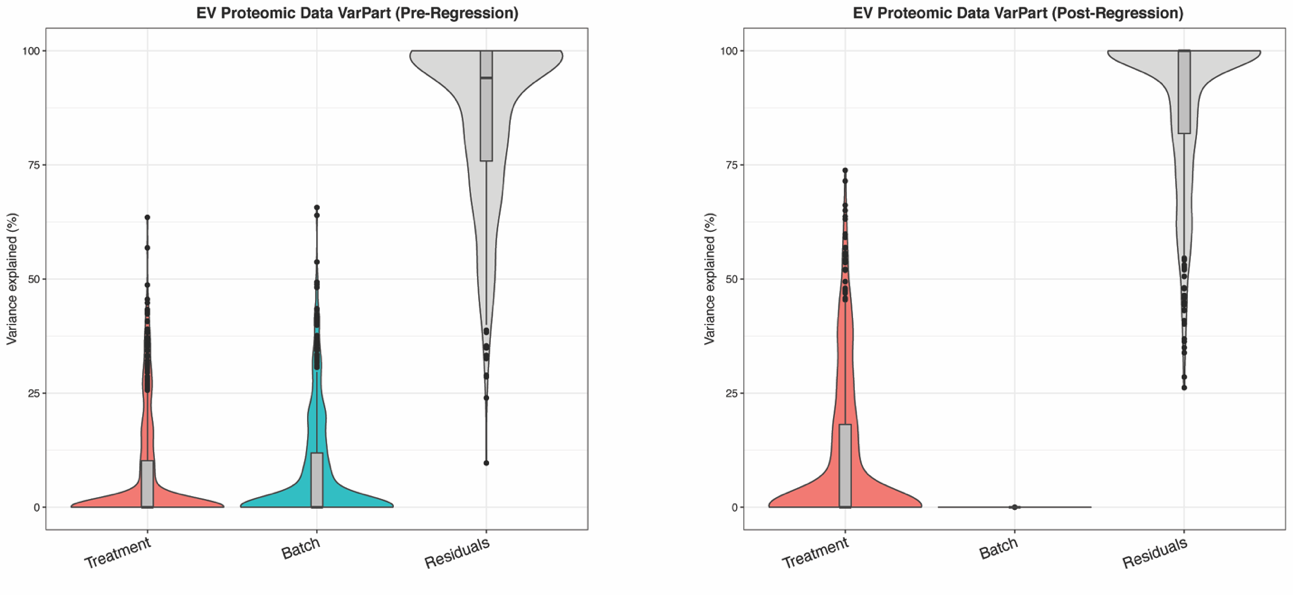

**Supplemental Figure 2: Pre and Post Regression Variance Partition Plots for BV2 EV Proteome**

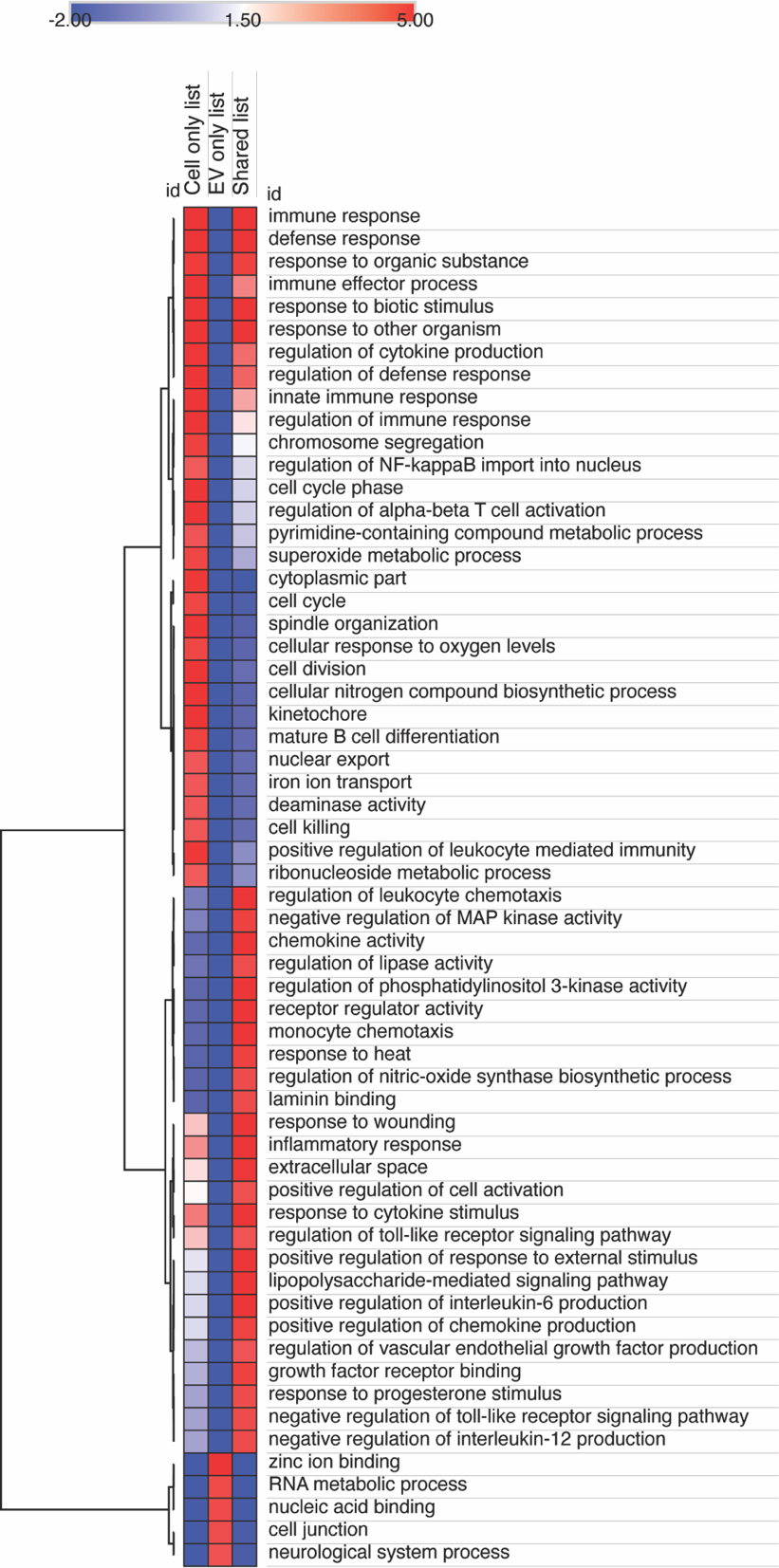

**Supplemental Figure 3:** Heatmap representation, based on enrichment Z-scores, of Gene Ontology from comparison of 2,949 upregulated mRNAs from BV2 cells treated with LPS to 1,040 upregulated mRNAs from LPS treated BV2-derived EVs (Figure 5F).
