## Additional File 5 for "Identification of state-specific proteomic and transcriptomic signatures of microglia-derived extracellular vesicles"

admera health

#### Sample ID Table & QC Results

| Sample # | Admera Health Sample ID | Customer Sample ID | Sample Type | Sample Volume (ul) | Admera Health Concentration (ng/ul) | Admera Health Total Quantity (ng) | RIN |
|---|---|---|---|---|---|---|---|
| 1 | 22082R-05-01 | CTL Cell (1) | RNA | 30 | 2800.0 | 84000 | 10.0 |
| 2 | 22082R-05-02 | CTL Cell (2) | RNA | 31 | 1690.0 | 52390 | 10.0 |
| 3 | 22082R-05-03 | CTL Cell (3) | RNA | 33 | 2380.0 | 78540 | 10.0 |
| 4 | 22082R-05-04 | LPS Cell (1) | RNA | 29 | 1130.0 | 32770 | 10.0 |
| 5 | 22082R-05-05 | LPS Cell (2) | RNA | 33 | 784.0 | 25872 | 10.0 |
| 6 | 22082R-05-06 | LPS Cell (3) | RNA | 33 | 1010.0 | 33330 | 10.0 |
| 7 | 22082R-05-07 | IL-10 Cell (1) | RNA | 32 | 1550.0 | 49600 | 10.0 |
| 8 | 22082R-05-08 | IL-10 Cell (2) | RNA | 32 | 1950.0 | 62400 | 10.0 |
| 9 | 22082R-05-09 | IL-10 Cell (3) | RNA | 33 | 1550.0 | 51150 | 10.0 |
| 10 | 22082R-05-10 | TGFB Cell (1) | RNA | 33 | 1680.0 | 55440 | 10.0 |
| 11 | 22082R-05-11 | TGFB Cell (2) | RNA | 33 | 2160.0 | 71280 | 10.0 |
| 12 | 22082R-05-12 | TGFB Cell (3) | RNA | 31 | 2160.0 | 66960 | 10.0 |
| 13 | 22082R-05-13 | CTL EV (1) | RNA | 18 | 0.1 | 2 | N/A |
| 14 | 22082R-05-14 | CTL EV (2) | RNA | 31 | 0.1 | 4 | N/A |
| 15 | 22082R-05-15 | CTL EV (3) | RNA | 18 | 0.2 | 4 | N/A |
| 16 | 22082R-05-16 | LPS EV (1) | RNA | 18 | 0.2 | 4 | N/A |
| 17 | 22082R-05-17 | LPS EV (2) | RNA | 18 | 0.9 | 16 | 2.0 |
| 18 | 22082R-05-18 | LPS EV (3) | RNA | 18 | 0.2 | 3 | N/A |
| 19 | 22082R-05-19 | IL-10 EV (1) | RNA | 18 | 0.1 | 2 | N/A |
| 20 | 22082R-05-20 | IL-10 EV (2) | RNA | 18 | 0.4 | 7 | N/A |
| 21 | 22082R-05-21 | IL-10 EV (3) | RNA | 18 | 0.3 | 6 | N/A |
| 22 | 22082R-05-22 | TGFB EV (1) | RNA | 18 | 0.3 | 5 | N/A |
| 23 | 22082R-05-23 | TGFB EV (2) | RNA | 18 | 0.3 | 5 | N/A |
| 24 | 22082R-05-24 | TGFB EV (3) | RNA | 18 | 0.2 | 4 | N/A |

### TapeStation Image

#### Filename: 22082-05-01-12-IQC-TS-01252023.cRNA

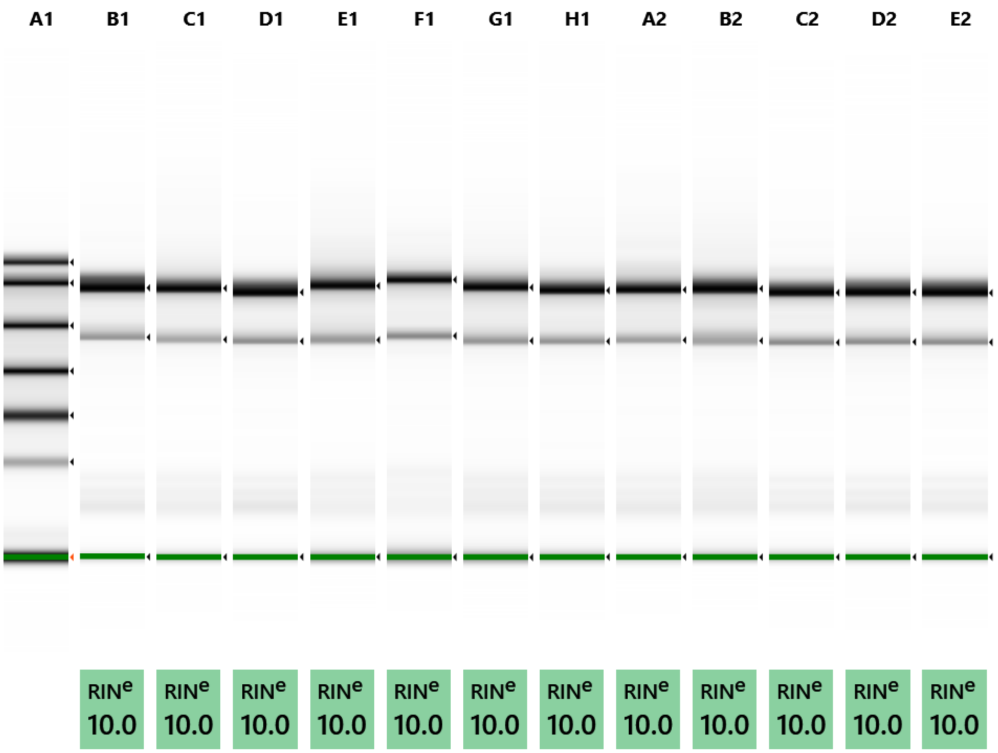


**Default image (Contrast 50%), Image is Scaled to Sample**

#### *Sample Info*

| Well | $RIN^e$ | 28S/18S (Area) | Conc. [ng/µl] | Sample Description | Alert | Observations |
|---|---|---|---|---|---|---|
| A1 | - | - | 84.9 | Electronic Ladder | | Ladder |
| B1 | 10.0 | 2.4 | 318 | 22082R-05-01 | | |
| C1 | 10.0 | 3.2 | 199 | 22082R-05-02 | | |
| D1 | 10.0 | 2.0 | 229 | 22082R-05-03 | | |
| E1 | 10.0 | 2.7 | 118 | 22082R-05-04 | | |
| F1 | 10.0 | 2.5 | 74.2 | 22082R-05-05 | | |
| G1 | 10.0 | 2.6 | 108 | 22082R-05-06 | | |
| H1 | 10.0 | 3.0 | 157 | 22082R-05-07 | | |
| A2 | 10.0 | 3.1 | 233 | 22082R-05-08 | | |
| B2 | 10.0 | 2.7 | 176 | 22082R-05-09 | | |
| C2 | 10.0 | 3.1 | 179 | 22082R-05-10 | | |
| D2 | 10.0 | 3.2 | 228 | 22082R-05-11 | | |
| E2 | 10.0 | 3.3 | 238 | 22082R-05-12 | | |

#### A1: Electronic Ladder

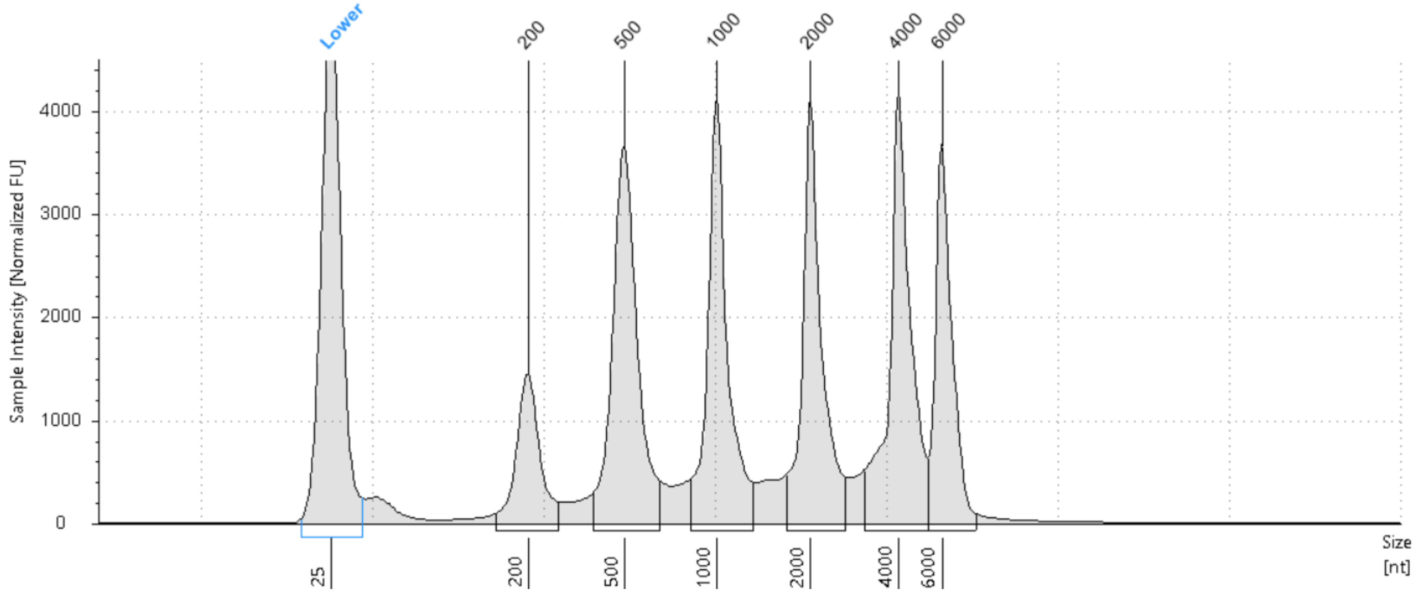


#### Sample Table

| Well | RIN$^{e}$ | 28S/18S (Area) | Conc. [ng/µl] | Sample Description | Alert | Observations |
|---|---|---|---|---|---|---|
| A1 | - | - | 84.9 | Electronic Ladder | | Ladder |

#### Peak Table

| Size [nt] | Calibrated Conc. [ng/µl] | Assigned Conc. [ng/µl] | Peak Molarity [nmol/l] | % Integrated Area | Peak Comment | Observations |
|---|---|---|---|---|---|---|
| 25 | 40.0 | 40.0 | 4710 | - | | Lower Marker |
| 200 | 5.94 | - | 87.4 | 7.80 | | |
| 500 | 15.9 | - | 93.6 | 20.88 | | |
| 1000 | 14.2 | - | 41.8 | 18.62 | | |
| 2000 | 13.8 | - | 20.3 | 18.11 | | |
| 4000 | 15.5 | - | 11.4 | 20.38 | | |
| 6000 | 10.8 | - | 5.31 | 14.22 | | |

*B1: 22082R-05-01*

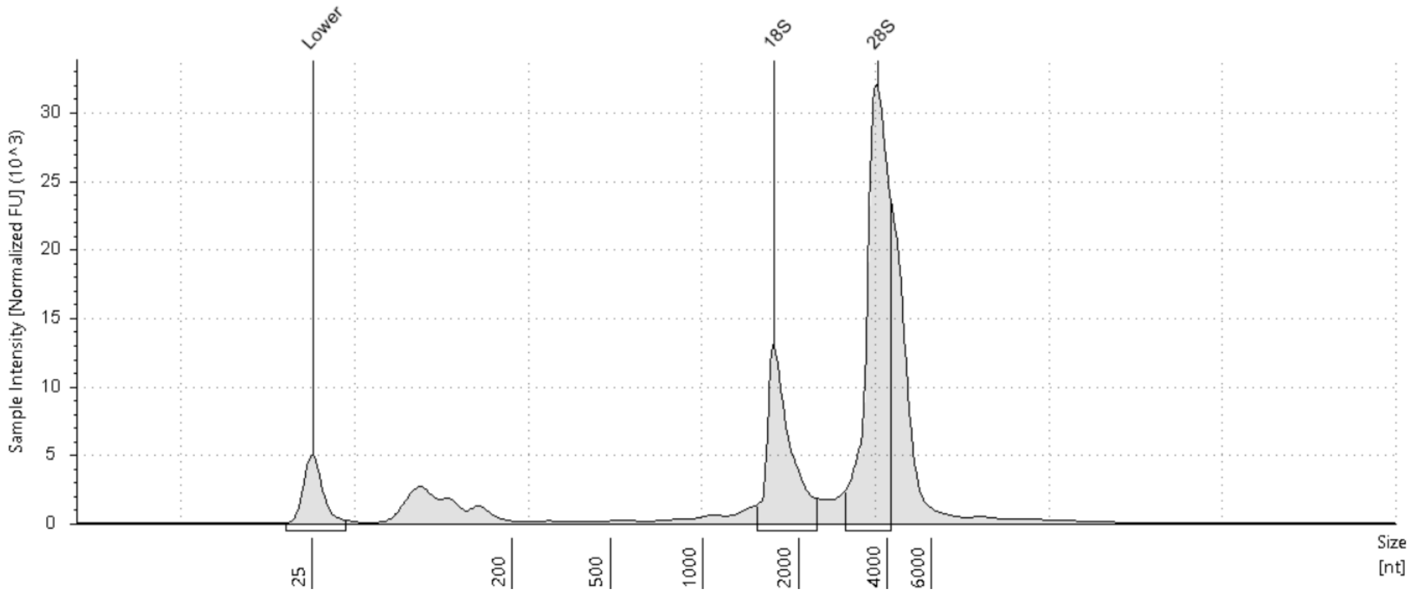


#### Sample Table

| Well | $RIN^e$ | 28S/18S (Area) | Conc. [ng/µl] | Sample Description | Alert | Observations |
|---|---|---|---|---|---|---|
| B1 | 10.0 | 2.4 | 318 | 22082R-05-01 | | |

#### Peak Table

| Size [nt] | Calibrated Conc. [ng/µl] | Assigned Conc. [ng/µl] | Peak Molarity [nmol/l] | % Integrated Area | Peak Comment | Observations |
|---|---|---|---|---|---|---|
| 25 | 40.0 | 40.0 | 4710 | - | | Lower Marker |
| 1668 | 57.2 | - | 101 | 29.80 | | 18S |
| 3683 | 135 | - | 108 | 70.20 | | 28S |

*C1: 22082R-05-02*

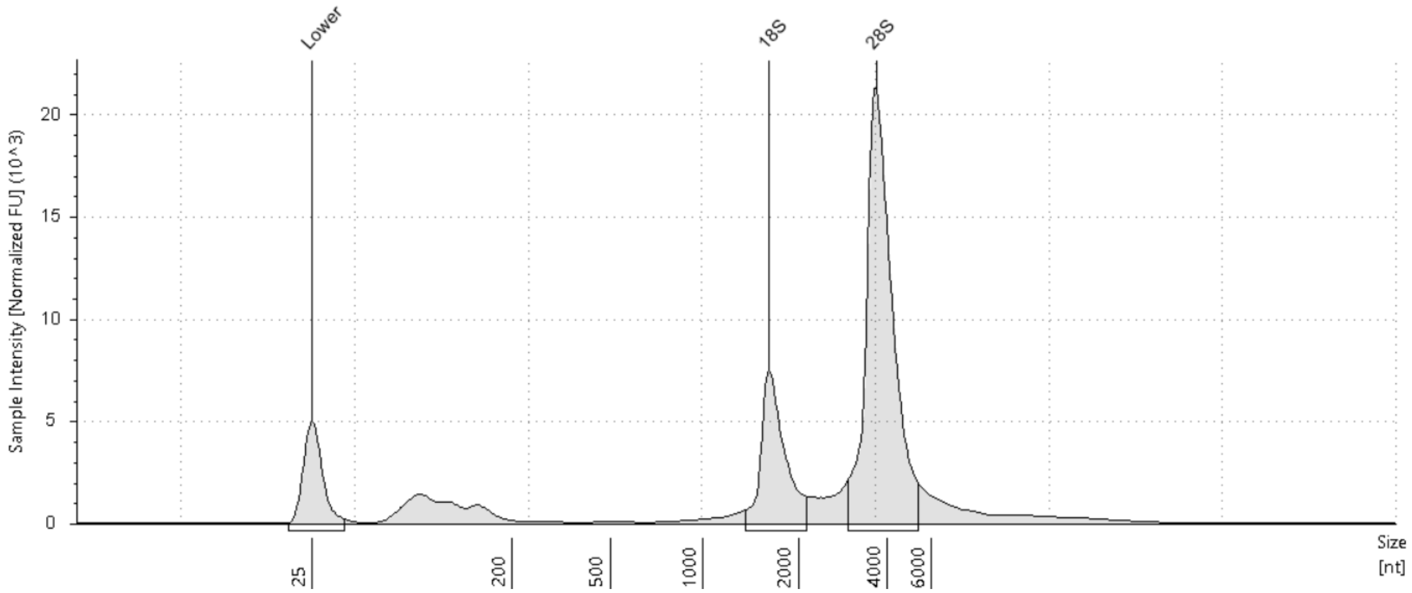


#### Sample Table

| Well | RIN[e] | 28S/18S (Area) | Conc. [ng/µl] | Sample Description | Alert | Observations |
|---|---|---|---|---|---|---|
| C1 | 10.0 | 3.2 | 199 | 22082R-05-02 | | |

#### Peak Table

| Size [nt] | Calibrated Conc. [ng/µl] | Assigned Conc. [ng/µl] | Peak Molarity [nmol/l] | % Integrated Area | Peak Comment | Observations |
|---|---|---|---|---|---|---|
| 25 | 40.0 | 40.0 | 4710 | - | | Lower Marker |
| 1616 | 34.9 | - | 63.5 | 23.98 | | 18S |
| 3671 | 111 | - | 88.6 | 76.02 | | 28S |

*D1: 22082R-05-03*

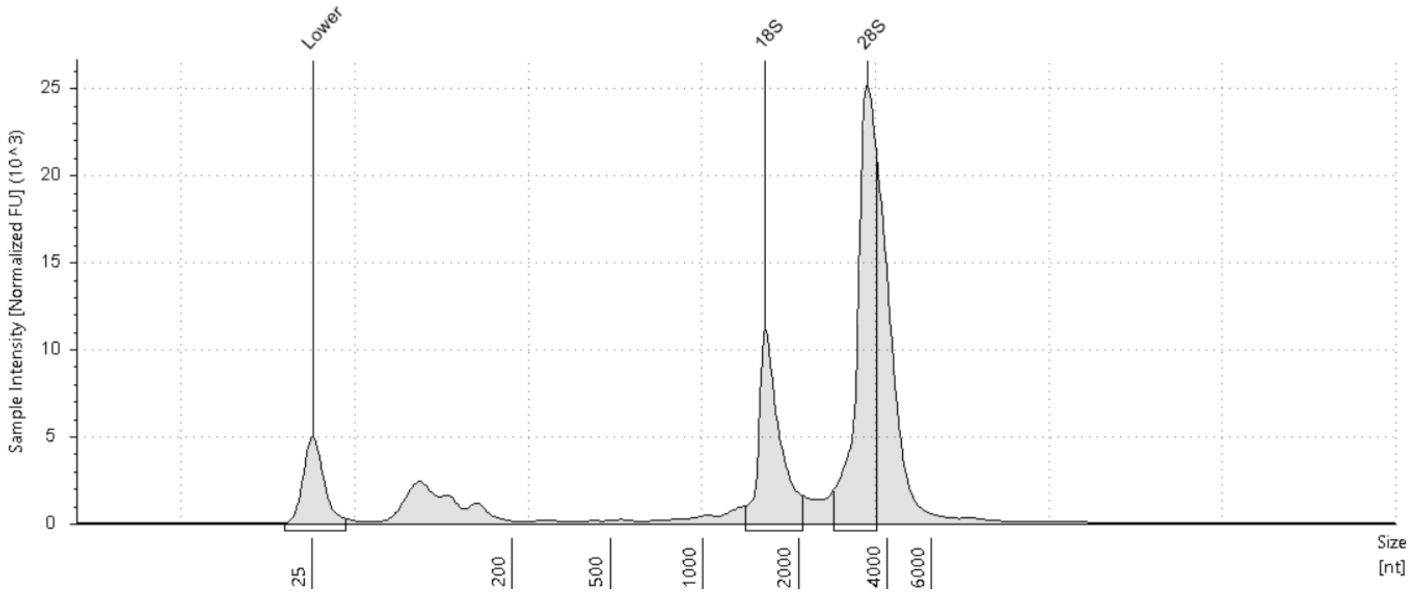


#### Sample Table

| Well | RIN$^e$ | 28S/18S (Area) | Conc. [ng/µl] | Sample Description | Alert | Observations |
|---|---|---|---|---|---|---|
| D1 | 10.0 | 2.0 | 229 | 22082R-05-03 | | |

#### Peak Table

| Size [nt] | Calibrated Conc. [ng/µl] | Assigned Conc. [ng/µl] | Peak Molarity [nmol/l] | % Integrated Area | Peak Comment | Observations |
|---|---|---|---|---|---|---|
| 25 | 40.0 | 40.0 | 4710 | - | | Lower Marker |
| 1570 | 42.3 | - | 79.3 | 33.29 | | 18S |
| 3424 | 84.8 | - | 72.9 | 66.71 | | 28S |

*E1: 22082R-05-04*

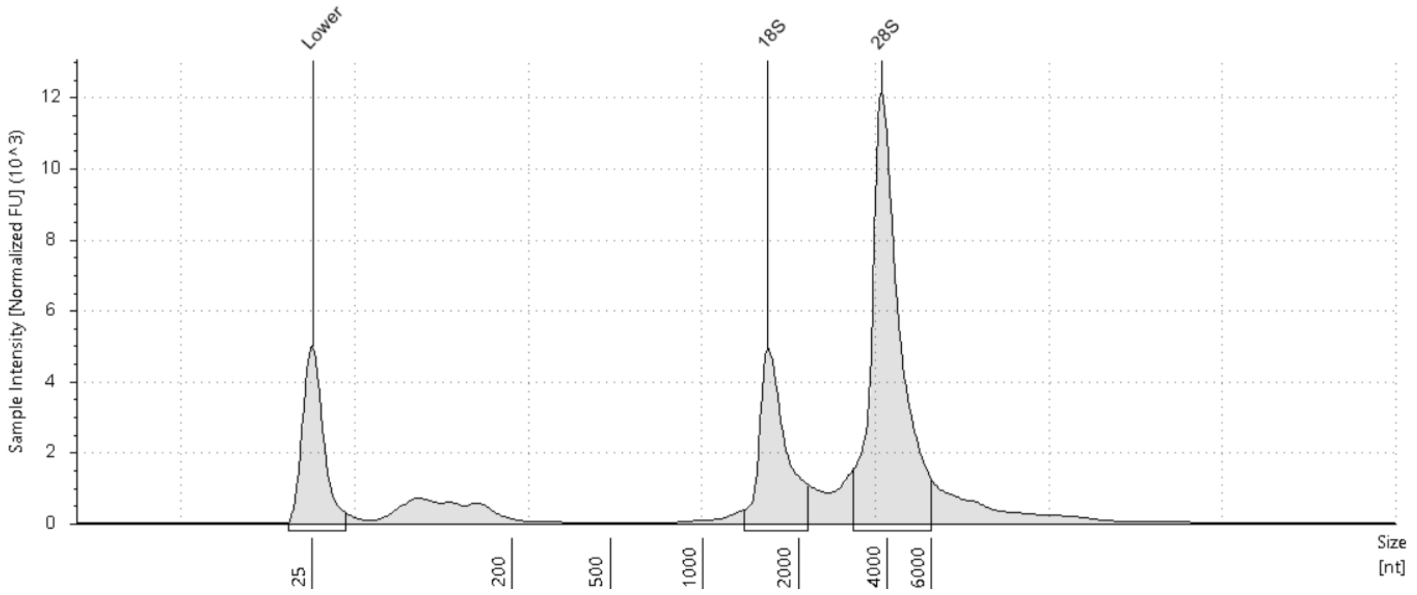


#### Sample Table

| Well | RIN$^e$ | 28S/18S (Area) | Conc. [ng/µl] | Sample Description | Alert | Observations |
|---|---|---|---|---|---|---|
| E1 | 10.0 | 2.7 | 118 | 22082R-05-04 | | |

#### Peak Table

| Size [nt] | Calibrated Conc. [ng/µl] | Assigned Conc. [ng/µl] | Peak Molarity [nmol/l] | % Integrated Area | Peak Comment | Observations |
|---|---|---|---|---|---|---|
| 25 | 40.0 | 40.0 | 4710 | - | | Lower Marker |
| 1606 | 23.3 | - | 42.7 | 27.33 | | 18S |
| 3822 | 62.0 | - | 47.7 | 72.67 | | 28S |

*F1: 22082R-05-05*

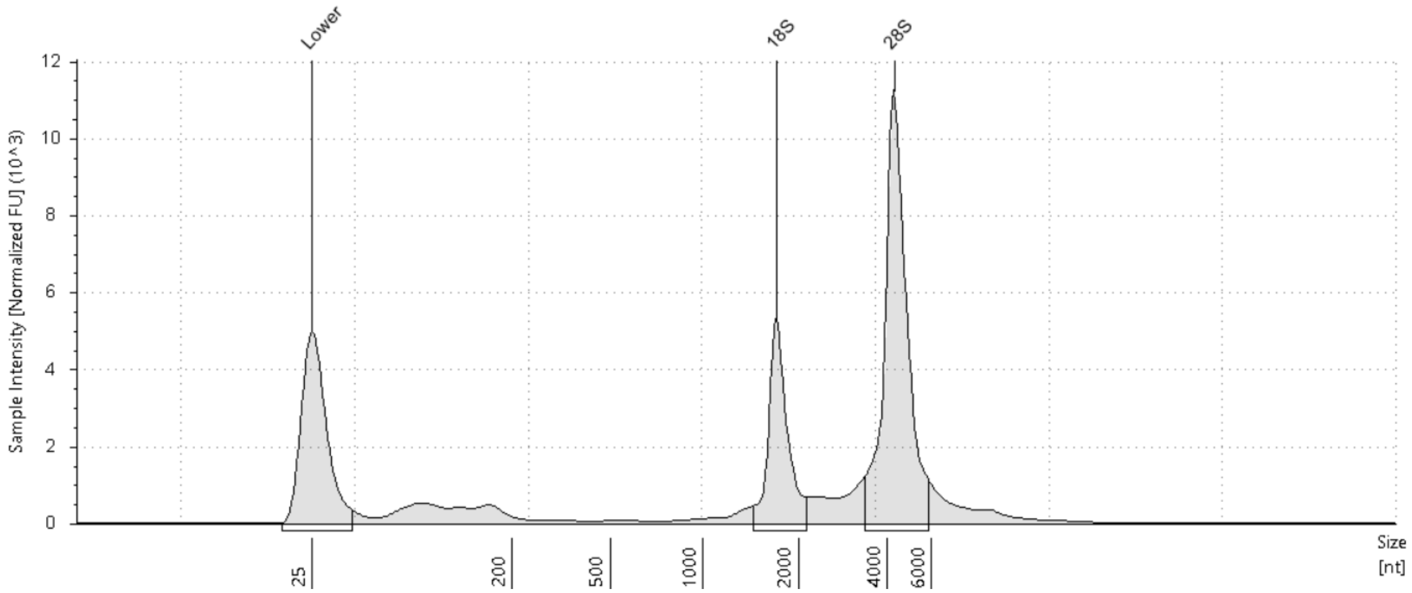


#### Sample Table

| Well | RIN^e | 28S/18S (Area) | Conc. [ng/µl] | Sample Description | Alert | Observations |
|---|---|---|---|---|---|---|
| F1 | 10.0 | 2.5 | 74.2 | 22082R-05-05 | | |

#### Peak Table

| Size [nt] | Calibrated Conc. [ng/µl] | Assigned Conc. [ng/µl] | Peak Molarity [nmol/l] | % Integrated Area | Peak Comment | Observations |
|---|---|---|---|---|---|---|
| 25 | 40.0 | 40.0 | 4710 | - | | Lower Marker |
| 1711 | 14.7 | - | 25.4 | 28.55 | | 18S |
| 4265 | 36.9 | - | 25.5 | 71.45 | | 28S |

*G1: 22082R-05-06*

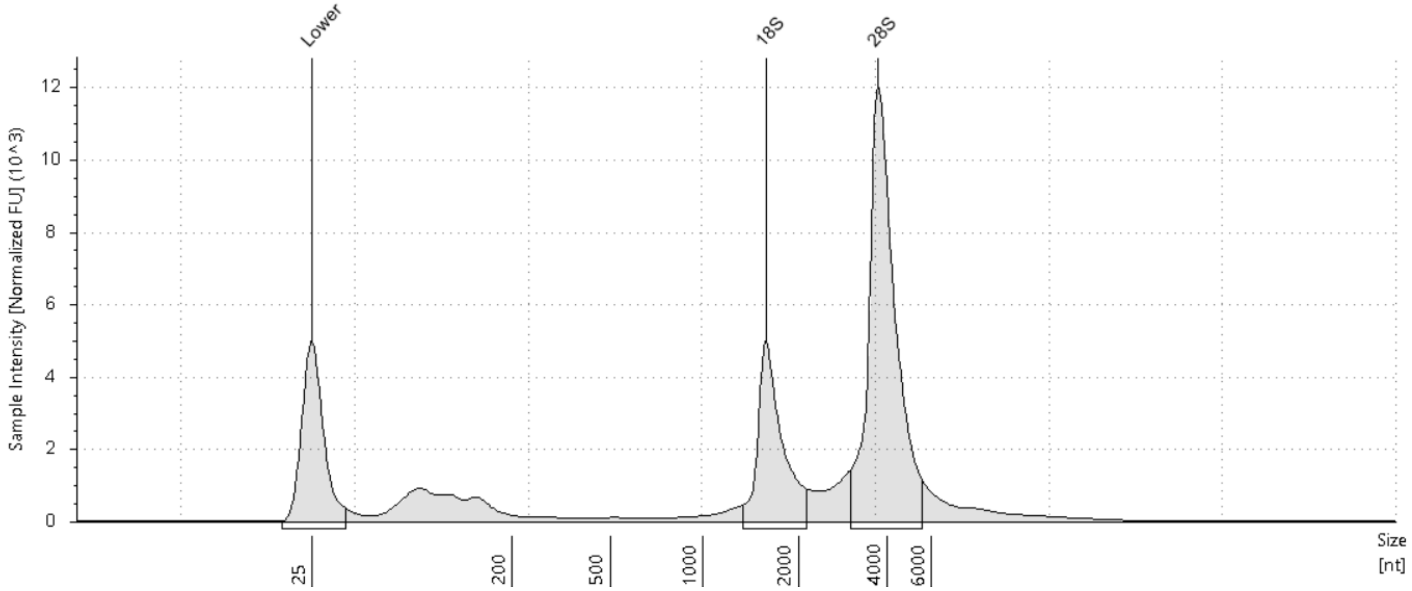


#### Sample Table

| Well | RIN$^e$ | 28S/18S (Area) | Conc. [ng/µl] | Sample Description | Alert | Observations |
|---|---|---|---|---|---|---|
| G1 | 10.0 | 2.6 | 108 | 22082R-05-06 | | |

#### Peak Table

| Size [nt] | Calibrated Conc. [ng/µl] | Assigned Conc. [ng/µl] | Peak Molarity [nmol/l] | % Integrated Area | Peak Comment | Observations |
|---|---|---|---|---|---|---|
| 25 | 40.0 | 40.0 | 4710 | - | | Lower Marker |
| 1590 | 21.0 | - | 38.9 | 27.53 | | 18S |
| 3730 | 55.4 | - | 43.6 | 72.47 | | 28S |

*H1: 22082R-05-07*

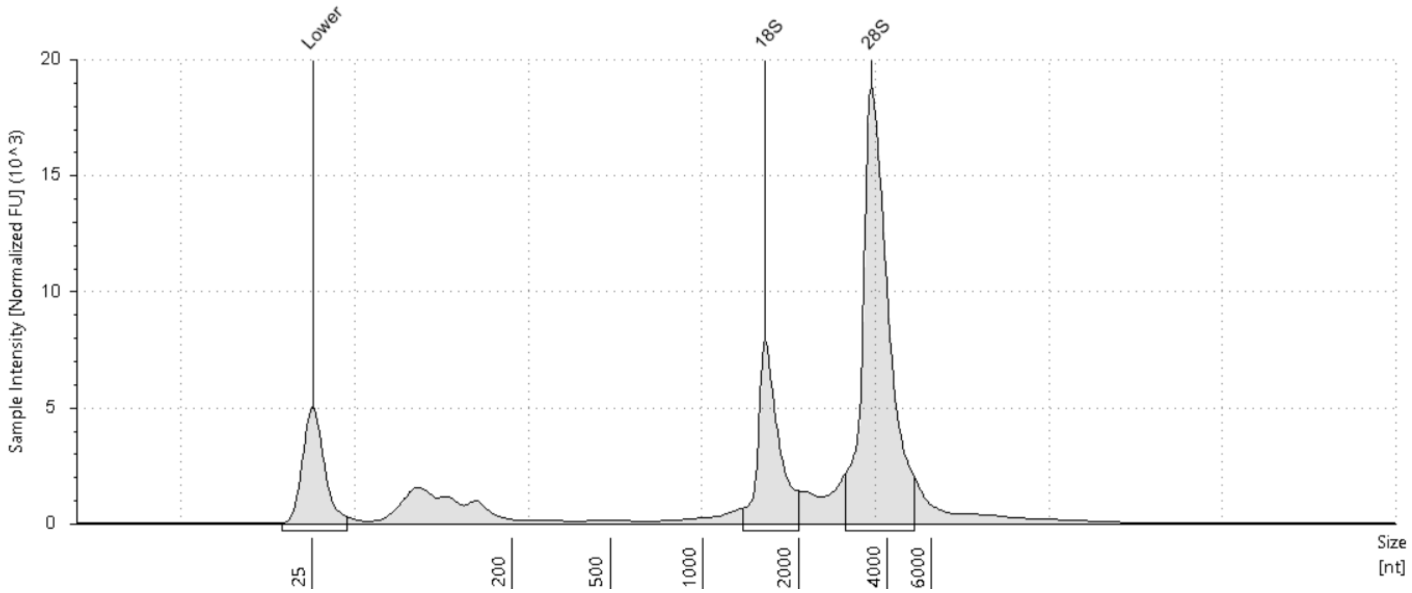


#### Sample Table

| Well | RIN^e | 28S/18S (Area) | Conc. [ng/µl] | Sample Description | Alert | Observations |
|---|---|---|---|---|---|---|
| H1 | 10.0 | 3.0 | 157 | 22082R-05-07 | | |

#### Peak Table

| Size [nt] | Calibrated Conc. [ng/µl] | Assigned Conc. [ng/µl] | Peak Molarity [nmol/l] | % Integrated Area | Peak Comment | Observations |
|---|---|---|---|---|---|---|
| 25 | 40.0 | 40.0 | 4710 | - | | Lower Marker |
| 1571 | 27.9 | - | 52.2 | 24.94 | | 18S |
| 3536 | 84.0 | - | 69.9 | 75.06 | | 28S |

*A2: 22082R-05-08*

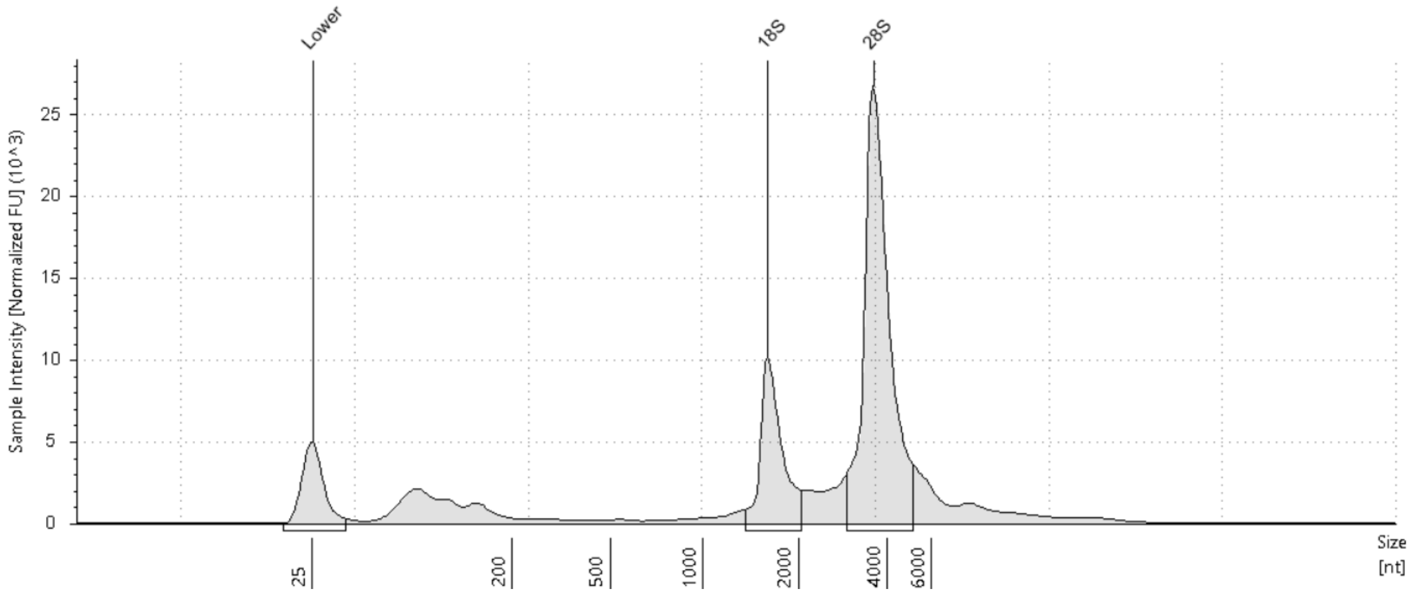


#### Sample Table

| Well | RIN[e] | 28S/18S (Area) | Conc. [ng/µl] | Sample Description | Alert | Observations |
|---|---|---|---|---|---|---|
| A2 | 10.0 | 3.1 | 233 | 22082R-05-08 | | |

#### Peak Table

| Size [nt] | Calibrated Conc. [ng/µl] | Assigned Conc. [ng/µl] | Peak Molarity [nmol/l] | % Integrated Area | Peak Comment | Observations |
|---|---|---|---|---|---|---|
| 25 | 40.0 | 40.0 | 4710 | - | | Lower Marker |
| 1600 | 38.1 | - | 70.0 | 24.47 | | 18S |
| 3575 | 118 | - | 96.7 | 75.53 | | 28S |

*B2: 22082R-05-09*

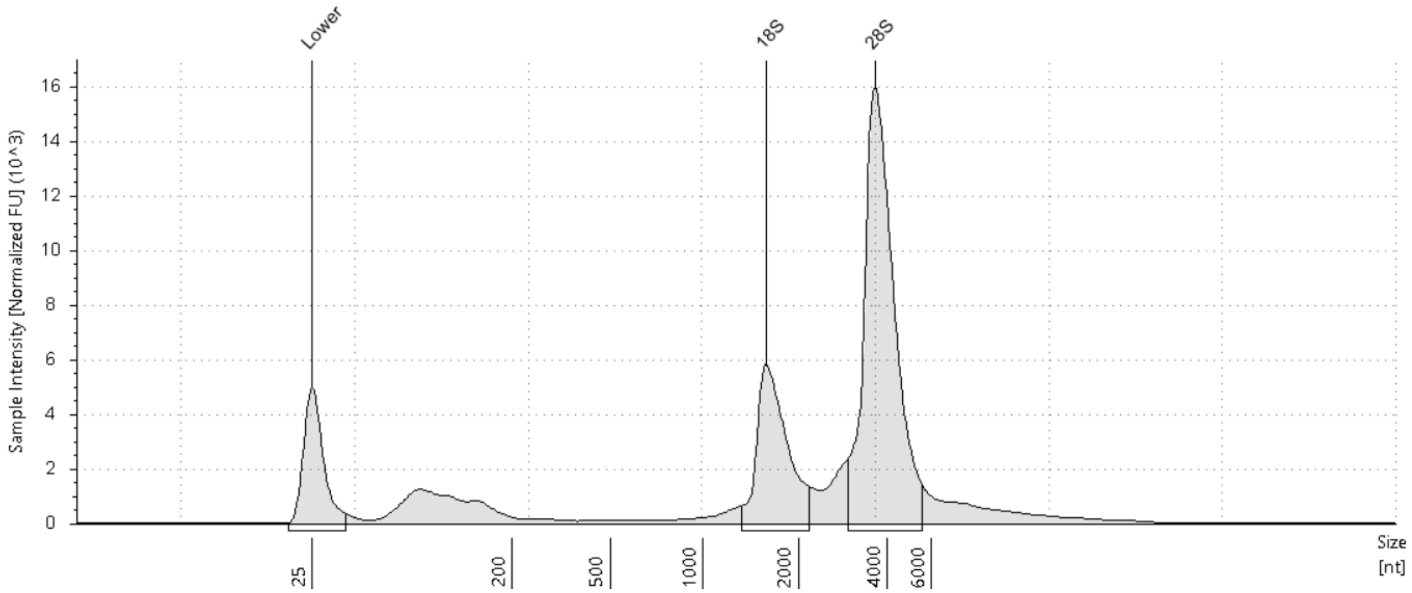


#### Sample Table

| Well | RIN[e] | 28S/18S (Area) | Conc. [ng/µl] | Sample Description | Alert | Observations |
|---|---|---|---|---|---|---|
| B2 | 10.0 | 2.7 | 176 | 22082R-05-09 | | |

#### Peak Table

| Size [nt] | Calibrated Conc. [ng/µl] | Assigned Conc. [ng/µl] | Peak Molarity [nmol/l] | % Integrated Area | Peak Comment | Observations |
|---|---|---|---|---|---|---|
| 25 | 40.0 | 40.0 | 4710 | - | | Lower Marker |
| 1590 | 33.6 | - | 62.1 | 26.72 | | 18S |
| 3659 | 92.2 | - | 74.1 | 73.28 | | 28S |

*C2: 22082R-05-10*

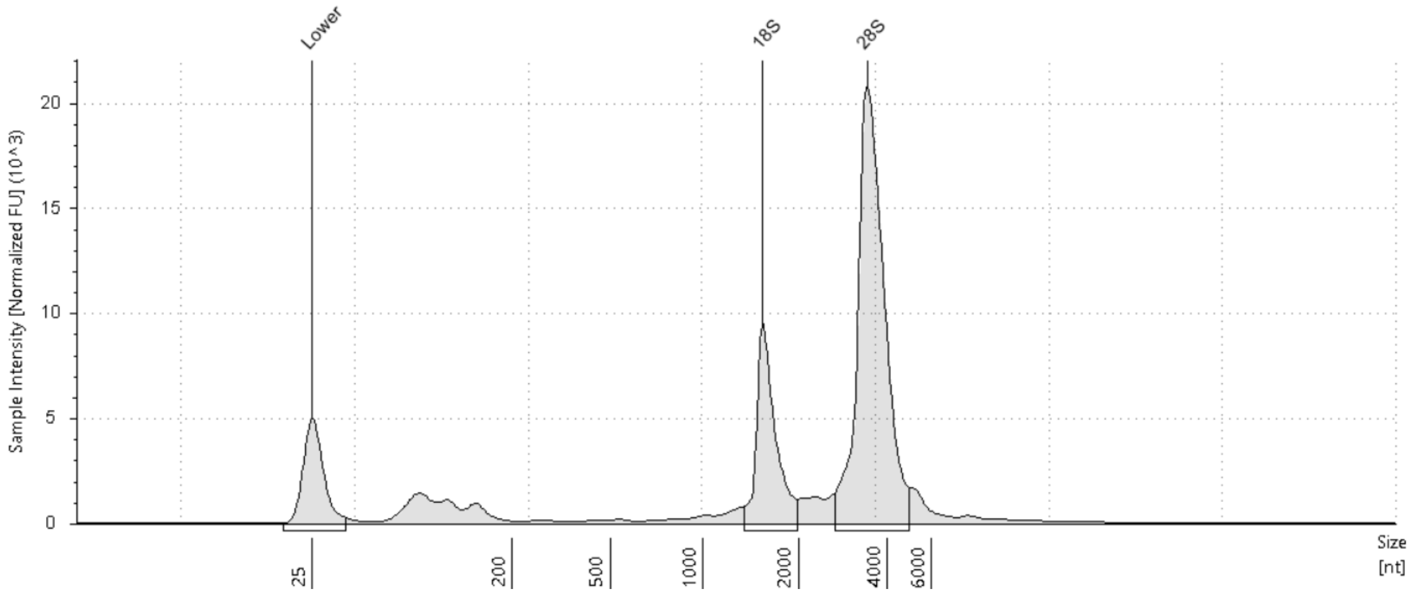


#### Sample Table

| Well | RIN^e | 28S/18S (Area) | Conc. [ng/µl] | Sample Description | Alert | Observations |
|---|---|---|---|---|---|---|
| C2 | 10.0 | 3.1 | 179 | 22082R-05-10 | | |

#### Peak Table

| Size [nt] | Calibrated Conc. [ng/µl] | Assigned Conc. [ng/µl] | Peak Molarity [nmol/l] | % Integrated Area | Peak Comment | Observations |
|---|---|---|---|---|---|---|
| 25 | 40.0 | 40.0 | 4710 | - | | Lower Marker |
| 1545 | 33.0 | - | 62.8 | 24.14 | | 18S |
| 3417 | 104 | - | 89.2 | 75.86 | | 28S |

*D2: 22082R-05-11*

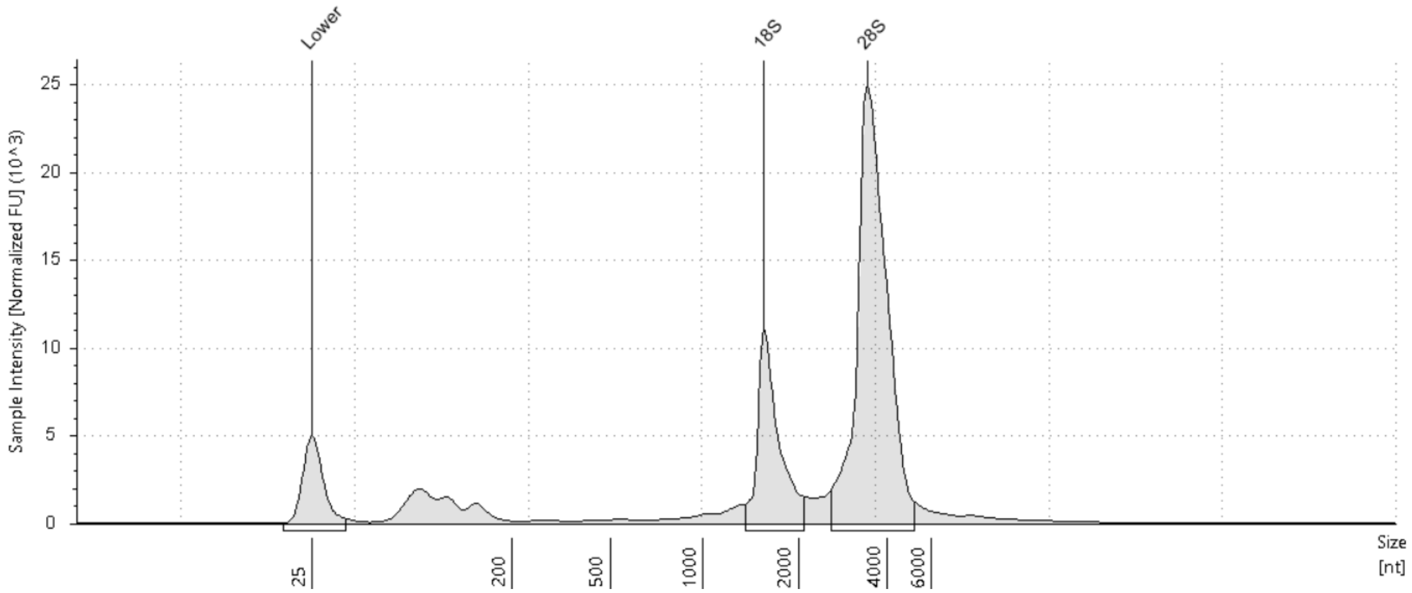


#### Sample Table

| Well | RIN$^{e}$ | 28S/18S (Area) | Conc. [ng/µl] | Sample Description | Alert | Observations |
|---|---|---|---|---|---|---|
| D2 | 10.0 | 3.2 | 228 | 22082R-05-11 | | |

#### Peak Table

| Size [nt] | Calibrated Conc. [ng/µl] | Assigned Conc. [ng/µl] | Peak Molarity [nmol/l] | % Integrated Area | Peak Comment | Observations |
|---|---|---|---|---|---|---|
| 25 | 40.0 | 40.0 | 4710 | - | | Lower Marker |
| 1562 | 42.6 | - | 80.2 | 23.80 | | 18S |
| 3442 | 136 | - | 116 | 76.20 | | 28S |

*E2: 22082R-05-12*

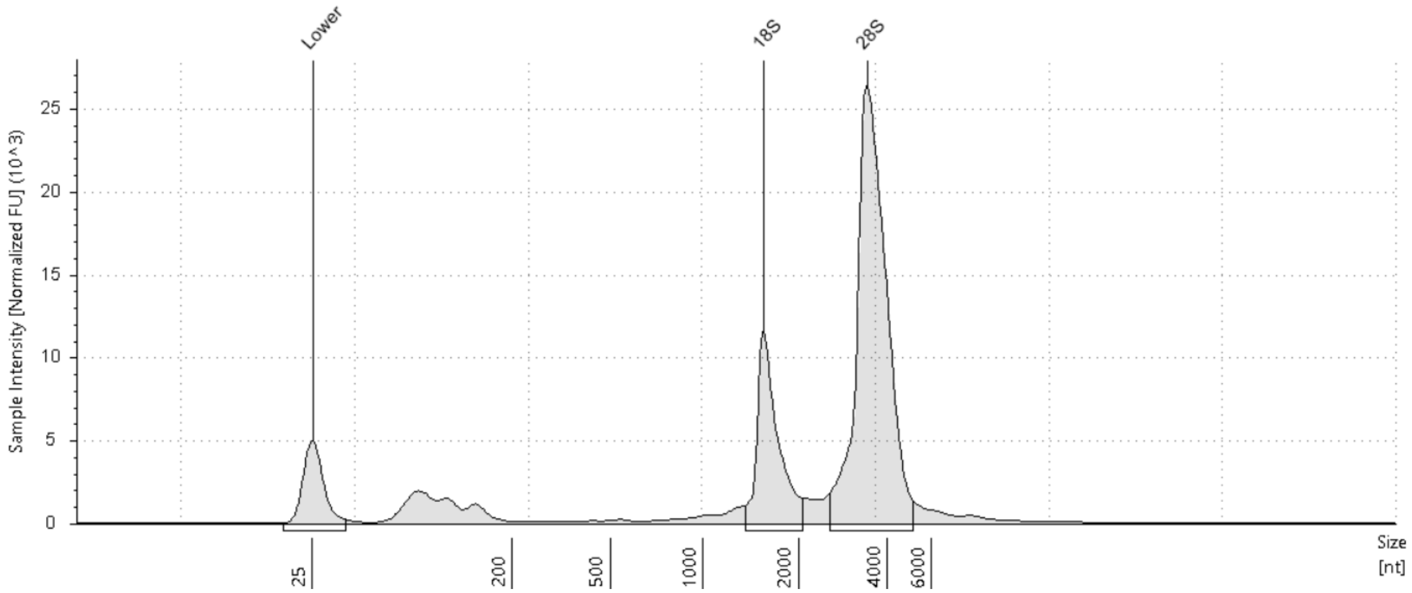


#### Sample Table

| Well | RIN^e | 28S/18S (Area) | Conc. [ng/µl] | Sample Description | Alert | Observations |
|---|---|---|---|---|---|---|
| E2 | 10.0 | 3.3 | 238 | 22082R-05-12 | | |

#### Peak Table

| Size [nt] | Calibrated Conc. [ng/µl] | Assigned Conc. [ng/µl] | Peak Molarity [nmol/l] | % Integrated Area | Peak Comment | Observations |
|---|---|---|---|---|---|---|
| 25 | 40.0 | 40.0 | 4710 | - | | Lower Marker |
| 1550 | 44.5 | - | 84.5 | 23.44 | | 18S |
| 3401 | 146 | - | 126 | 76.56 | | 28S |

#### Filename: 22082-05-13-24-IQC-TS-01252023.cHSRNA

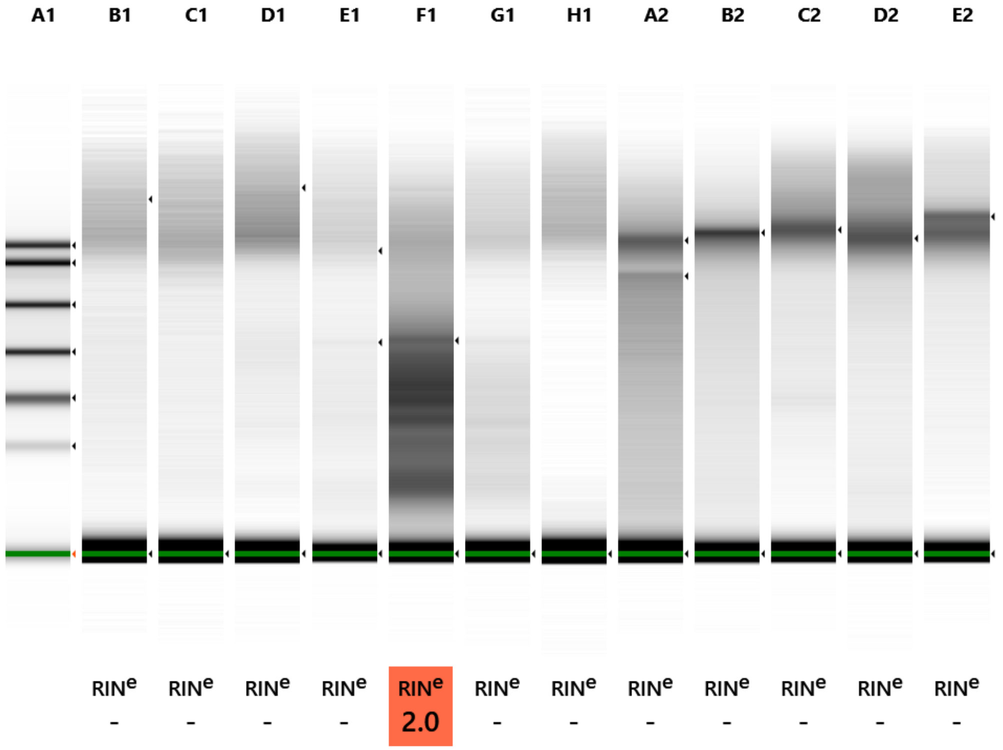


**Default image (Contrast 50%), Image is Scaled to Sample**

##### *Sample Info*

| Well | RINe | 28S/18S (Area) | Conc. [pg/µl] | Sample Description | Alert | Observations |
|---|---|---|---|---|---|---|
| A1 | - | - | 3750 | Electronic Ladder | | Ladder |
| B1 | - | - | 135 | 22082R-05-13 | | |
| C1 | - | - | 129 | 22082R-05-14 | | |
| D1 | - | - | 198 | 22082R-05-15 | | |
| E1 | - | - | 212 | 22082R-05-16 | | |
| F1 | 2.0 | - | 906 | 22082R-05-17 | | |
| G1 | - | - | 179 | 22082R-05-18 | | |
| H1 | - | - | 87.4 | 22082R-05-19 | | |
| A2 | - | - | 373 | 22082R-05-20 | | |
| B2 | - | - | 321 | 22082R-05-21 | | |
| C2 | - | - | 282 | 22082R-05-22 | | |
| D2 | - | - | 281 | 22082R-05-23 | | |
| E2 | - | - | 238 | 22082R-05-24 | | |

#### *A1: Electronic Ladder*

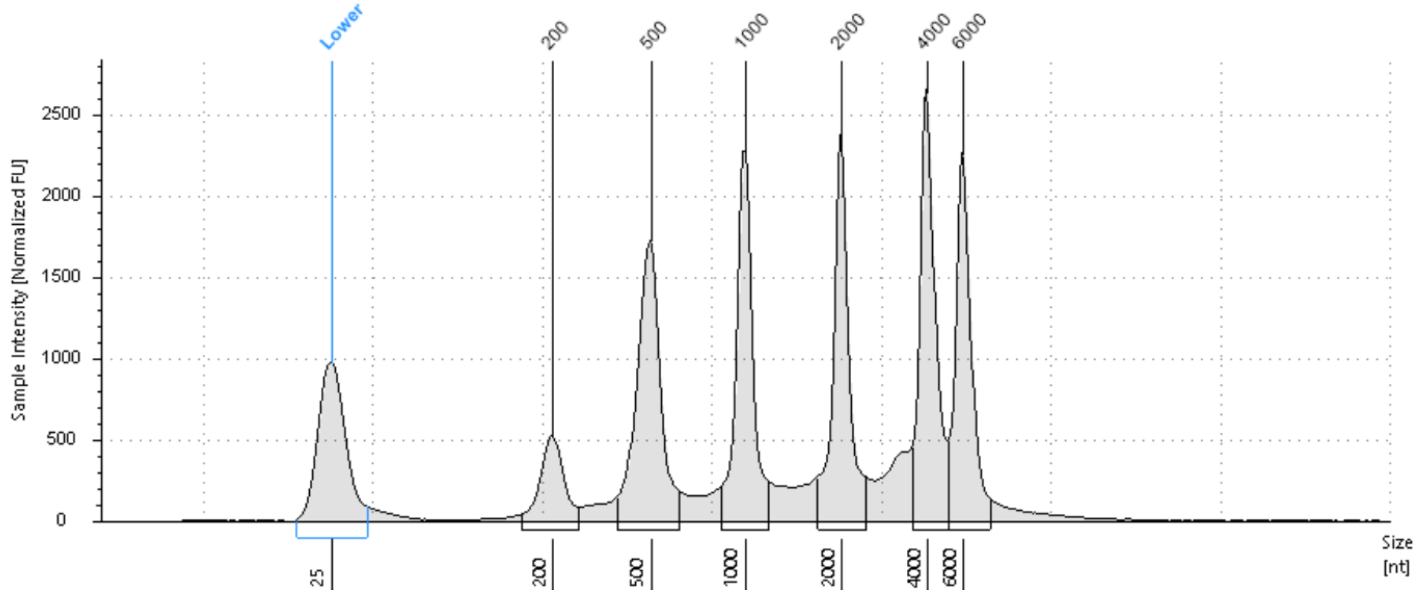


#### Sample Table

| Well | RIN$^e$ | 28S/18S (Area) | Conc. [pg/µl] | Sample Description | Alert | Observations |
|---|---|---|---|---|---|---|
| A1 | - | - | 3750 | Electronic Ladder | | Ladder |

#### Peak Table

| Size [nt] | Calibrated Conc. [pg/µl] | Assigned Conc. [pg/µl] | Peak Molarity [pmol/l] | % Integrated Area | Peak Comment | Observations |
|---|---|---|---|---|---|---|
| 25 | 700 | 700 | 82400 | - | | Lower Marker |
| 200 | 189 | - | 2770 | 5.98 | | |
| 500 | 638 | - | 3760 | 20.25 | | |
| 1000 | 557 | - | 1640 | 17.66 | | |
| 2000 | 580 | - | 854 | 18.42 | | |
| 4000 | 634 | - | 466 | 20.11 | | |
| 6000 | 554 | - | 272 | 17.58 | | |

*B1: 22082R-05-13*

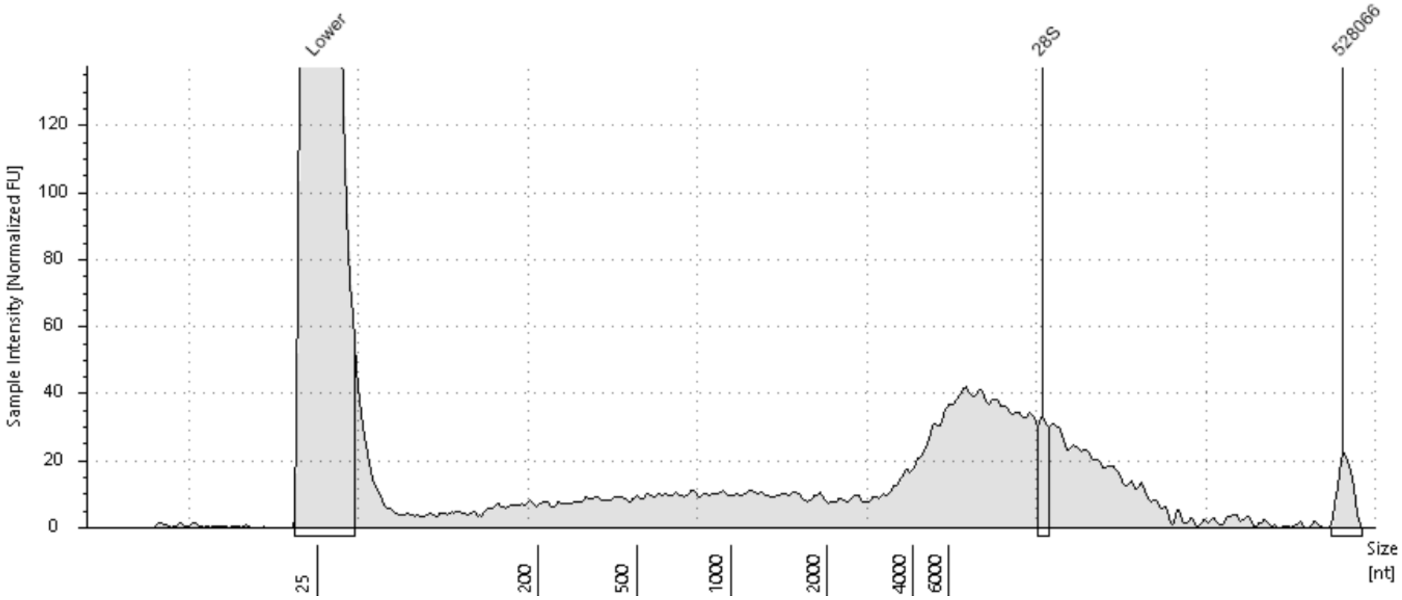


#### Sample Table

| Well | RIN$^e$ | 28S/18S (Area) | Conc. [pg/µl] | Sample Description | Alert | Observations |
|---|---|---|---|---|---|---|
| B1 | - | - | 135 | 22082R-05-13 | | |

#### Peak Table

| Size [nt] | Calibrated Conc. [pg/µl] | Assigned Conc. [pg/µl] | Peak Molarity [pmol/l] | % Integrated Area | Peak Comment | Observations |
|---|---|---|---|---|---|---|
| 25 | 700 | 700 | 82400 | - | | Lower Marker |
| 17500 | 4.07 | - | 0.685 | 49.71 | | 28S |
| 528066 | 4.12 | - | 0.0230 | 50.29 | | |

*C1: 22082R-05-14*

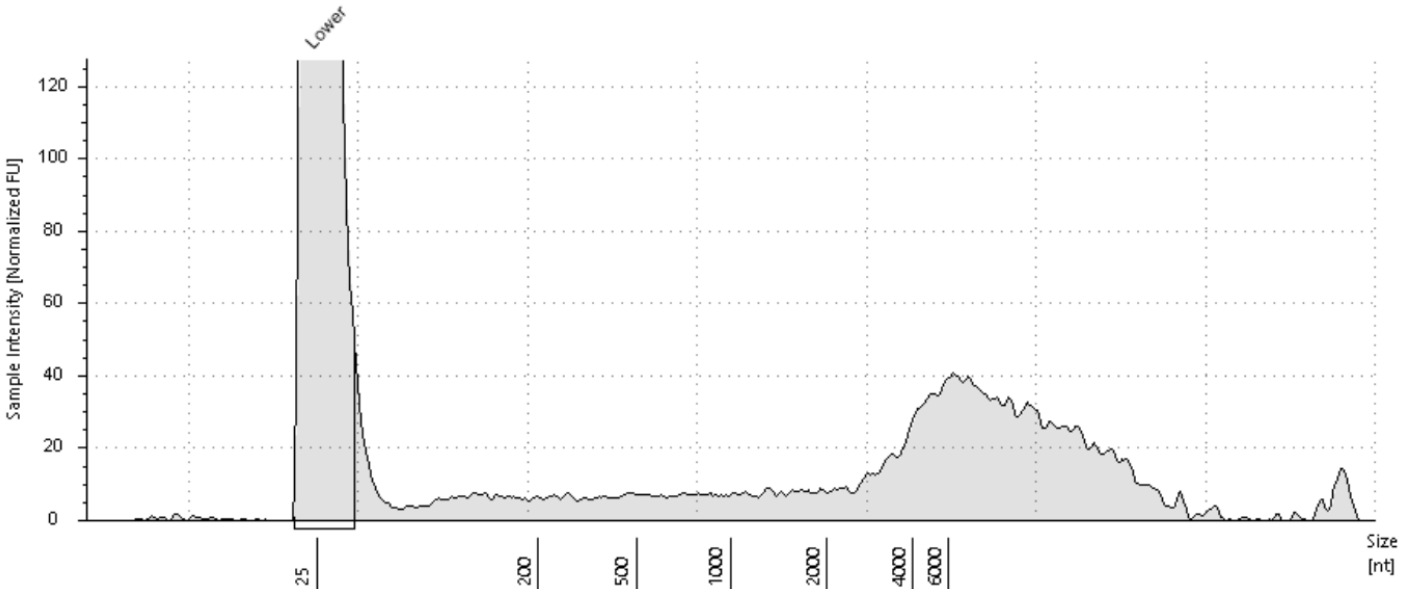


#### Sample Table

| Well | RIN[e] | 28S/18S (Area) | Conc. [pg/µl] | Sample Description | Alert | Observations |
|---|---|---|---|---|---|---|
| C1 | - | - | 129 | 22082R-05-14 | | |

#### Peak Table

| Size [nt] | Calibrated Conc. [pg/µl] | Assigned Conc. [pg/µl] | Peak Molarity [pmol/l] | % Integrated Area | Peak Comment | Observations |
|---|---|---|---|---|---|---|
| 25 | 700 | 700 | 82400 | - | | Lower Marker |

*D1: 22082R-05-15*

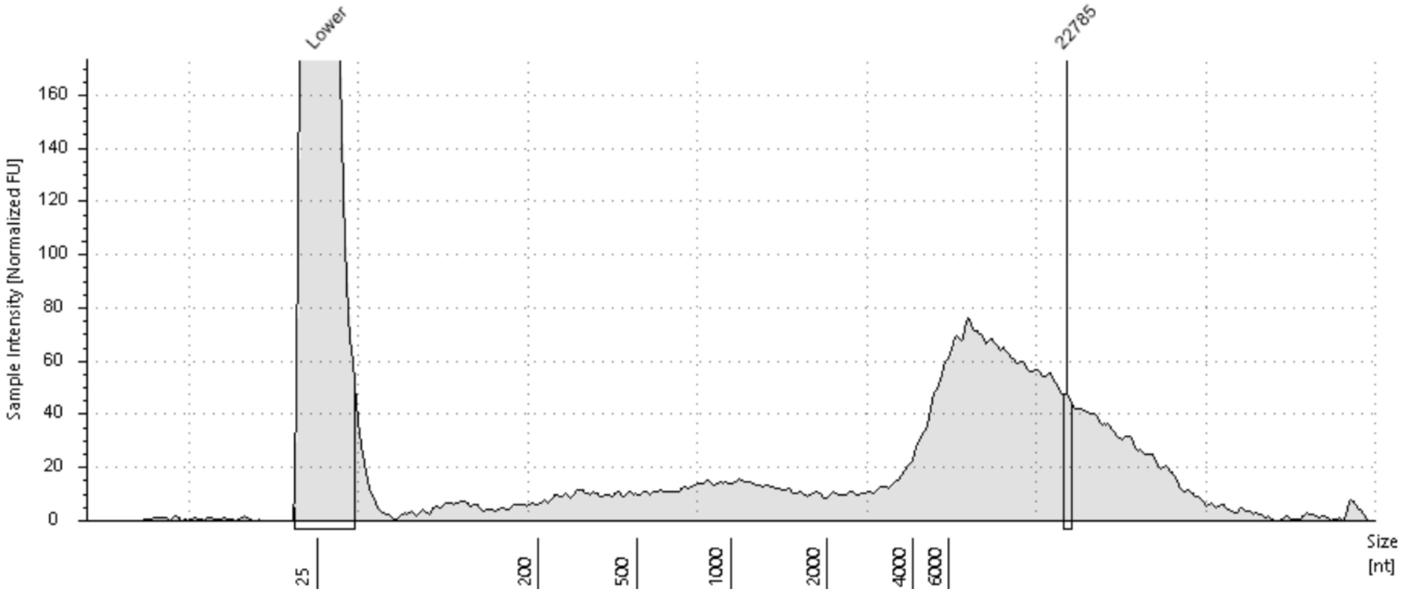


#### Sample Table

| Well | RIN$^{e}$ | 28S/18S (Area) | Conc. [pg/µl] | Sample Description | Alert | Observations |
|---|---|---|---|---|---|---|
| D1 | - | - | 198 | 22082R-05-15 | | |

#### Peak Table

| Size [nt] | Calibrated Conc. [pg/µl] | Assigned Conc. [pg/µl] | Peak Molarity [pmol/l] | % Integrated Area | Peak Comment | Observations |
|---|---|---|---|---|---|---|
| 25 | 700 | 700 | 82400 | - | | Lower Marker |
| 22785 | 4.74 | - | 0.612 | 100.00 | | |

*E1: 22082R-05-16*

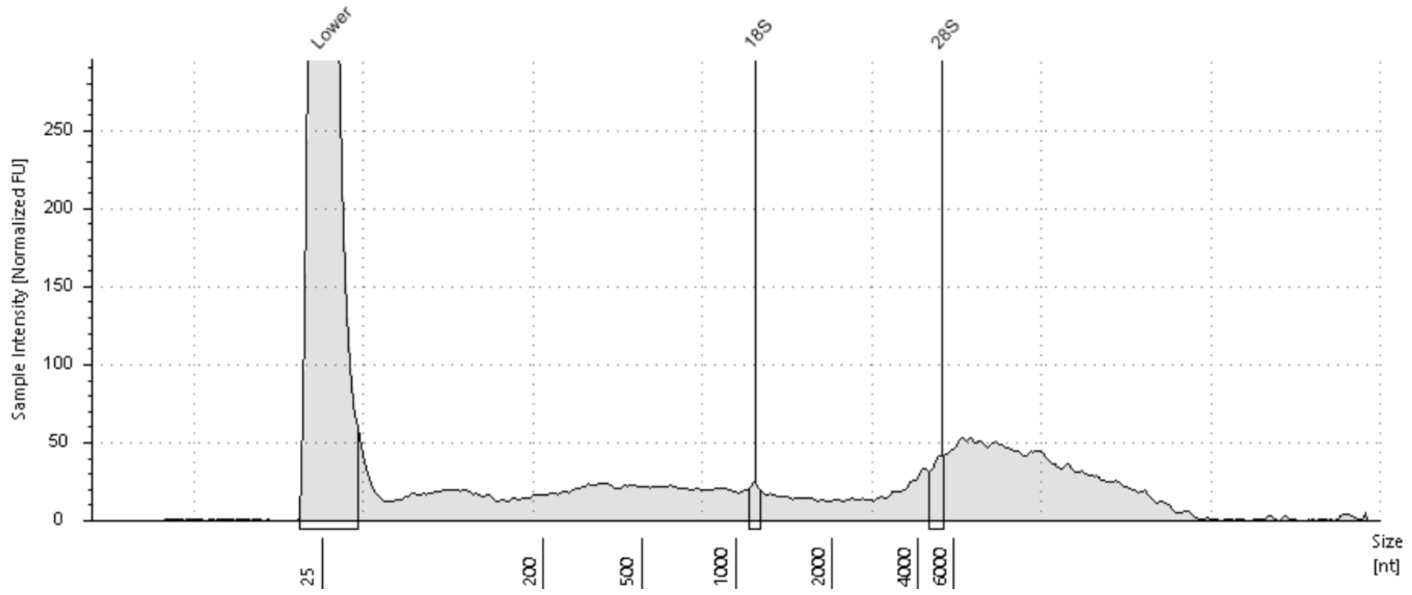


#### Sample Table

| Well | RIN[e] | 28S/18S (Area) | Conc. [pg/µl] | Sample Description | Alert | Observations |
|---|---|---|---|---|---|---|
| E1 | - | - | 212 | 22082R-05-16 | | |

#### Peak Table

| Size [nt] | Calibrated Conc. [pg/µl] | Assigned Conc. [pg/µl] | Peak Molarity [pmol/l] | % Integrated Area | Peak Comment | Observations |
|---|---|---|---|---|---|---|
| 25 | 700 | 700 | 82400 | - | | Lower Marker |
| 1142 | 3.09 | - | 7.95 | 31.19 | | 18S |
| 5228 | 6.81 | - | 3.83 | 68.81 | | 28S |

*F1: 22082R-05-17*

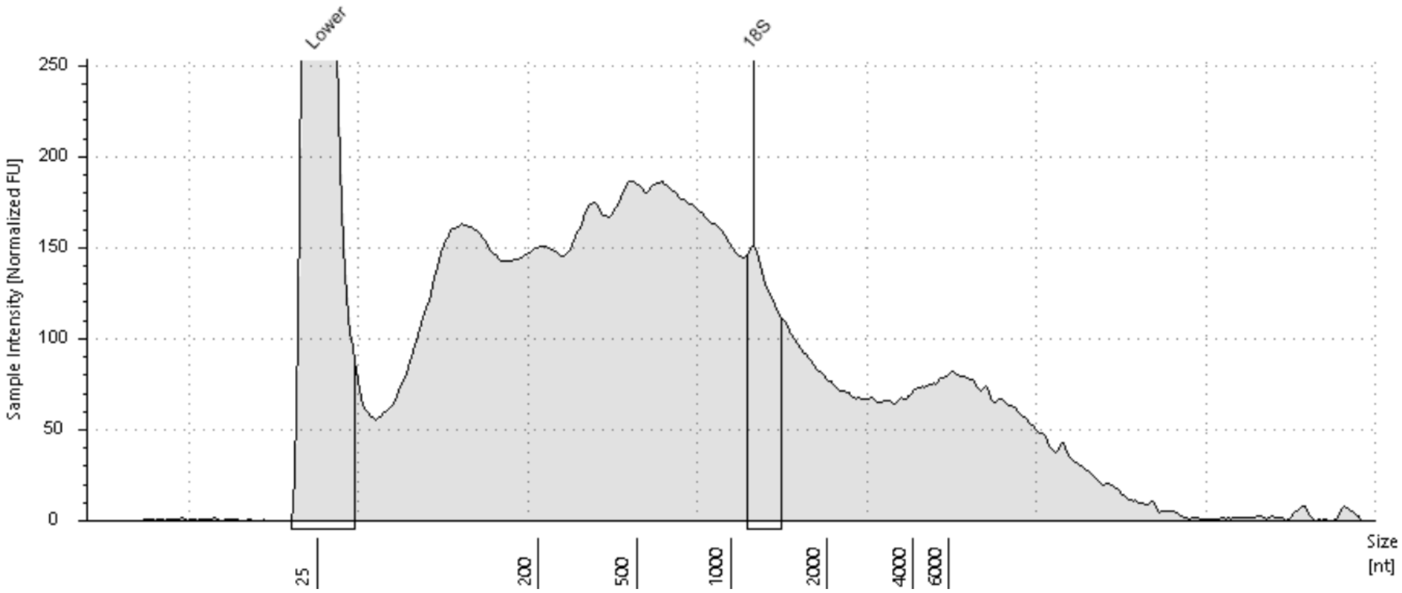


#### Sample Table

| Well | RIN$^e$ | 28S/18S (Area) | Conc. [pg/µl] | Sample Description | Alert | Observations |
|---|---|---|---|---|---|---|
| F1 | 2.0 | - | 906 | 22082R-05-17 | | |

#### Peak Table

| Size [nt] | Calibrated Conc. [pg/µl] | Assigned Conc. [pg/µl] | Peak Molarity [pmol/l] | % Integrated Area | Peak Comment | Observations |
|---|---|---|---|---|---|---|
| 25 | 700 | 700 | 82400 | - | | Lower Marker |
| 1176 | 50.3 | - | 126 | 100.00 | | 18S |

*G1: 22082R-05-18*

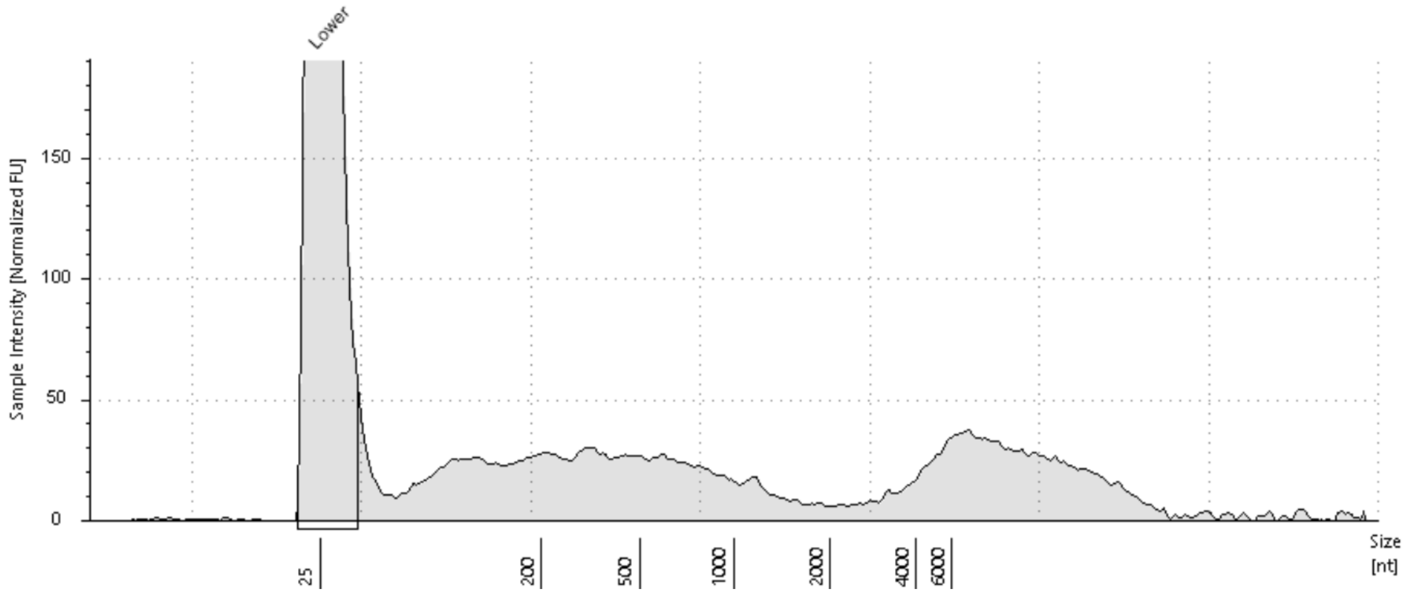


#### Sample Table

| Well | RIN^e | 28S/18S (Area) | Conc. [pg/µl] | Sample Description | Alert | Observations |
|---|---|---|---|---|---|---|
| G1 | - | - | 179 | 22082R-05-18 | | |

#### Peak Table

| Size [nt] | Calibrated Conc. [pg/µl] | Assigned Conc. [pg/µl] | Peak Molarity [pmol/l] | % Integrated Area | Peak Comment | Observations |
|---|---|---|---|---|---|---|
| 25 | 700 | 700 | 82400 | - | | Lower Marker |

## *H1: 22082R-05-19*

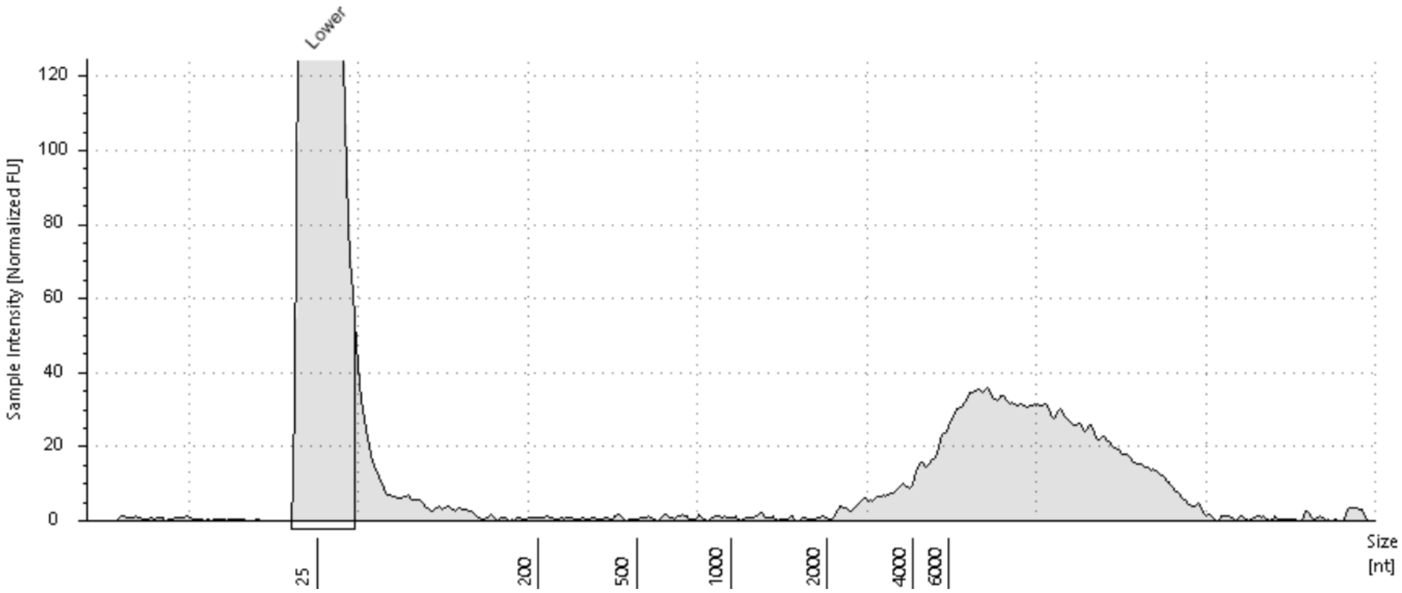


##### Sample Table

| Well | RIN[e] | 28S/18S (Area) | Conc. [pg/µl] | Sample Description | Alert | Observations |
|---|---|---|---|---|---|---|
| H1 | - | - | 87.4 | 22082R-05-19 | | |

##### Peak Table

| Size [nt] | Calibrated Conc. [pg/µl] | Assigned Conc. [pg/µl] | Peak Molarity [pmol/l] | % Integrated Area | Peak Comment | Observations |
|---|---|---|---|---|---|---|
| 25 | 700 | 700 | 82400 | - | | Lower Marker |

*A2: 22082R-05-20*

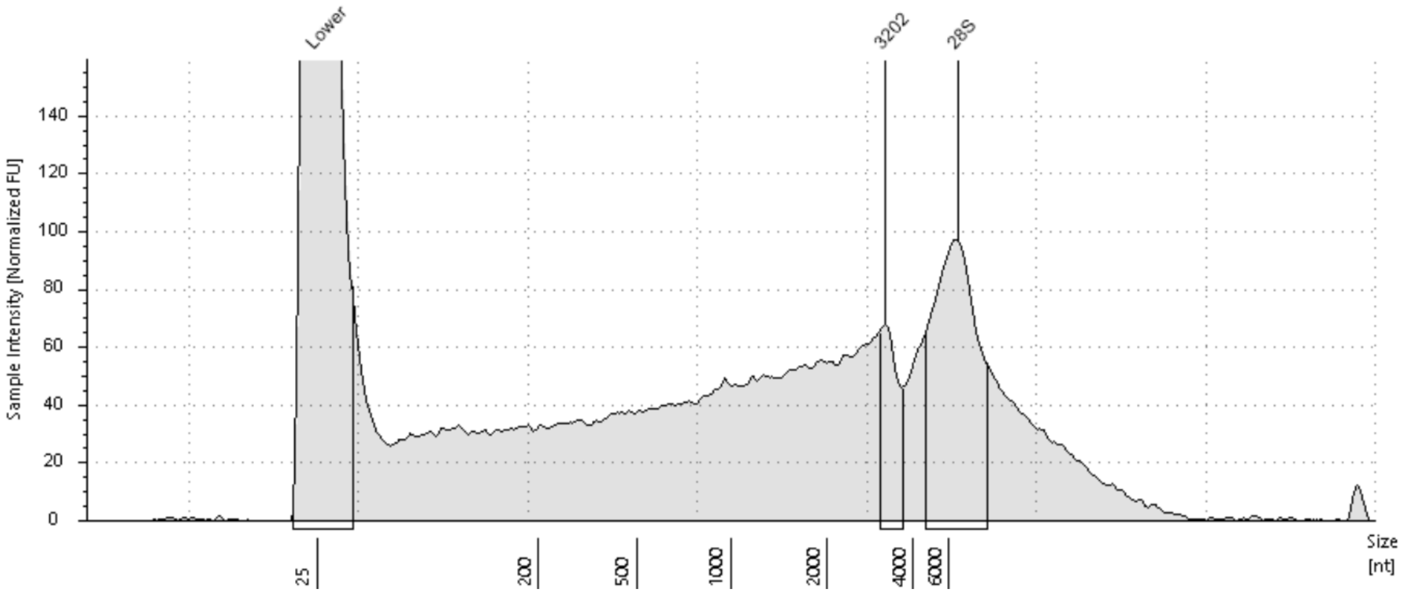


#### Sample Table

| Well | RIN$^e$ | 28S/18S (Area) | Conc. [pg/µl] | Sample Description | Alert | Observations |
|---|---|---|---|---|---|---|
| A2 | - | - | 373 | 22082R-05-20 | | |

#### Peak Table

| Size [nt] | Calibrated Conc. [pg/µl] | Assigned Conc. [pg/µl] | Peak Molarity [pmol/l] | % Integrated Area | Peak Comment | Observations |
|---|---|---|---|---|---|---|
| 25 | 700 | 700 | 82400 | - | | Lower Marker |
| 3202 | 16.0 | - | 14.7 | 22.12 | | |
| 6696 | 56.5 | - | 24.8 | 77.88 | | 28S |

### *B2: 22082R-05-21*

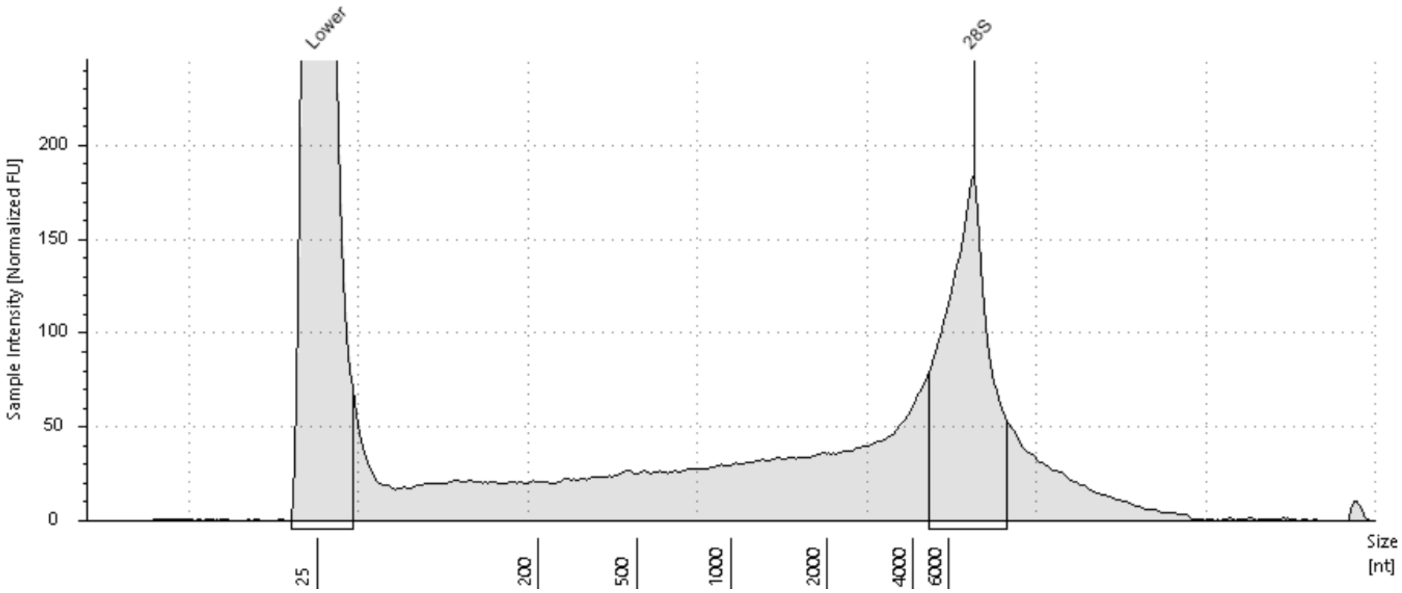


#### Sample Table

| Well | RIN$^e$ | 28S/18S (Area) | Conc. [pg/µl] | Sample Description | Alert | Observations |
|---|---|---|---|---|---|---|
| B2 | - | - | 321 | 22082R-05-21 | | |

#### Peak Table

| Size [nt] | Calibrated Conc. [pg/µl] | Assigned Conc. [pg/µl] | Peak Molarity [pmol/l] | % Integrated Area | Peak Comment | Observations |
|---|---|---|---|---|---|---|
| 25 | 700 | 700 | 82400 | - | | Lower Marker |
| 8023 | 101 | - | 37.0 | 100.00 | | 28S |

*C2: 22082R-05-22*

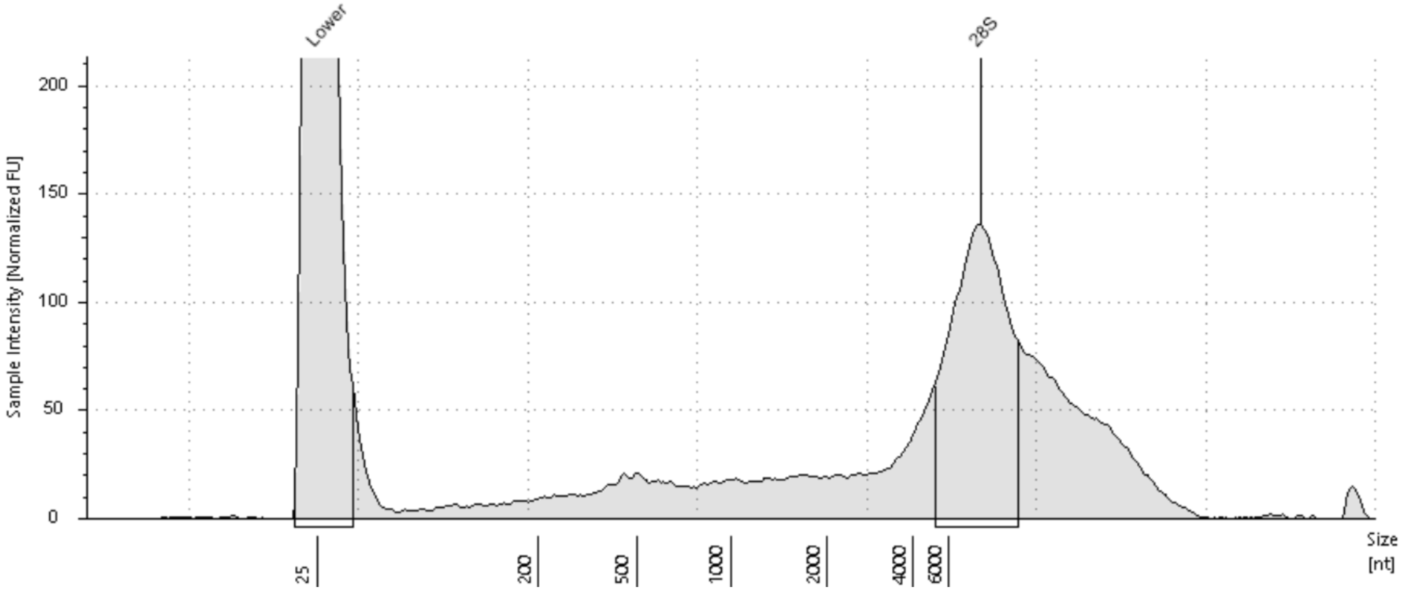


#### Sample Table

| Well | RINe | 28S/18S (Area) | Conc. [pg/µl] | Sample Description | Alert | Observations |
|---|---|---|---|---|---|---|
| C2 | - | - | 282 | 22082R-05-22 | | |

#### Peak Table

| Size [nt] | Calibrated Conc. [pg/µl] | Assigned Conc. [pg/µl] | Peak Molarity [pmol/l] | % Integrated Area | Peak Comment | Observations |
|---|---|---|---|---|---|---|
| 25 | 700 | 700 | 82400 | - | | Lower Marker |
| 8525 | 100 | - | 34.6 | 100.00 | | 28S |

*D2: 22082R-05-23*

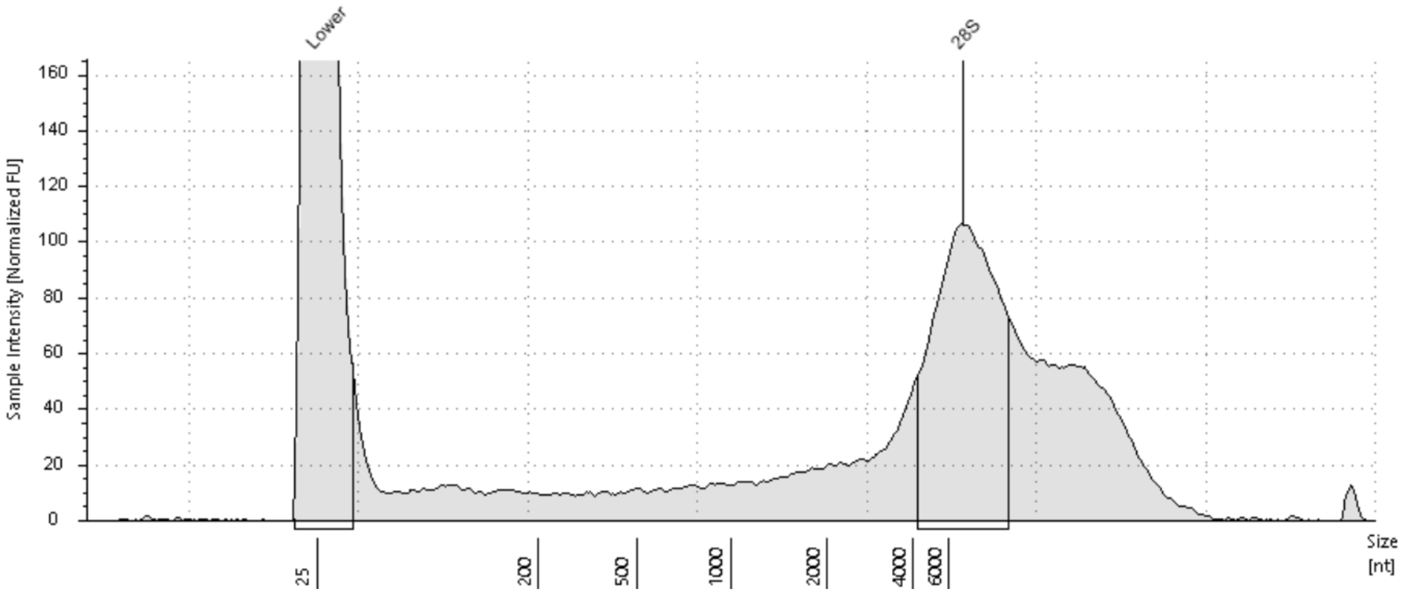


#### Sample Table

| Well | RIN[e] | 28S/18S (Area) | Conc. [pg/µl] | Sample Description | Alert | Observations |
|---|---|---|---|---|---|---|
| D2 | - | - | 281 | 22082R-05-23 | | |

#### Peak Table

| Size [nt] | Calibrated Conc. [pg/µl] | Assigned Conc. [pg/µl] | Peak Molarity [pmol/l] | % Integrated Area | Peak Comment | Observations |
|---|---|---|---|---|---|---|
| 25 | 700 | 700 | 82400 | - | | Lower Marker |
| 7010 | 94.9 | - | 39.8 | 100.00 | | 28S |

*E2: 22082R-05-24*

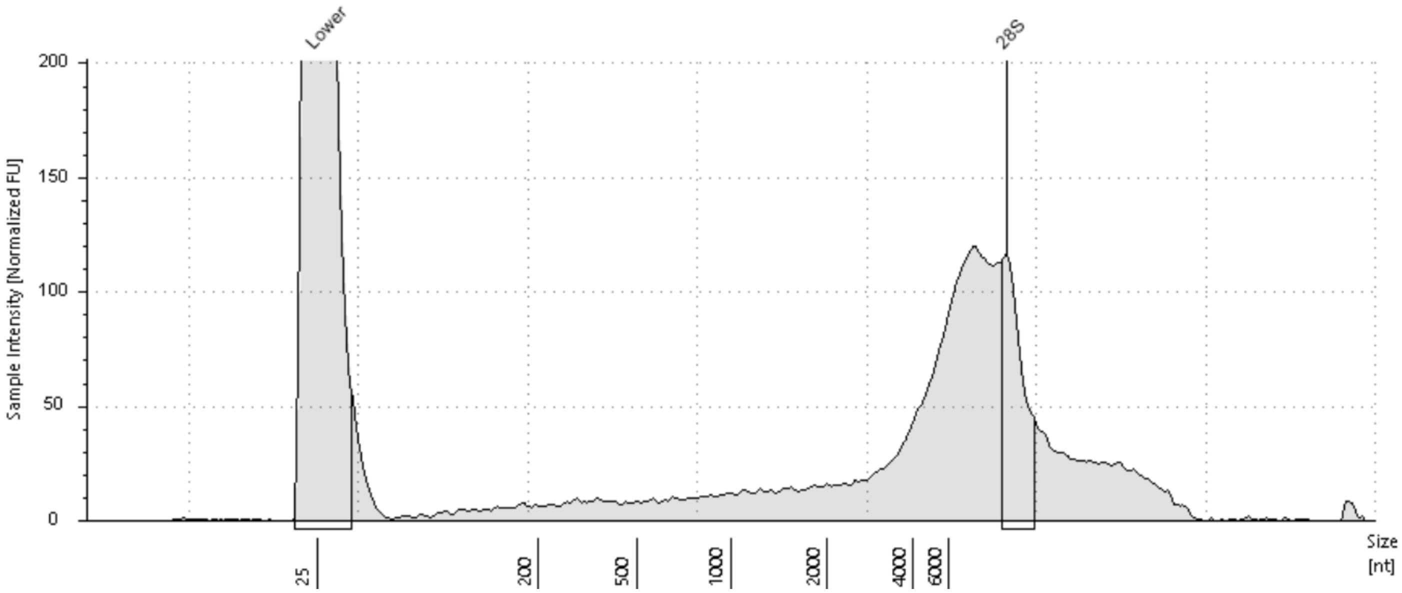


#### Sample Table

| Well | RIN[e] | 28S/18S (Area) | Conc. [pg/µl] | Sample Description | Alert | Observations |
|---|---|---|---|---|---|---|
| E2 | - | - | 238 | 22082R-05-24 | | |

#### Peak Table

| Size [nt] | Calibrated Conc. [pg/µl] | Assigned Conc. [pg/µl] | Peak Molarity [pmol/l] | % Integrated Area | Peak Comment | Observations |
|---|---|---|---|---|---|---|
| 25 | 700 | 700 | 82400 | - | | Lower Marker |
| 11672 | 34.1 | - | 8.59 | 100.00 | | 28S |
